# RNA Alternative Splicing and Polyadenylation and Regulation of the Glomerular Filtration Barrier

**DOI:** 10.1101/2024.09.26.615280

**Authors:** Monoj K Das, Amy Webb, Mahika Yarram, Christian Reilly, Lalith Punepalle, Claire Bryant, Rajgopal Govindarajan, Claire L. Moore, Shipra Agrawal

## Abstract

**Background:** Glomerular disease, characterized by podocyte injury and proteinuria, can lead to chronic kidney disease and end stage kidney disease. We hypothesized that the glomerular pathophysiology is associated with mRNA alternative splicing and polyadenylation of glomerular genes and of critical podocyte and slit diaphragm components that regulate the filtration barrier.

**Methods:** Glomerular damage, accompanied by proteinuria, was induced by puromycin-aminonucleoside or adriamycin to mimic human minimal change disease or focal segmental glomerulosclerosis (FSGS), respectively, and RNA-seq analyses was performed. Alternatively spliced and alternatively polyadenylated events through differential exon and poly(A) site usage were queried in JunctionSeq and APATrap pipelines. These events were further mapped on podocyte and glomerular landscape, analyzed and modulated for slit diaphragm components, and cis- and trans-regulatory elements were identified.

**Results:** Altered glomerular mRNA processing by alternative splicing/polyadenylation was identified in 136/71 and 1875/746 genes in minimal change disease and FSGS models, respectively. Transcript annotation and prioritization of significant alternative splicing and polyadenylation identified key events in several podocyte and slit diaphragm genes with novel and established roles. Alternative splicing of critical slit diaphragm components, the junction protein TJP1/ZO1 and microtubule associating protein ITM2B was further characterized. Alternative polyadenylation of core members of the slit diaphragm, *NPHS1*, *NPHS2* and NEPH1 was analyzed with potential alteration of microRNA binding sites between the proximal vs distal poly (A) site usage in their mRNAs. Concomitantly, dysregulation of trans-regulatory elements (polyadenylation and splicing factors), was discovered in these models of nephropathies. Additionally, beneficial proteinuria-reducing treatments, pioglitazone and GQ16 reversed many alternatively spliced and polyadenylated events. Moreover, GWAS SNPs as potential cis-regulatory elements were identified in several genes from the human nephrotic syndrome database. Finally, we demonstrated proof-of-concept principle of chemically modified splice switching oligonucleotides in modulating *TJP1* alternative splicing in podocytes.

**Conclusion:** Findings from our studies identified that glomerular pathophysiology and disruption of the filtration barrier is associated with alternative splicing and alternative polyadenylation of glomerular genes, many of which are crucial determinants of podocyte structure and function and the slit diaphragm complex.

**Key Points:**

- Alternative mRNA processing adds an important layer of transcriptome complexity in podocyte and slit diaphragm components during glomerular injury.
- Trans- and cis-elements may offer potential mechanisms to regulate alternative splicing and polyadenylation during podocyte injury.
- Antisense oligonucleotides can regulate mRNA processing events, thus offering a potential RNA-based therapeutic strategy to treat glomerular disease.

## Introduction

Glomerular disease is characterized by podocyte injury and loss, and manifests with high-grade proteinuria and additional co-morbidities^1–6^. Various forms of glomerular disease can unfortunately be refractory to existing treatments, and progress to chronic kidney disease (CKD) and end stage kidney disease^7–12^. Thus, there is a constant unmet clinical need to develop targeted therapies for glomerular disease, fueled by basic molecular studies.

Research has unraveled remarkable complexities of mRNA processing and its roles in normal cellular function and disease^13, 14^. ∼95% of mammalian genes undergo alternative splicing^15^, which can result in different proteins with distinct functionality. Alternative splicing also determines mRNA stability, localization, translation efficiency and micro-RNA sensitivity^16, 17^. Moreover, we and others have shown that alternative splicing is associated with many cellular processes and diseases such as differentiation, cancer, and immunity^16, 18–22^. While, there are a few reports of aberrant alternative splicing in a handful of genes relevant to CKD and glomerular disease^23^, including our work on PPARγ^24^, the mechanisms and role of alternative splicing in glomerulopathies is lacking.

mRNA polyadenylation is an essential maturation step in which precursor-mRNA is cleaved and trimmed at its 3’-untranslated-region (UTR) and a poly(A)-tail is added^25^. Change in the position of the poly(A) site through a process called alternative polyadenylation plays an important, increasingly appreciated role in regulation of gene expression, cellular functions and pathological processes^26–34^. Alternative polyadenylation occurs in ∼70% of genes^25, 35^, and shortening of the 3’ UTR can remove regulatory sequences that control RNA stability, translation, and subcellular localization^26, 27, 29, 32, 33^, whereas coding region shortening can dramatically alter protein function^28, 30, 31, 34^. While global changes in alternative polyadenylation have been observed in development and tumor progression, there are no reports of alternative polyadenylation in glomerulopathies.

We hypothesized that glomerular disease pathophysiology is interlinked with alternative processing of podocyte and glomerular mRNAs of key genes that regulate the filtration barrier. To study these events, we employed animal models of glomerular disease, JunctionSeq and APATrap pipelines, differentiated human podocytes and the GWAS glomerular disease database.

## Methods

### Animal Models

The animal studies were conducted under the approval and guidelines of the Institution Animal Care and Use Committee at Nationwide Children’s Hospital. Briefly, male Wistar rats were injected intravenously (IV) with puromycin aminonucleoside (PAN, 50mg/kg) or adriamycin (7.5mg/kg) to model minimal change disease or focal segmental glomerulosclerosis (FSGS), respectively, and euthanized on Day 11 and 21 respectively, at which time kidneys were harvested and glomeruli isolated using the sequential sieving method and total RNA was isolated from the glomeruli. Kidneys were paraffin embedded and processed for histology. Serum and urine chemistry were performed on serial samples. Details are provided in the Supplementary Methods.

Additional details regarding methods and procedures, including RNASeq and Differential Expression of Genes (DEGs), JunctionSeq^36^, rMATS-turbo^37^ and APATrap Analyses^38^, Heatmaps, Pathway Analysis and Ontology Enrichment, GWAS Correlation with alternatively spliced and alternatively polyadenylated genes, Reverse Transcriptase-Polymerase Chain Reaction (RT-PCR), 3’ Rapid Amplification of cDNA Ends (RACE) Assay, Splice-Switching Oligonucleotide (SSO) Design and Assay, Western Blotting and Immunofluorescence Microscopy, and Podocyte Culture and Treatments are provided in the Supplementary Data, and in **Table 1**.

**Table 1:**
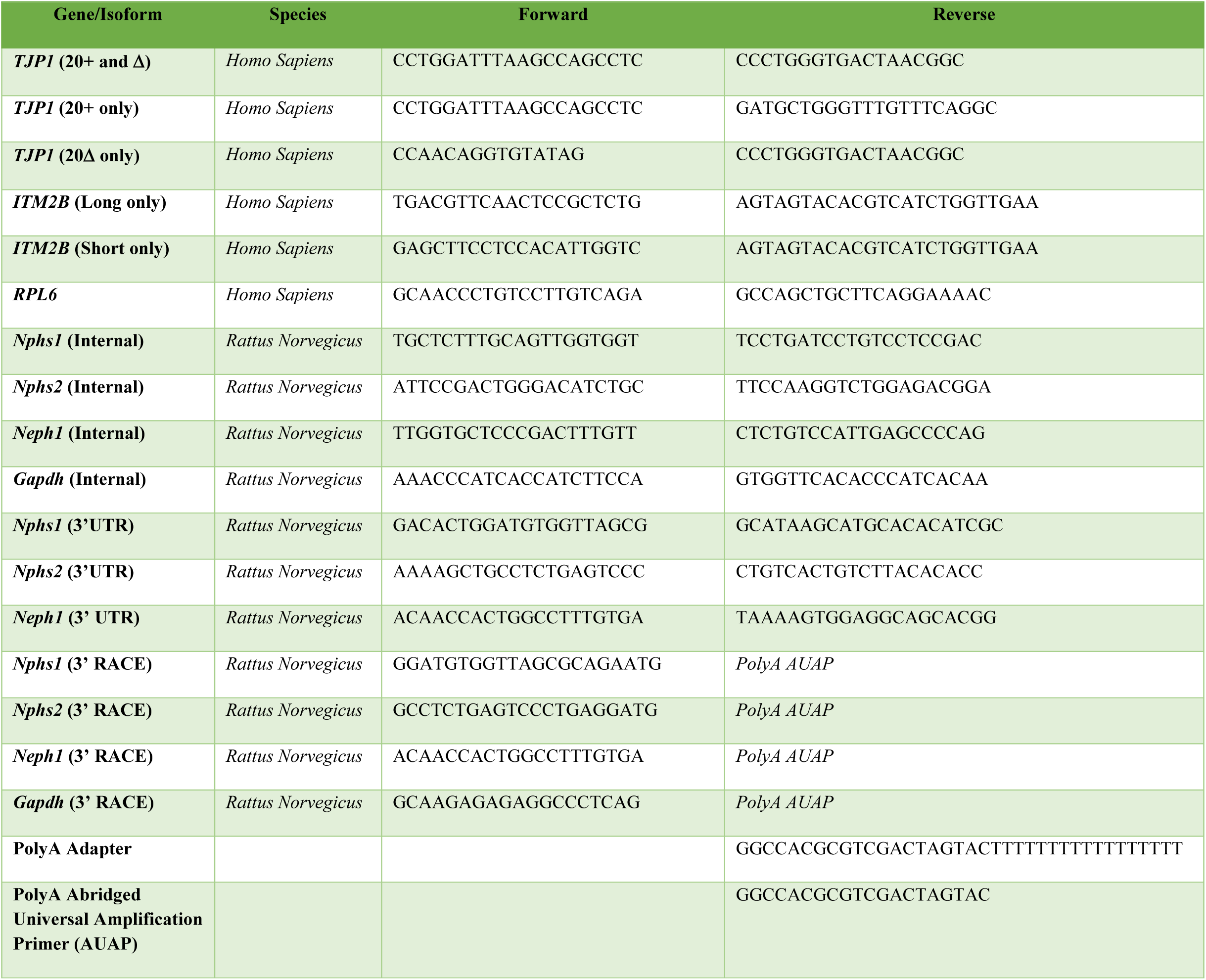
Primers (5’ to 3’) used in the study.

### Statistics

Animal experiments were performed with n=4/group to achieve sufficient power (80%) and downstream analyses was performed after confirmation of significant proteinuria induction in both PAN and adriamycin groups. Statistical analyses were performed and adjusted p values used as described in Supplemental Data for RNASeq DEG, JunctionSeq, rMATS, APATrap data and GWAS SNPs threshold. Differentially expressed genes (DEGs) were chosen with adjusted p<0.05 and abs(logFC)>1<1, unless otherwise noted. All other experimental data were obtained from at least three independent experiments and expressed as mean±SEM. For UPCR, serum chemistry, qRT-PCR and densitometry analysis, one-way ANOVA or t test was used and p<0.05 was considered as significant.

## Results

### I. PAN and Adriamycin-Nephropathy Models Revealed Common and Distinct Transcriptome Signatures

#### Glomerular and Podocyte Injury in PAN and ADR Nephropathy Models

To study the alternative processing of glomerular mRNAs during disease, we used two well-established rat models of podocyte and glomerular injury; PAN-induced minimal change disease and adriamycin-induced FSGS) (**Figure 1A**). Proteinuria induced by IV PAN injection appeared by Day 4 and massive proteinuria was observed in adriamycin-injected rats, starting from Day 7 (**Figures 1B and C**). Histology revealed podocyte hypertrophy in the adriamycin-model and multifocal dilation of proximal tubules in both PAN and adriamycin-models (**Figure 1D)**, along with disease-related changes in serum chemistry (**Supplementary Figure S1)**.

**Figure 1.**
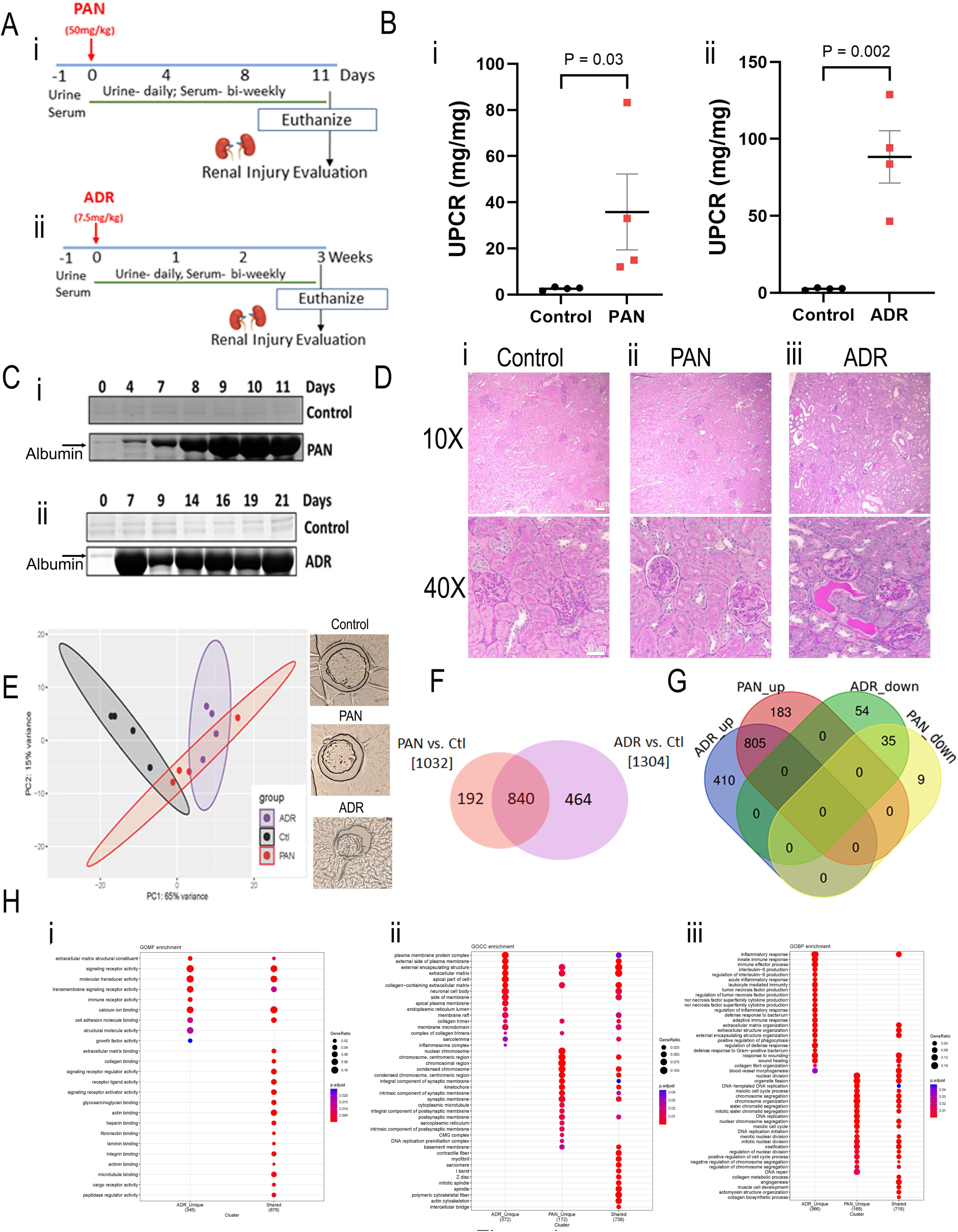
Puromycin amino-nucleoside (PAN)- and adriamycin (ADR)-induced models of nephropathy revealed common and distinct transcriptome signatures in glomerulopathies. **(A)** Schematic for PAN and adriamycin (ADR)-induced nephropathy models showing a single IV injection of **(Ai)** PAN (50 mg/kg) and **(Aii)** adriamycin (ADR) (7.5 mg/kg) in male Wistar rats resulted in **(B)** induction of proteinuria and increased urinary protein/creatinine ratio (UPCR) compared to healthy controls. PAN: 35.83 mg/mg ± SEM 16.47 mg/mg, Day 11; Adriamycin (ADR): 88.31 mg/mg ± SEM 16.95 mg/mg, Day 21. **(C)** Representative urine gels showing temporal increase in albumin bands in both PAN and adriamycin (ADR) models. **(D)** Periodic acid schiff (PAS) stained images (10X and 40X) of the kidney cortex of a representative rat from each group: **(Di)** control, **(Dii)** PAN-induced nephropathy, and **(Diii)** adriamycin (ADR)-induced nephropathy. Normal kidney parenchyma was observed in healthy controls. In contrast, PAN- and adriamycin (ADR)-injured rats displayed multifocal dilation of proximal tubules due to the presence of intraluminal protein casts, with a much higher increase with adriamycin (ADR) injury. At the observed timepoints there were no visible histologic glomerular changes in the PAN model (Day 11) In the adriamycin (ADR) rats (Day 21), many of the glomeruli showed marked hypertrophy of podocytes with prominent intracytoplasmic vacuolation. At least 10 glomerular fields were observed per rat (n=4) per group. Scale bar 50 µm, 40X images and 100 µm, 10X images. **(E)** RNA Seq followed by Principal Component analysis (PCA) depicting a glomerular transcriptomic shift in PAN and adriamycin (ADR) groups compared to controls. Each dot represents the glomerular transcriptome of a healthy and PAN and adriamycin (ADR)-injured rat. Integrity of the glomeruli for each group is shown by representative glomerulus embedded in Histogel on the right. Scale bar 10 µm. **(F)** Venn diagram of differentially expressed genes (DEGs) observed in the glomeruli of PAN and adriamycin (ADR) models compared to healthy controls. 840 DEGs are common, 192 DEGs unique to PAN model, and 464 DEGs unique to adriamycin (ADR) model. **(G)** Common and distinct upregulated and downregulated DEGs in PAN and adriamycin (ADR) disease models. 805 genes are upregulated and 35 genes are downregulated in both PAN and adriamycin (ADR). 183 and 410 genes were distinctly upregulated, and 9 and 54 genes were distinctly downregulated in the PAN and adriamycin (ADR) models, respectively. **(H)** Ontology enrichment analysis identified the top cellular components (GOCC, **Hi**), biological processes (GOBP, **Hii**), and molecular functions (GOMF, **Hiii**) associated with each disease model for their unique DEGs and shared DEGs. (**Hi**) The cellular components of DEGs unique to adriamycin (ADR) included extracellular matrix and collagen trimer, those unique to PAN included basement membrane and synaptic membrane and of DEGs shared between the two models included actin cytoskeleton and intercellular bridges. (**Hii**) The biological processes identified from the DEGs unique to adriamycin (ADR) included extracellular matrix and collagen fibril organization and IL6 and TNFα production, those unique to PAN included regulation of nuclear division and chromosome segregation and of DEGs shared between the two models included extracellular matrix organization and collagen biosynthesis. (**Hiii**) The molecular functions identified from the DEGs unique to adriamycin (ADR) included extracellular matrix binding and of DEGs shared between the two models included actin, integrin and microtubule binding.

#### RNA-Seq Analyses Revealed Unique and Shared Changes in Transcriptomic Landscape and Pathways in PAN and ADR Models

To gain insights into the genes and molecular pathways, we performed deep paired RNA Seq analysis from glomeruli isolated by sequential sieving method^39^ from healthy and PAN and adriamycin-injured rats (Day 11 and 21, respectively). Distinct transcriptomic shifts were observed in PAN and adriamycin groups compared to controls (**Figure 1E**), and a total of 1,032 DEGs in PAN vs Control (988↑, 44↓) and 1,304 DEGs in adriamycin vs Control (1215↑, 89↓) (adj *p*<0.05 and log2FC>1<1) (**Figure 1F**). Further analysis identified common and distinct DEGs for each model (**Figure 1G**). Ontology enrichment analysis identified the top cellular components (**Figure 1Hi**), biological processes (**Figure 1Hii**), and molecular functions (**Figure 1Hiii**) for each model with unique and shared DEGs. These included extracellular matrix, actin cytoskeleton, intercellular bridges, cytokine synthesis, and components of synaptic membrane. These were further categorized by overall and ⇅ DEGs in PAN and adriamycin models (**Supplementary Figure S2**).

Deconvolution of the transcriptome identified the contribution towards pathogenesis from a large number of genes specific to podocytes as well as other glomerular cell types, including mesangial, endothelial cells and parietal epithelial cells (**Supplementary Figure S3, Table S1**). Moreover, we identified DEGs in several established causal genes with known association with glomerular disease and podocytopathies, as well as novel genes without prior association in the transcriptome of PAN and adriamycin nephropathy models (**Supplementary Figure S4**).

### II. mRNA Alternative Splicing Occurred During Glomerular Disease Pathogenesis

#### Glomerular mRNAs Underwent Alternative Splicing during Glomerular Injury

We used the JunctionSeq program^36^ to analyze glomerular RNASeq dataset to identify differential exon usage in the PAN and adriamycin models. Glomerular injury identified a total of 136 (PAN) and 1875 (adriamycin) alternatively spliced glomerular genes vs. controls (adj *p*<0.05) (**Figure 2A,B; Supplementary Figure S5**). Among these, 104 genes were shared, while 32 and 1771 were unique to the PAN and adriamycin model, respectively. Distribution of alternative splicing events in PAN and adriamycin groups was detected using rMATS-turbo^37^, with the skipped exon identified as the most prevalent type of event (**Figure 2C)**.

**Figure 2.**
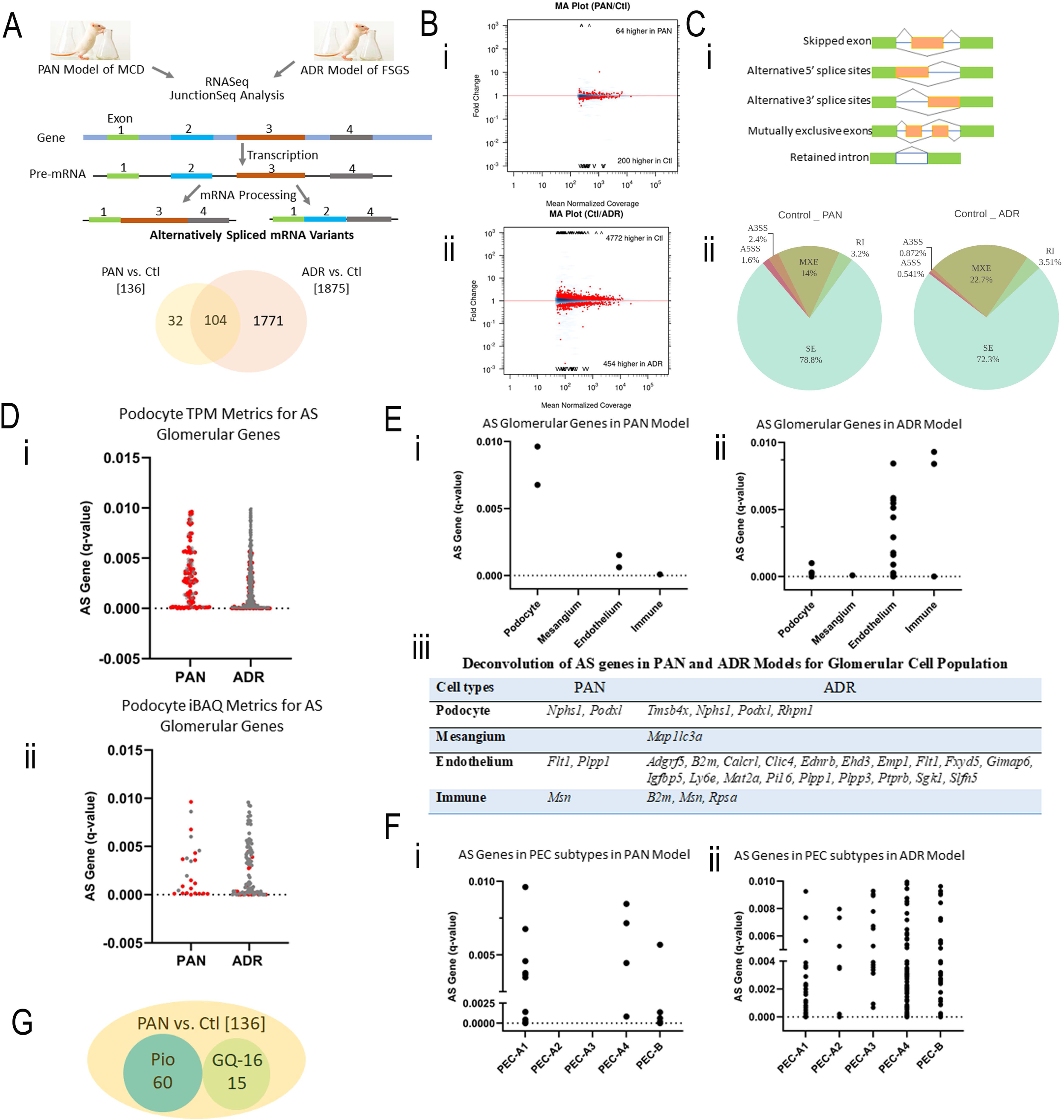
Glomerular mRNAs were alternatively spliced (AS) in puromycin amino-nucleoside (PAN)- and adriamycin (ADR)-induced nephropathy models mimicking minimal change disease (MCD) and FSGS. **(A)** Schematic showing the experimental and JunctionSeq bioinformatic workflow to detect differential exon usage or alternatively spliced mRNA transcripts (shown as mutually exclusive exons) in the glomeruli of rats subjected to PAN- and adriamycin (ADR)-induced injury. Alternatively spliced transcripts can also be generated due to additional splicing events (such as alternative 5’ and 3’ splice sites usage, intron retention, exon skipping/inclusion) not depicted in the schematic. Venn diagram depicts the common and distinct alternatively spliced genes detected in both the models compared to healthy rats. **(B)** MA plots depict the relationship between normalized coverage and fold change between groups highlighting the measures that are much higher in one group than the other, observed in **(Bi)** PAN and **(Bii)** adriamycin (ADR) models. **(Ci)** Different kinds of alternative splicing events, and **(Cii)** their Distribution in PAN and Adriamycin (ADR). Pie charts illustrate the proportion of five alternative splicing event types: Skipped exon (SE), Retained intron (RI), Alternative 5’ splice sites (A5SS), Alternative 3’ splice sites (A3SS) and Mutually exclusive exons (MXE) detected across PAN and adriamycin (ADR) injury compared to healthy control using rMATS-turbo. The distribution highlights differences in splicing patterns, with statistically significant events selected based on an FDR threshold (≤0.05). **(D)** Podocyte metrics. **(Ci)** Podocyte-specific transcriptomic dataset of 6593 genes (TPM > 10, *q* < 0.05), identified 115 and 1342 alternatively spliced genes (*q* < 0.05) in PAN and adriamycin (ADR) models. 90 common alternatively spliced genes between the two models are in red. **(Dii)** Podocyte-specific iBAQ dataset of 551 proteins (*q* < 0.05) identified 24 and 96 alternatively spliced genes (*q* < 0.05) in PAN and adriamycin (ADR) models. 17 common alternatively spliced genes between the two models are in red. **(E)** Deconvolution distribution of alternatively spliced genes across the different glomerular cell types, including podocytes, mesangial cells, endothelial cells and a small fraction of immune cells in **(Ei)** PAN and **(Eii)** adriamycin (ADR) models. **(Eiii)** Genes from the deconvolution represented in PAN and adriamycin (ADR) models. **(F)** Deconvolution distribution of alternatively spliced genes across the different parietal epithelial cell (PEC) subtypes, including PEC-A1, PEC-A2, PEC-A3, PEC-A4 and PEC-B population, in **(Fi)** PAN and **(Fii)** adriamycin (ADR) models. **(G)** Venn Diagram showing alternatively spliced genes reversed with Pio and GQ-16 treatments in PAN model.

#### Robust Podocyte Metrics and Contribution from Other Glomerular Cells Identified for Alternative Splicing Events

PAN-minimal change disease and adriamycin-FSGS nephropathies are mainly characterized by podocyte injury, although other glomerular cells contribute towards pathogenesis. We deconvoluted the JunctionSeq-derived dataset by cross-referencing the significant alternative splicing with podocyte-specific transcriptomic and proteomic dataset^40^, and with the glomerular top 50 most variable cell-type specific genes^41^. From the podocyte-specific transcriptomic dataset (6593 genes; TPM>10, *q*<0.05)^40^, we identified 115 (PAN), 1342 (adriamycin), and 90 (PAN and adriamycin) alternatively spliced genes (*q*<0.05) (**Figure 2Di**). From the podocyte-specific iBAQ dataset (551 proteins; *q*<0.05)^40^, we identified 24 (PAN), 96 (adriamycin), and 17 (PAN and adriamycin) alternatively spliced genes (*q*<0.05) (**Figure 2Dii**). <u>Podocyte</u> specific genes *Nphs1, Podxl1* were alternatively spliced in both disease models, while adriamycin-model presented with additional alternative splicing events in *Tmsb4x* and *Rhpn1* (**Figure 2Ei-iii**). <u>Mesangial</u> specific gene *Map1lc3a* was alternatively spliced in adriamycin, and a few endothelial genes were alternatively spliced in either both the models (i.e., *Flt1*), or uniquely in the adriamycin-model (i.e*. Sgk1, Adgrf5, Ehd3*). <u>Immune</u> <u>cell</u> specific gene, *Msn*, was alternatively spliced in both disease models, however, *Rpsa and B2m w*ere uniquely alternatively spliced in adriamycin. To address the diverse roles of parietal epithelial sub-types in the podocyte-regeneration and glomerular pathophysiology, we deconvoluted the significant alternative splicing events with parietal epithelial sub-populations^42^. We identified alternative splicing events in all five parietal epithelial sub-populations, with the most events clustered in the podocyte-progenitor A1 sub-population and surprisingly in the tubular-progenitors A4 sub-population, especially in the adriamycin-model (**Figure 2Fi and ii, Supplementary TableS2**).

#### Glomerular mRNAs Underwent Alternative Splicing in Genes with Novel and Established Roles in Glomerulopathies

In order to correlate the identified alternative splicing events in glomerular disease with genes that have established causal roles in human patients, we cross-referenced the alternatively spliced dataset with 80 monogenic glomerular disease genes^43^ and 49 podocytopathy genes associated with steroid resistant nephrotic syndrome ^44^. We identified 8 (PAN) and 19 (adriamycin) alternatively splicing genes with established causal roles in glomerular disease, which mostly localized within the podocytes, endothelium or glomerular basement membrane (GBM) (**Table 2a)**. Among these, 7 were common, and 14 only found in the adriamycin-model. The common genes included those localized to the podocyte membrane (*Podxl*) and cytoskeletal scaffold (*Tns2, Inf2, Itsn2*) and matrix genes in the GBM (e.g. *Lamb2* and *Col4a4*). The adriamycin-model exhibited distinct genes in the GBM (*Col4a3/5*) and in the slit diaphragm (*Nphs1*). Furthermore, comparison with genes associated with steroid resistant nephrotic syndrome identified 4 (PAN) and 15 (adriamycin) alternatively spliced genes (**Table 2b**). In summary, our data reflects on a new mode of regulation of genes with established causal roles in human glomerular disease (**Table 2c, Supplementary TableS3**). Additionally, this study uncovered several novel genes without any prior established link to mutations in human glomerular disease, such as *Tjp1* and *Itm2b* (**Table 2d, Supplementary TableS4**).

**Table 2:**
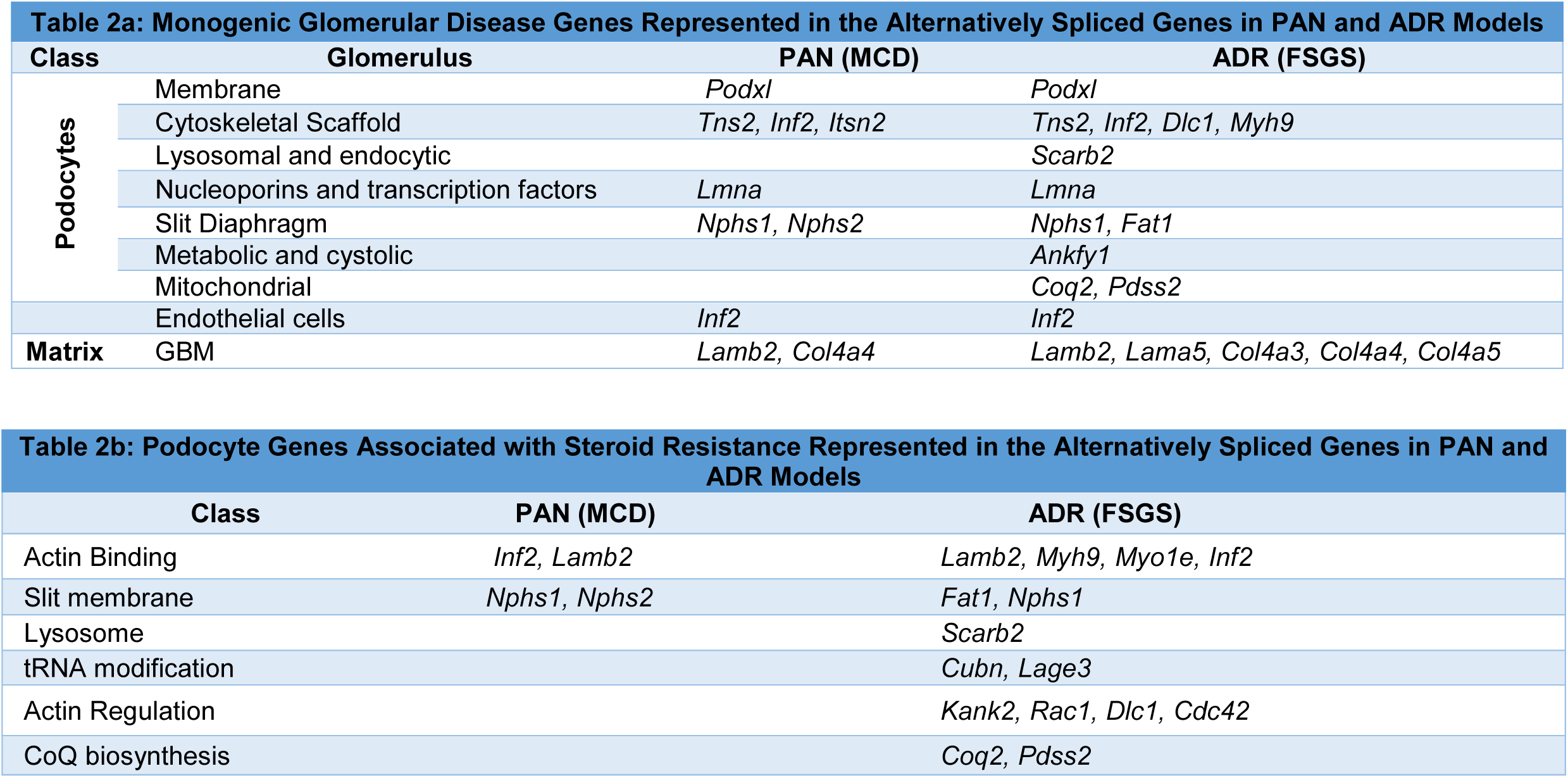
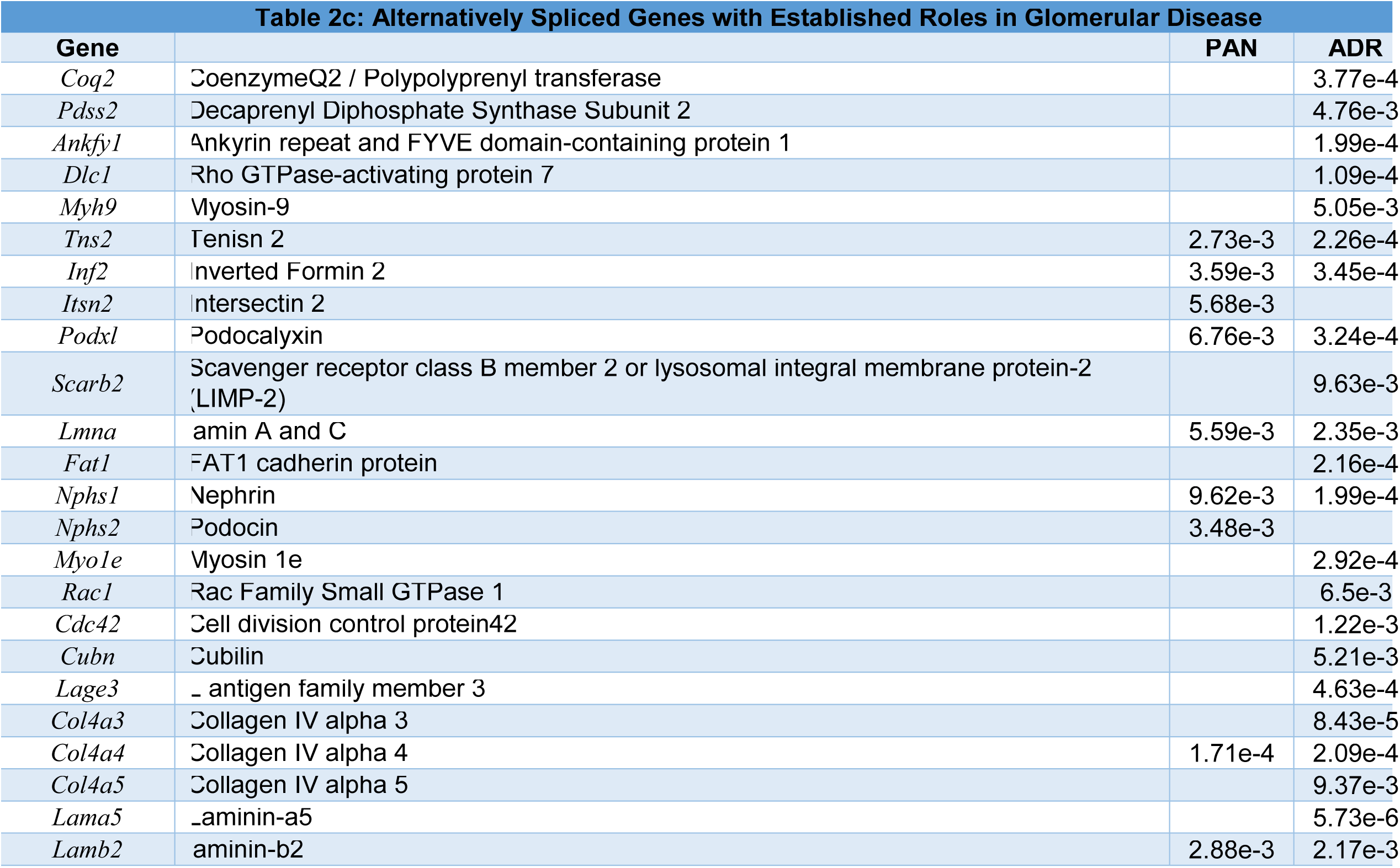
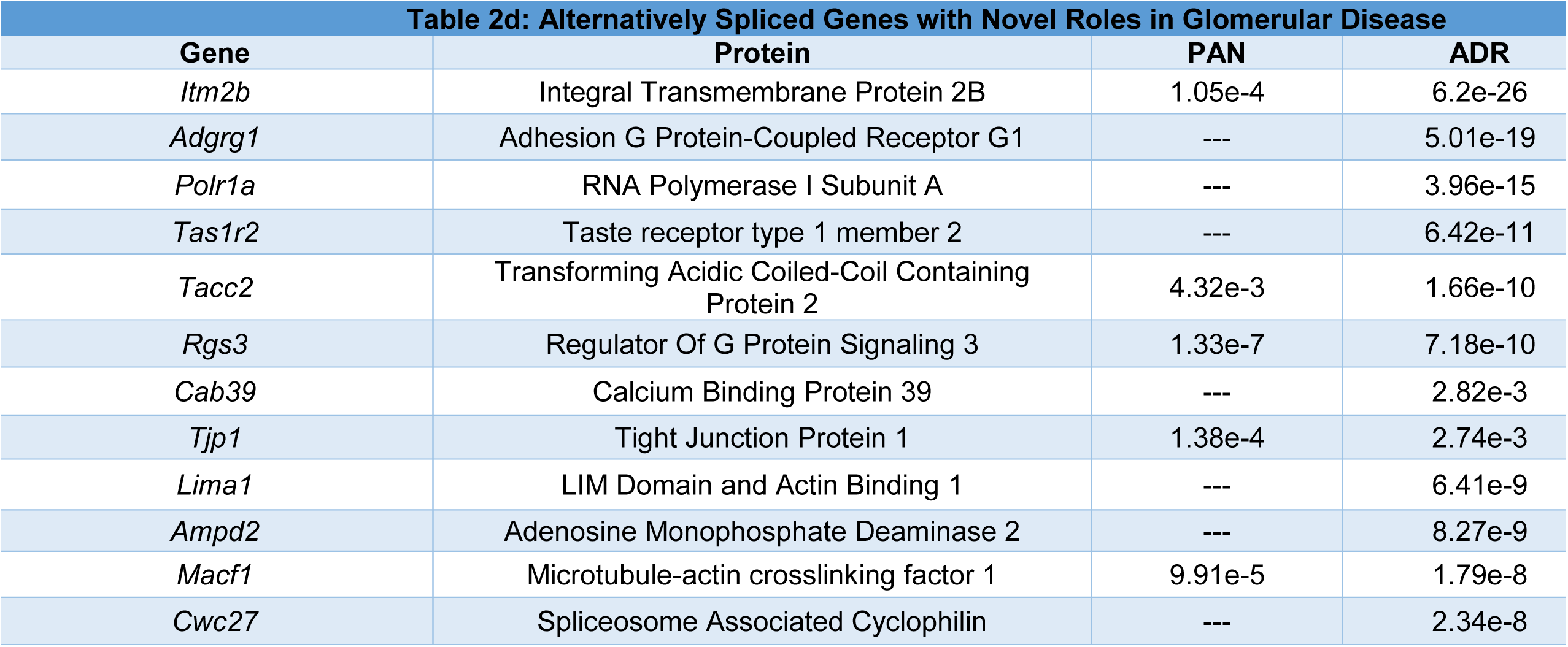
Alternatively Spliced Genes.

Furthermore, we identified many of the alternative splicing events reversed with proteinuria-reducing treatments^39^, Pio (60) and GQ16 (15), suggesting the potential crucial role of alternative splicing of these genes in regulating podocyte health (**Figure 2G, Supplementary Tables S5,S6**).

### III. mRNA Alternative Polyadenylation Occurred During Glomerular Disease Pathogenesis

#### Glomerular mRNAs Underwent Alternative Polyadenylation during Glomerular Injury and a Global Shift from Proximal to Distal Poly(A) Site Usage

To dissect the role of alternative polyadenylation in podocyte and glomerular biology and pathophysiology, we applied APATrap^38^ to the RNASeq datasets to identify alternatively polyadenylated mRNAs in the PAN and adriamycin-models. Glomerular injury identified alternative polyadenylation in a total of 71 (PAN) and 746 (adriamycin) glomerular genes (adj *p*<0.05) (**Figure 3A**). Among these, 55 genes were shared, while 15 were unique to the PAN model and 689 unique to the adriamycin model. A proximal to distal shift in poly(A) site usage was observed in 61 (PAN) and 715 (adriamycin) genes, and a reverse shift from distal to proximal poly(A) site usage in 10 (PAN) and 31 (adriamycin) genes, suggesting an overall lengthening of the 3’UTR in disease (**Figure 3Bi,ii**).

**Figure 3.**
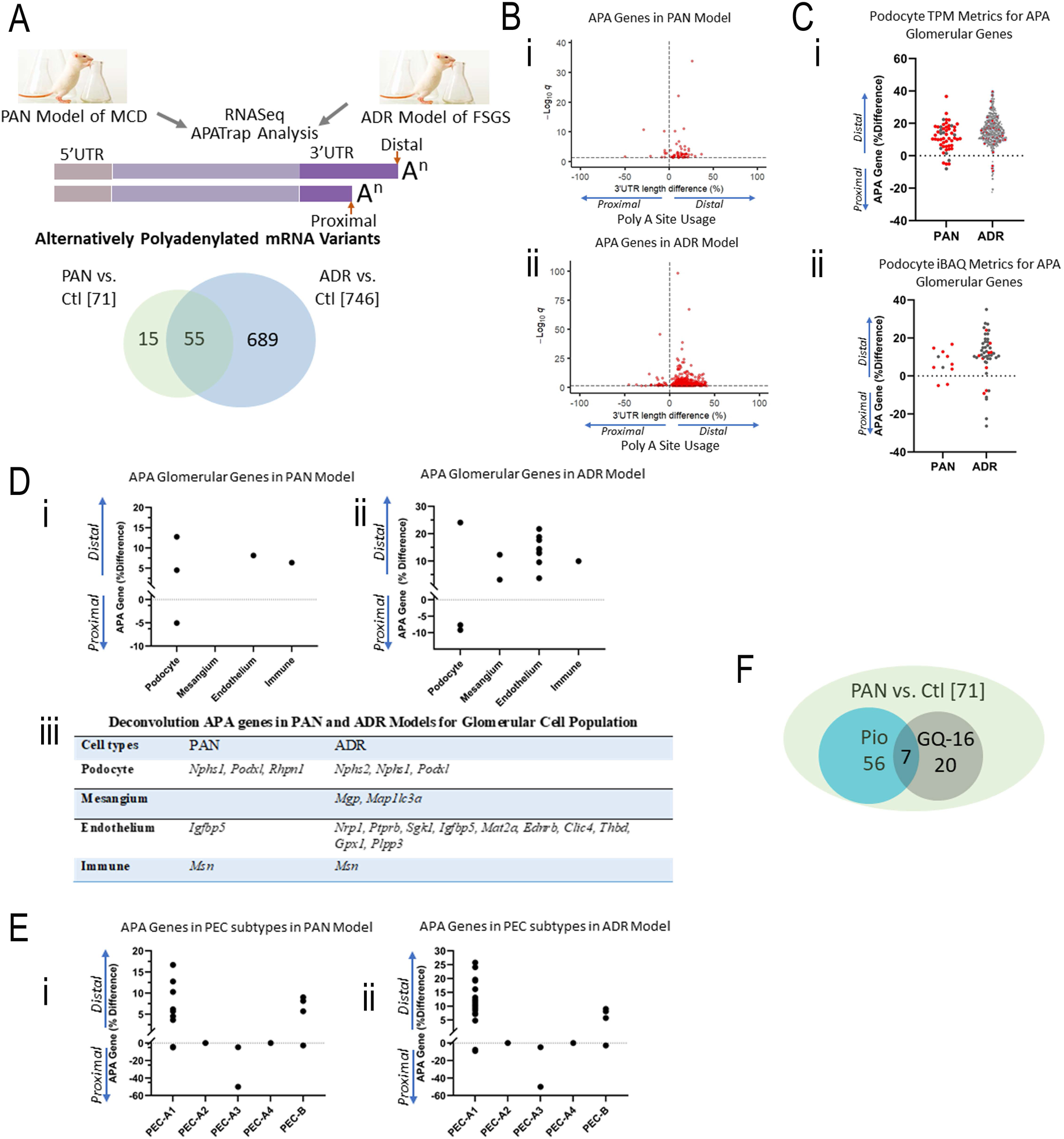
Glomerular mRNAs were alternatively polyadenylated (APA) in puromycin amino-nucleoside (PAN)- and adriamycin (ADR)-induced nephropathy models mimicking minimal change disease (MCD) and FSGS. **(Ai)** Schematic showing the experimental and APATrap bioinformatic workflow to detect APA mRNA transcripts in the glomeruli of rats subjected to PAN- and adriamycin (ADR)-induced injury. Venn diagram depicts the common and distinct alternatively polyadenylated genes detected in both the models compared to healthy rats. **(B)** Volcano plots depict the 3’UTR length difference and a shift towards distal vs proximal poly A site usage for each of the significant alternatively polyadenylated genes observed in the **(Bi)** PAN and **(Bii)** adriamycin (ADR) model. **(C)** Podocyte metrics. **(Ci)** Podocyte-specific transcriptomic dataset of 6593 genes (TPM > 10, *q* < 0.05), identified 55 and 567 APA genes (*q* < 0.05) in PAN and adriamycin (ADR) models. 44 common APA genes between the two models are in red. **(Cii)** Podocyte-specific iBAQ dataset of 551 proteins (*q* < 0.05) identified 11 and 48 APA genes (*q* < 0.05) in PAN and adriamycin (ADR) models. 9 common alternatively polyadenylated genes between the two models are in red. **(D)** Deconvolution distribution of alternatively polyadenylated genes across the different glomerular cell types, including podocytes, mesangial cells, endothelial cells and a small fraction of immune cells in **(Di)** PAN and **(Dii)** adriamycin (ADR) models. **(Diii)** Genes from the deconvolution represented in PAN and adriamycin (ADR) models. **(E)** Deconvolution distribution of alternatively polyadenylated genes across the different parietal epithelial cell (PEC) subtypes, including PEC-A1, PEC-A2, PEC-A3, PEC-A4 and PEC-B population, in **(Ei)** PAN and **(Eii)** adriamycin (ADR) models. **(F)** Venn Diagram showing alternatively polyadenylated genes reversed with Pio and GQ-16 treatments in PAN model.

#### Robust Podocyte Metrics and Contribution from Other Glomerular Cells Identified for Alternatively Polyadenylated Events

Deconvolution of the APATrap-derived dataset by cross-referencing with the podocyte-specific transcriptomic dataset (6593 genes; TPM>10, *q*<0.05)^40^, identified 55 (PAN), 567 (adriamycin) and 44 (PAN and adriamycin) alternatively polyadenylated genes (*q*<0.05) (**Figure 3Ci**). From the podocyte-specific iBAQ dataset (551 proteins; *q*<0.05)^40^, we identified 11 (PAN), 48 (adriamycin) and 9 (PAN and adriamycin) alternatively polyadenylated genes (*q*<0.05) (**Figure 3Cii**). Deconvolution with glomerular cell-type dataset^41^ identified <u>Podocyte</u> specific genes *Nphs1* and *Podxl1* in both models, while the adriamycin-model presented with additional alternative polyadenylation in *Nphs2* (**Figure 3Di-iii**). <u>Mesangial-</u>specific gene *Map1lc3a* and <u>endothelial</u> specific gene *Thbd* showed alternative polyadenylation in adriamycin. Deconvolution by parietal epithelial sub-populations^42^ identified alternative polyadenylation events in mostly podocyte-progenitor A1 sub-population, both in PAN and adriamycin-models (**Figure 3Ei, ii, Supplementary TableS7**).

#### Glomerular mRNAs Underwent Alternative Polyadenylation in Genes with Novel and Established Roles in Glomerulopathies, with Varied miRNA Targeted Sites

Comparison of alternative polyadenylation-dataset in glomerular disease with 80 causal genes in human patients^43^ identified 7 (PAN) and 10 (adriamycin) alternatively polyadenylated genes (**Table 3a)**. These mostly localized within the podocytes and in the slit diaphragm. Among these, 6 were common (*Podxl, Myh9, Kirrel1/Neph1, Nphs1, Nphs2, Cd151)*, while 4 were found only in the adriamycin-model (*Dlc1, Scarb2, Wt1, Thbd*). *Lmx1b* was uniquely identified in the PAN model. Comparison with 49 podocytopathy genes associated with steroid resistant nephrotic syndrome^44^ revealed 3 (PAN) and 7 (adriamycin) alternatively polyadenylated genes (**Table 3b**). In summary, our data on glomerular alternative polyadenylation in the two disease models suggested an additional layer of regulation of genes with established causal roles in human glomerular disease **(Table 3c, Supplementary Table 8)**. Additionally, this study uncovered novel genes without any prior established causal roles from human glomerular disease **(Table 3d, Supplementary Table 9)**. For each alternatively polyadenylated gene, predicted miRNA sites in the 3’UTR were identified (miRDB.org).

**Table 3:**
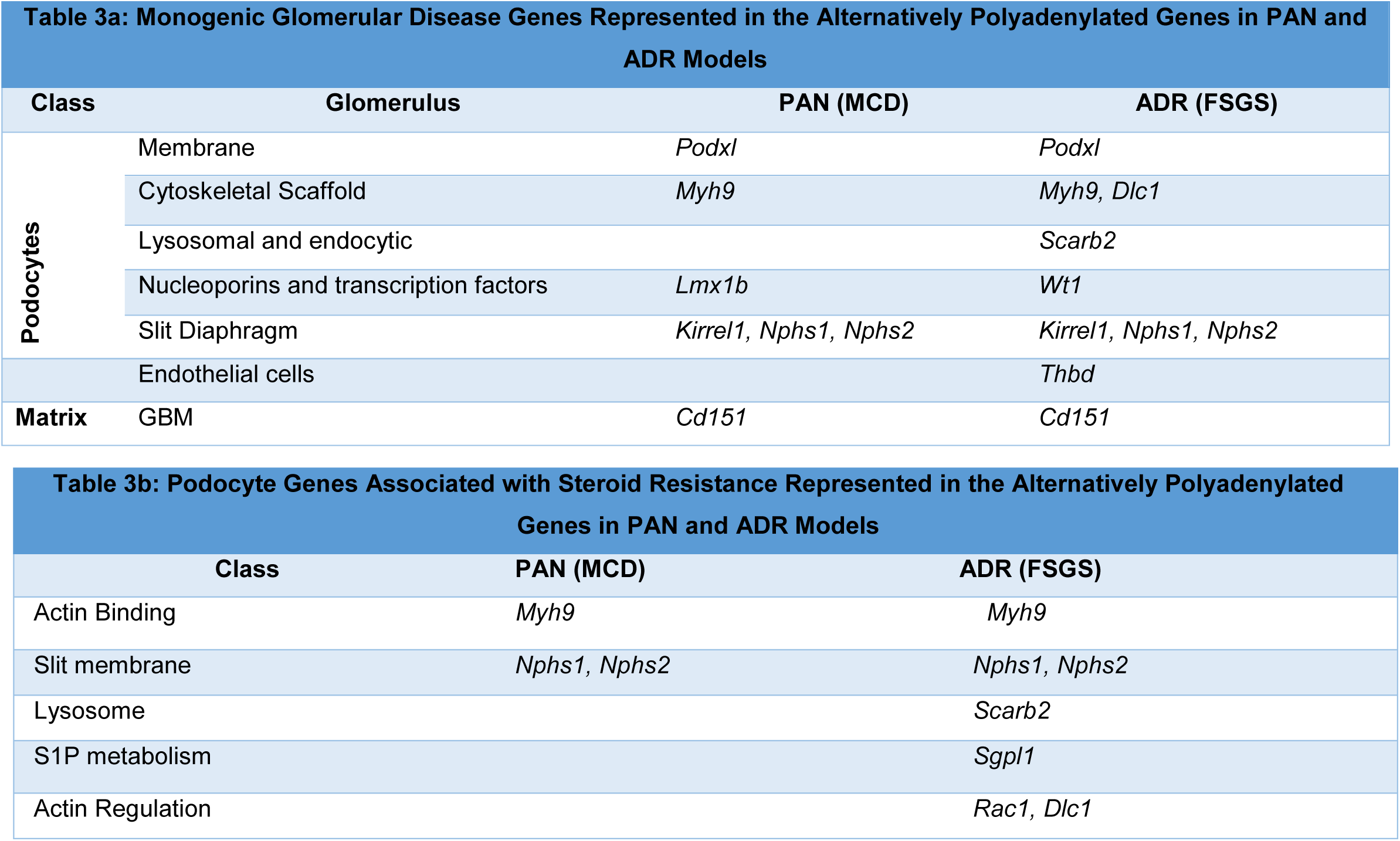
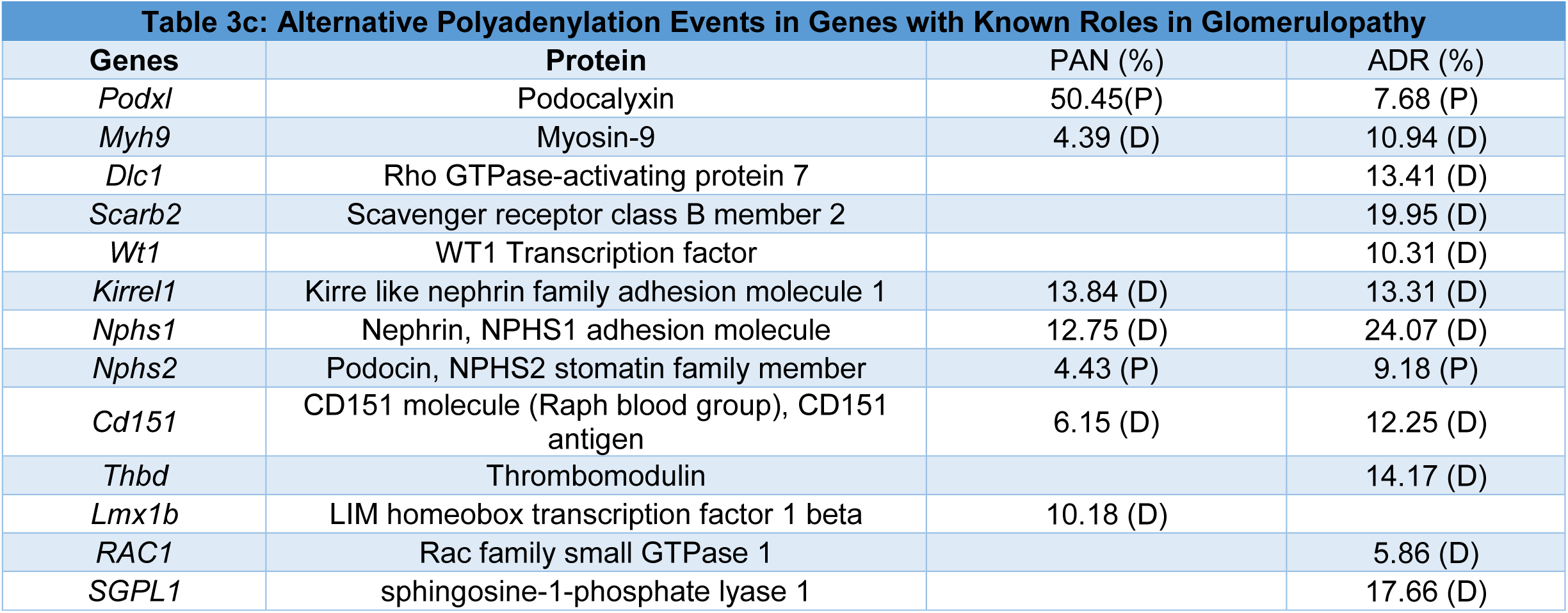
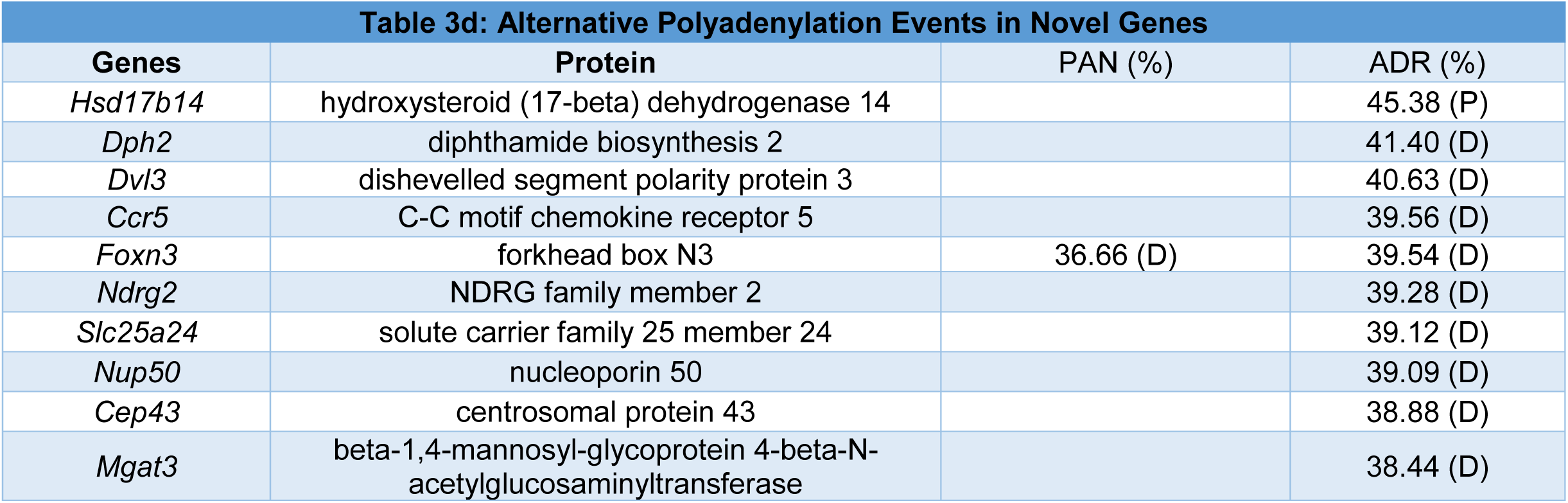
Alternatively Polyadenylated Genes.

Furthermore, we identified many of the alternative polyadenylation events (e.g. *Kirrel1*) reversed with proteinuria-reducing treatments^39^, Pio (56) and GQ-16 (20), suggesting the potential crucial role of alternative polyadenylation of these genes in regulating podocyte health (**Figure 3F, Supplementary Tables S10,S11**).

### IV. Trans-elements such as RNA Binding Proteins and Splicing and Polyadenylation Factors were Dysregulated in Glomerular Disease Models

#### RNA Binding Proteins (RBPs) and Splicing Factors were Dysregulated in Glomerulopathies

Due to complete lack of information on the possible trans-regulatory roles of RBPs and splicing factors in regulating alternative splicing in podocytes, we compiled a database of ∼400 RBPs that play an established role in mRNA splicing^45, 46^ and cross-referenced with the DEGs (adj*p*<0.05,FC>1<1) in PAN and adriamycin-model datasets (**Figure 4Ai,ii, Supplementary TableS12**). 112 (PAN) and 155 (adriamycin) splicing factors were upregulated (96 common), including U1/2/4/5/6/11/12 small nuclear ribonucleoproteins (snRNPs), splicing factors recruited in A, B, and C complexes, specific alternative splicing factors, and RNA binding motif (RBM) proteins.

**Figure 4.**
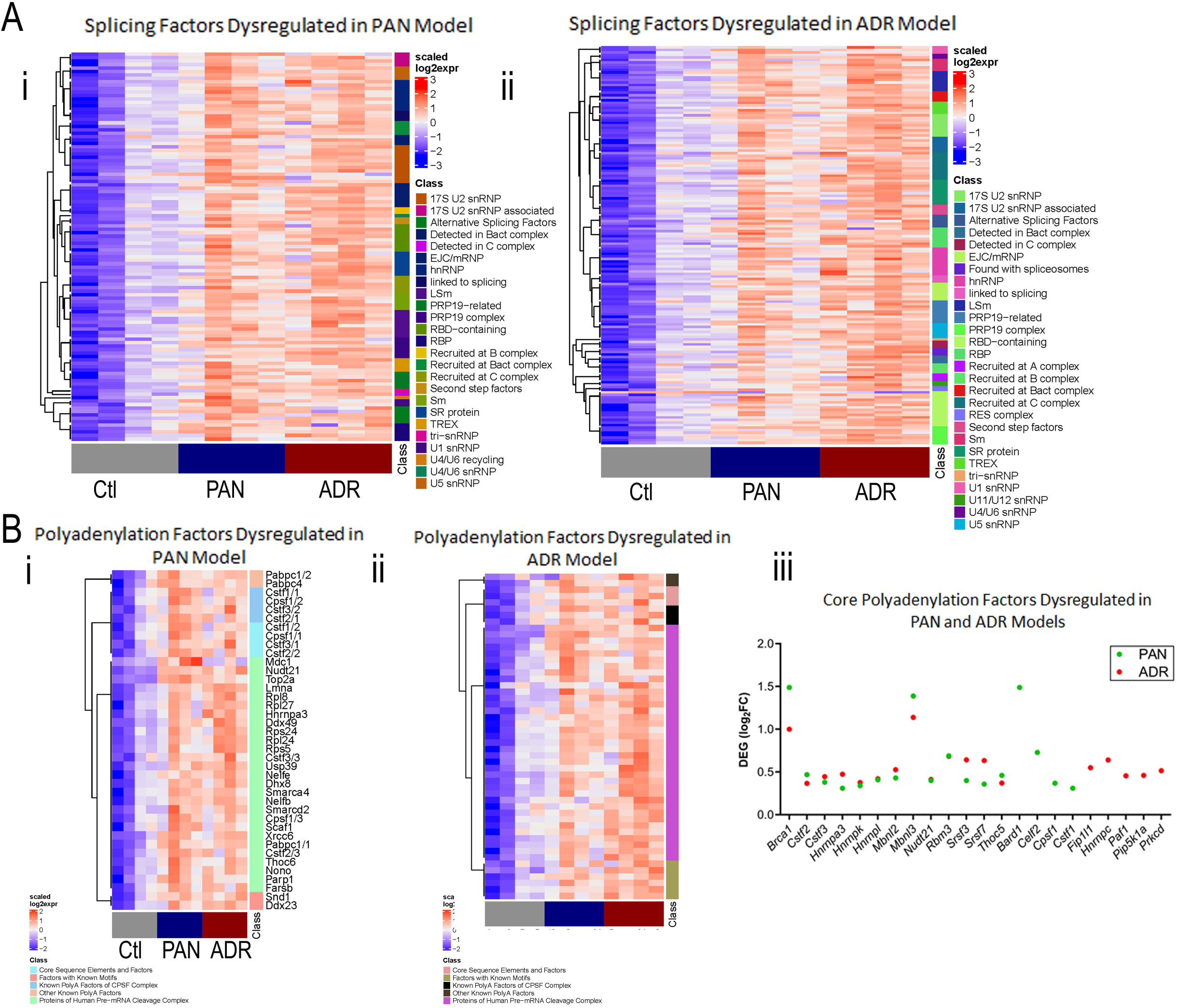
A multitude of splicing and polyadenylation factors, RNA binding proteins and 3’ processing factors were dysregulated in the PAN and adriamycin (ADR) models of glomerular disease. **(A)** Heatmap of DEGs in the PAN and adriamycin (ADR) models represented in the dataset of ∼400 splicing factors and RNA binding proteins (compiled for this study) that play an established role in mRNA splicing. **(i)** Cross-referenced with the DEGs (adj *p* <0.05, FC>1, <1) in the PAN RNASeq dataset, **(ii)** Cross-referenced with the DEGs (adj *p* <0.05, FC>1, <1) in the adriamycin (ADR) RNASeq dataset. **(B)** Heatmap of DEGs in the PAN and adriamycin (ADR) models represented in the dataset of ∼209 pre-mRNA 3’ processing and polyadenylation factors (compiled for this study) **(i)** Cross-referenced with the DEGs (adj *p* <0.05, FC>1, <1) in the PAN RNASeq dataset, **(ii)** Cross-referenced with the DEGs (adj *p* <0.05, FC>1, <1) in the adriamycin (ADR) RNASeq dataset. **(Biii)** “Core” polyadenylation factors cross-referenced with the DEGs (adj *p* <0.05, FC>1, <1) in the PAN and adriamycin (ADR) RNASeq dataset.

#### Pre-mRNA 3’ Processing and Polyadenylation Factors were Dysregulated in Glomerulopathies

The molecular mechanisms underlying APA involve variations in the concentration of core 3’ end processing factors and several RBPs^47^. To evaluate the contribution of these trans-elements in alternative polyadenylation events during glomerular disease, we compiled a database of core and ∼209 factors of the polyadenylation machinery^47–49^ and cross-referenced with the DEGs (adj*p*<0.05,FC>1<1) in PAN and adriamycin-model datasets (**Figure 4Bi,ii,iii Supplementary TableS13**). 39 (PAN) and 51 (adriamycin) polyadenylation factors were upregulated (36 common). These factors included various proteins of the pre-mRNA cleavage complex. Additional transcription and translation machinery factors were identified to be dysregulated in both the models (**Supplementary Figure S6, TableS14**).

The above data combined suggest a potential trans-regulatory mechanism of the observed alternative splicing and alternative polyadenylation events in glomerular disease models.

Additionally, cross-referencing of the GWAS signals from multi-population study (38,463 participants, 2440 cases) of SNPs associated with steroid-sensitive nephrotic syndrome (NS)^50^ with the genomic region of 136 alternatively spliced genes and 3’ UTRs of 71 alternatively polyadenylated genes in the PAN-minimal change disease model dataset identified potential SNPs that may serve as cis-elements for alternative mRNA processing **(Supplementary Figure S7)**.

### V. Alternative Splicing of the Core Slit Diaphragm Components TJP1 and ITM2B Revealed Downstream Alterations

Glomerular disease is characterized by podocyte injury and disruption of the slit diaphragm. TJP1/ZO1 (Tight junction protein 1), and ITM2B (Integral membrane protein 2b) have been identified as core and crucial components of the slit diaphragm^51–57^. Intrigued by the identification of these components in our alternative splicing dataset, we further validated and characterized the consequences of these events at RNA and protein level. NCBI genome browser plots of Rat *Itm2b* and *Tjp1* were constructed depicting mean normalized coverage obtained from the JunctionSeq and RNASeq datasets from control and disease groups (**Figure 5Ai, Bi**). Genomic organization of human *ITM2B* and *TJP1* gene exons show that alternative splicing of *ITM2B* uses either the ‘short’ (S) or the ‘long’ (L) Exon1, producing variants S or L (**Figure 5Aii**), and alternative splicing of *TJP1* makes variants 1 and 2 (with Ex20+ and Ex20Δ) (**Figure 5Bii**). Using primers for RT-PCR and qRT-PCR to capture alternatively spliced variants in human podocytes, a shift in the increased S/L ratio for *ITM2B* and decreased Ex20+/Δ ratio for *TJP1* was observed with injury (**Figure 5Aii,iii**, **5Bii,iii**). Design and use of chemically-modified splice switching oligonucleotides (SSOs) to sterically block the Int19/Ex20 junction, was able to reduce Ex20 inclusion from ∼30% to 9% in a dose dependent manner and thus modulate *TJP1* splicing (**Figure 5Bii)**. Antibodies against ITM2B captured the L form, proteolytically cleaved fragments^58^ and possibly the S form (similar in size to the N-terminal cleaved fragment) (**Figure 5Aiv)**. Moreover, the L form and N terminal cleaved fragment reduced while the short-cleaved fragment increased with adriamycin injury. Antibodies against TJP1 captured the α+ (Ex20+) and αΔ (Ex20Δ) forms, showing a reduction in α+ with PAN and adriamycin injury (**Figure 5Biv)**. Furthermore, most ITM2B co-localized with α-tubulin in differentiated podocytes, indicating its association with microtubules (**Figure 5Av)**. Antibodies capturing the total (L+S) and L-specific ITM2B revealed the presence of both L and S forms in podocytes, reduction of cytosolic L form and shift of S form towards perinuclear and nuclear region during injury (**Figure 5Av)**. For TJP1, while the 20+ form expressed uniformly in the glomeruli and tubular epithelium, the 20Δ form is highly expressed in the glomeruli relative to other cells of the kidney (also seen in human protein atlas database) (**Figure 5Bv**). Moreover, a reduction was observed of the 20+ form (after accounting for adriamycin auto-fluorescence), and increase of the 20Δ form specifically at junction regions with injury, which was more pronounced with adriamycin.

**Figure 5.**
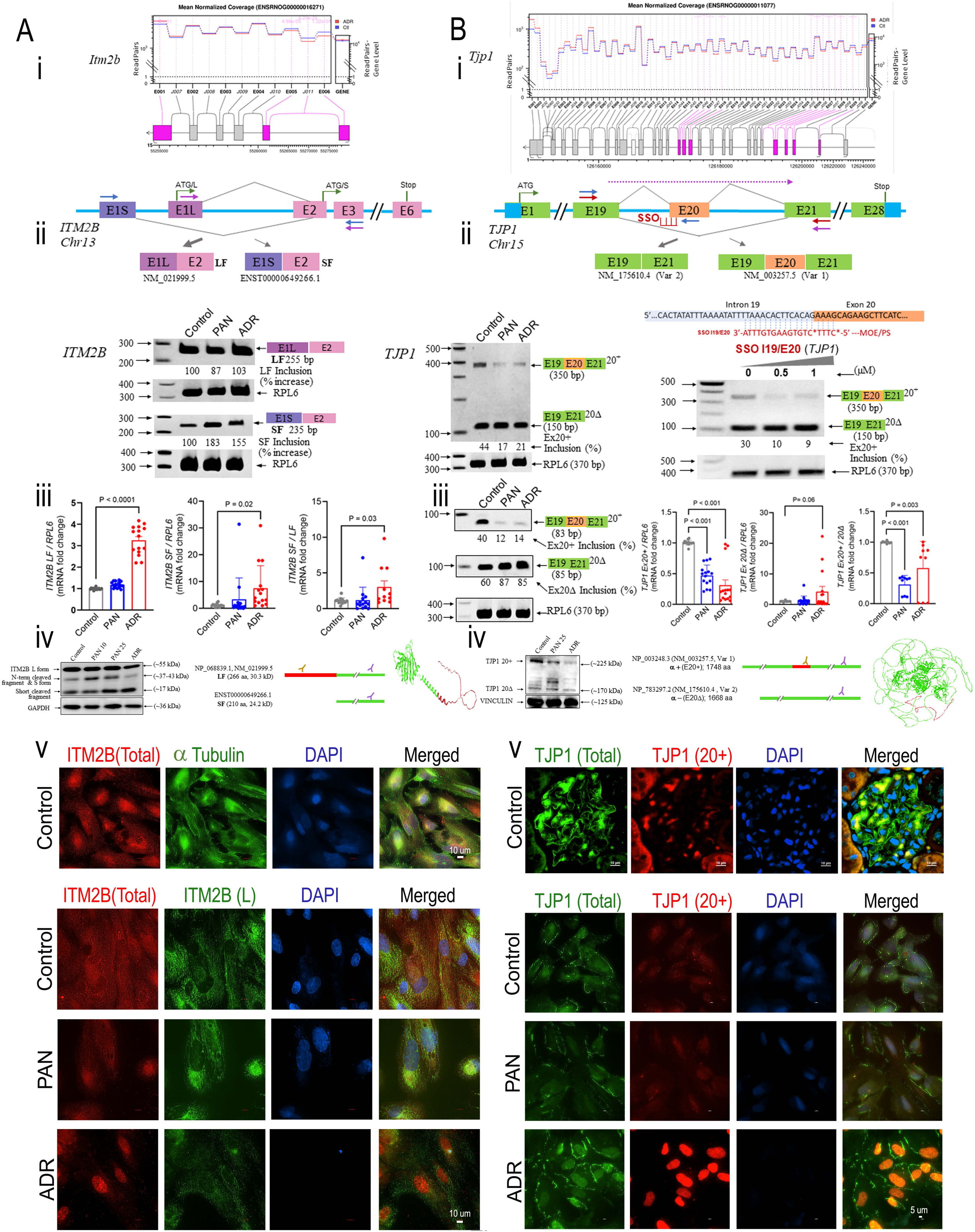
Alternative splicing (AS) of the core slit diaphragm components TJP1 and ITM2B revealed downstream alterations. **(Ai, Bi)** NCBI genome browser plots were constructed from the Rnor_6.0 primary assembly for mean normalized coverage (n=4 each group) obtained from the JuntionSeq and RNASeq datasets for glomeruli from control and adriamycin (ADR)-injured rats. Reads for each exon and junction (left y axis), and the entire gene (right y axis) is depicted for **(Ai)** *Itm2b* and **(Bi)** *Tjp1* for control (blue bars) and adriamycin (ADR)-injured glomeruli (red bars). Due to reverse gene orientation, the exon numbers are depicted in reverse. **(Aii, Bii)** Genomic organization of human *ITM2B* and *TJP1* gene exons, highlighting the alternatively spliced exons, primers, translation start and stop sites for each variant are depicted. **(Aii)** Alternative splicing of *ITM2B* makes use of either the ‘short’ (S) or the ‘long’ (L) E1, producing transcript variant forms S or L. These use different ATG start codons, resulting in-frame but N-terminally different protein forms S and L. **(Bii)** Alternative splicing of *TJP1* gives rise to transcript variant forms 1 and 2 (with Ex20+ and Ex20Δ), which results in different, but in-frame protein forms α+ and α−. **(Aii, iii and Bii, iii)** RT-PCR and qRT-PCR assays using specific primer pairs (shown as arrows on genes) to detect L (purple arrows) and S (blue arrows) splice variant forms for *ITM2B* **(Aii)** and both Ex20+ and Ex20Δ as doublet (red arrows), or specific Ex20+ (blue arrows) and Ex20Δ (purple arrows) spliced forms for *TJP1* **(Bii)** are shown in control, PAN-injured and adriamycin (ADR)-injured human podocytes and % inclusion of variant forms calculated. *RPL6* is shown and used as the housekeeping gene. An 87 ↓ and 103% ↑ in the expression of the L variant, and an even higher 183%↑ and 155% ↑in the expression of the S form of the *ITM2B* variants was detected in PAN- and adriamycin (ADR)-injured podocytes, respectively, indicating an overall shift in ↑S/L ratio with podocyte injury. The two *TJP1* variant forms 1 and 2 (with Ex20+ and Ex20Δ), could be captured with a common primer pair and a clear reduction in the % exon 20 inclusion from 44% to 17% and 21% was observed in PAN- and adriamycin (ADR)-injured podocytes, respectively. qRT-PCR experiments for specific variants corroborated these results. Chemically modified splice switching oligonucleotides (SSO) designed to sterically block the intron 19/ exon 20 junction site was able to show reduction in Ex20 inclusion from ∼30% to 8-9% in differentiated podocytes 48 hrs after transfection using Lipofectamine Ltx. The SSO sequence, modifications, complementarity and binding site on TJP1 pre-mRNA is depicted in red. **(Aiv, Biv)** Western blot assays to capture different protein forms of ITM2B and TJP1 are shown in Control, PAN (10 and 25 µg/ml, 48 hrs) and Adriamycin (ADR, 5µg/ml, 48 hrs) injured podocytes. **(Aiv)** Antibodies against total ITM2B captured the L form, cleaved fragments (ITM2B also gets proteolytically cleaved to give rise to different protein fragments in addition to L and S spliced forms) and possibly the S form (similar in size to the N-terminal Cleaved fragment). The L form and N terminal cleaved fragment reduced with adriamycin injury, while the short-cleaved fragment seems to increase in injury. The L encoded protein segment is highlighted in red in the ITM2B protein structure derived from ‘Sequence coverage visualizer’. **(Biv)** Antibodies against TJP1 captured the α+ (Ex20+) and αΔ (Ex20Δ) forms, with a reduction in α+ with PAN and adriamycin injury. The Ex20 encoded protein fragment is highlighted in red in the α+ form of TJP1/ZO1 protein structure derived from ‘Sequence coverage visualizer’. Antibodies able to capture the total (purple) and specific spliced forms (brown) used in the study are also shown in the schematic. The correct band sizes for ITM2B and TJP1 were estimated from fusion proteins generated in our laboratory (data not shown). **(Av, Bv)** Representative images of immunofluorescence microscopy of podocytes and glomeruli using ‘total’ and ‘spliced form specific’ antibodies against ITM2B and TJP1 are shown. **(Av)** ITM2B total (red), α-tubulin (green) and DAPI (blue) are shown in healthy differentiated human podocytes (upper panel). Scale bar 10 µm. Majority of ITM2B co-localized with α-tubulin in differentiated podocytes, indicating the association of ITM2B form with microtubules. ITM2B total (red), ITM2B L form (green) and DAPI (blue) are shown in Control, PAN (25µg/ml, 48 hours) and adriamycin (ADR, 5 µg/ml, 48 hours) injured podocytes (lower panel). Scale bar 10 µm. Both L and S forms could be detected in podocytes, with reduction of cytosolic L form and shift of S form towards nucleus during injury. Additional nuclear staining was observed due to auto-fluorescent property of adriamycin. **(Bv)** TJP1 total (green), TJP1 20+ (red) and DAPI (blue) are shown in glomeruli from healthy rat (upper panel). Scale bar 10 µm. The 20+ form seemed to be expressed uniformly in the glomeruli and tubular epithelium, while the 20Δ form was highly expressed in the glomeruli relative to other cells of the kidney (also seen in human protein atlas database). TJP1 total (green), TJP1 20+ (red) and DAPI (blue) are shown in Control, PAN (25µg/ml, 48 hours) and adriamycin (ADR, 5 µg/ml, 48 hours) injured podocytes (lower panel). Scale bar 5 µm. A clear reduction was observed of the 20+ form (after accounting for the auto-fluorescent property of adriamycin in the nucleus), and increase of the 20Δ form specifically at junction regions with injury, which was more pronounced in Adriamycin-induced injury.

Overall, these findings suggested that alternative splicing of *ITM2B* and *TJP1* resulted in different protein forms with differential localization and functions in healthy and injured podocytes.

### VI. Alternative Polyadenylation of the Core Slit Diaphragm Components, NPHS1, NPHS2 and NEPH1 Revealed Differential miRNA Binding Sites

Intrigued by the identification of *Nphs1*, *Nphs2* and *Neph1 (Kirrel1)* in our alternative polyadenylation dataset, we further characterized the consequences of these events by first plotting the reads obtained in our datasets at the polyA sites (**Figure 6Ai,iii,v; Supplementary FigureS8**) and secondly with the 3’ RACE and 3’UTR RT-PCR assays (**Figure 6A,ii,iv,vi**). Overall, the 3’ RACE and the APAtrap analyses suggested relative distal (*Neph1* and *Nphs1*) and proximal (*Nphs2*) shifts in poly(A) site usage. Furthermore, we characterized the potential consequences of alternative poly(A) site usage by identifying miRNAs with strong binding sites between the proximal and distal poly(A) sites of *Nphs1* and *Nphs2* (**Figure 6B**).

**Figure 6.**
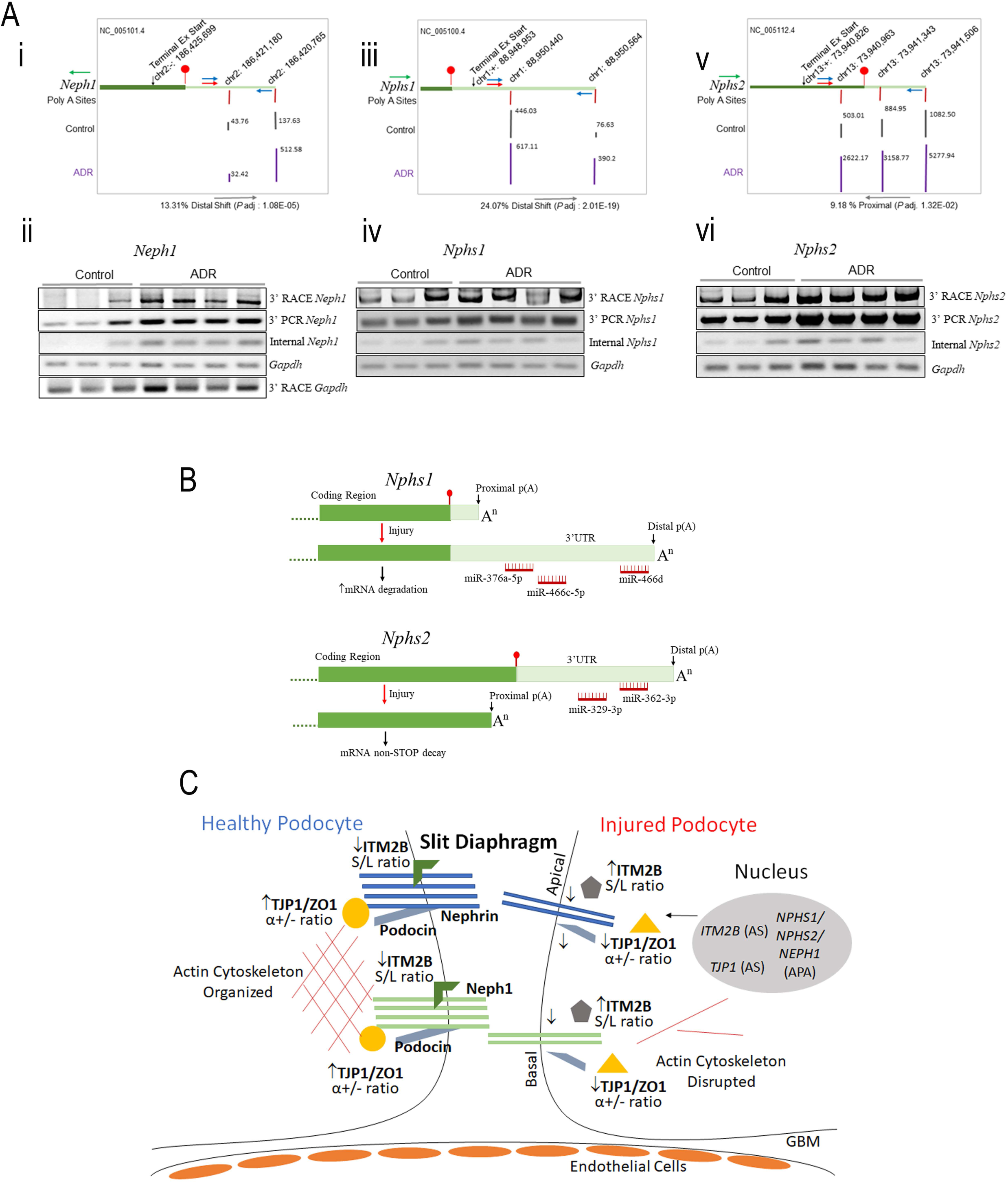
Alternative polyadenylation (APA) of the core slit-diaphragm components, **(Ai, ii)** *Neph1*, **(Aiii, iv)** *Nphs1*, and **(Av, vi)** *Nphs2* mRNAs. First, NCBI genome browser plots were constructed for *Neph1*, *Nphs1* and *Nphs2* using the Rnor_6.0 primary assembly and the reads obtained in our datasets were plotted at the polyA sites. Gene orientation is depicted by green arrow. The tracks show gene, terminal exon start site and polyA sites (red bars), translation stop site (red circle and bar) on the chromosome with ‘+’ ‘-‘ gene orientation. Mean number of reads (n=4 each group) are depicted for control (grey bars) and adriamycin (ADR)-injured glomeruli (purple bars). Second, primers were designed for 3’RACE, 3’UTR RT-PCR and internal RT-PCR assays to detect the products that could result from proximal vs distal polyA site usage in the rat glomeruli of control healthy and adriamycin (ADR)-injured rats. Gels show distal bands for 3’ RACE assays and RT-PCRs for 3’ UTR and internal regions for each gene and housekeeping *Gapdh*. Primers are shown on the gene tracks (blue for 3’ UTR PCR and red for 3’ RACE assay). Semi-quantitative analysis of the distal amplification bands obtained from these reactions (confirmed by sequencing, Supplementary Data) was performed with densitometry, and these were further normalized to internal amplification bands to account for increased adaptive expression observed in all the 3 genes examined. While, *Gapdh* polyA site usage remained similar in Control and adriamycin (ADR)-injured rats, ∼ 364% ↑ (*P*=0.02) in *Neph1* distal band, a trend of ∼ 365% ↑ in *Nphs1* distal band, and a trend of ∼ 70% ↓ in *Nphs2* distal band was detected. Overall, the 3’ RACE and the APAtrap analyses suggested relative distal (lengthening of *Neph1* and *Nphs1* mRNAs) and proximal (shortening of *Nphs2* mRNA) shifts in polyA site usage. **(B)** microRNA binding targets between the proximal and distal poly(A) sites of *Nphs1* and *Nphs2* mRNAs. In *Nphs1* mRNA, poly (A) site usage shifts to a distal site during injury, leading to a longer 3’ UTR. This extended 3’ UTR potentially alters post-transcriptional regulation by facilitating microRNA (miR) targeting (shown in red). These identified miRs include: miR-466d (binding score, 100), miR-376a-5p (score, 88) and miR-466c-5p (score, 76). In *Nphs2* mRNA, poly (A) site usage shifts to a proximal site during injury. This site is within exon 8, resulting in the loss of the canonical stop codon and potentially triggering non-stop decay. In addition, this shortened 3’ UTR potentially alters post-transcriptional regulation by removing microRNA (miR) targeting (shown in red). These identified miRs include: miR-362-3p (score, 86) and miR-329-3p (score, 86), amongst others (not depicted) with varying scores. **(C)** Schematic showing the possible roles of alternatively splicing of *TJP1* and *ITM2B* and alternative polyadenylation of *NEPH1*, *NPHS1* and *NPHS2* and in podocyte cytoskeleton and slit diaphragm regulation. In healthy podocytes, the core transmembrane components of slit diaphragm, NEPHRIN, PODOCIN, NEPH1, TJP1/ZO1 and ITM2B interact with each other and the actin cytoskeleton to maintain the structural and functional integrity of podocytes and the slit diaphragm. These healthy interactions involve relative proximal polyA site usage of *Neph1* and *Nphs1*, distal polyA site usage of *Nphs2*, and ↑*TJP1/ZO1 α+/α−* ratio and ↓*ITM2B S/L* ratios, which are reversed in injured podocytes due to alternative polyadenylation and alternative splicing. This is accompanied with decreased interactions with each other, and disruption of cytoskeleton and the slit diaphragm.

## Discussion

RNA therapeutics is a fast-evolving field, and in the past decade several pre-clinical and clinical studies have guided the potential for RNA therapies to treat different diseases^59–62^. Despite these advances, there is a considerable knowledge gap and unmet need to guide these concepts to podocytopathies. This stems from an absolute lack of understanding of mRNA processing, such as alternative splicing and alternative polyadenylation in glomerular pathophysiology. We utilized two well-established models of glomerular disease for minimal change disease and FSGS, to decipher the transcriptomic landscape at the level of the constitution and heterogeneity of mRNAs due to alternative splicing and alternative polyadenylation, beyond the transcript quantification. We identified several alternatively spliced and alternatively polyadenylated events, which potentially offer critical layers of gene regulation. Such events have been recognized as important steps in cellular development, normal physiology, and in pathological processes in other diseases^13, 14, 19–22, 26–34^. However, this is the first study to probe these mechanisms in podocyte biology and glomerulopathies. A few recent studies have highlighted the role of alternative splicing in proximal tubules in diabetic nephropathy^63^, and in kidney development^64^, but there is no precedence of investigating global alternative splicing and alternative polyadenylation in glomerular disease.

Transcriptomic and the alternative splicing and alternative polyadenylation profiles of minimal change disease model (PAN) and FSGS model (Adriamycin) revealed numerous previously unrecognized molecular alterations to mRNAs in the glomeruli, with differential contribution from podocytes, mesangial and endothelial cells and parietal epithelial subtypes. Moreover, while both glomerular disease types are common causes of idiopathic NS in pediatric and adult population, and are characterized by podocyte injury, they have different disease pathologies, etiologies, disease course and treatment outcome^65, 66^. FSGS, usually presents as a more severe form of treatment-resistant disease, and this was reflected in transcriptomic, alternative splicing and alternative polyadenylation profiles in the current study, as the adriamycin model exhibited more robust alterations. However, despite obvious distinct differences between the two models, we identified many overlapping genes, pathways and alternative splicing and alternative polyadenylation events, which indicated shared underlying mechanisms, and added strength to the notion that these disease forms might be different manifestations of the same progressive disease^67^.

We identified a reversal of many of the alternatively spliced and alternatively polyadenylated events with treatment modalities, Pio and GQ16, which we have previously demonstrated to be beneficial in reducing proteinuria and glomerular injury^39, 68, 69^. This suggested a pathogenic or adaptive role of these events, in addition to an associative role, although future studies are warranted. Additionally, we observed concomitant dysregulation of numerous RBPs, suggesting their trans-regulatory effects on alternative splicing and alternative polyadenylation. Of particular interest was the identification of alternative splicing (*TJP1/ZO1, ITM2B*) and alternative polyadenylation (*NPHS1, NPHS2, NEPH1*) events in podocyte and slit diaphragm genes, which are critical components of the filtration barrier. Podocyte-specific deletion of *Tjp1* has been shown to result in impaired foot process interdigitation, slit diaphragm formation, and glomerular dysfunction^70^, but there are is no indication of the role of *TJP1* variants (α+/α−) in podocytes^71^, despite their differential epithelial distribution^72^, and recently documented importance in stress fiber assembly in another system^73^. ITM2B is a newly identified crucial transmembrane member of the Nephrin and Neph1 interactome and of slit diaphragm^51^, a differential microtubular S vs L interactome of ITM2B has been recently identified in retina^74^, and the loss of its N terminal domain in the S form potentially lacks the transmembrane domain. Our study for the first time demonstrated the altered expression and localization of TJP1 α+ and α− and *ITM2B* S and L forms during podocyte injury. Additionally, we demonstrated altered proximal vs distal poly(A) site usage in *Neph1, Nphs1* and *Nphs2* and potential consequences for microRNA accessibility. Shortening vs lengthening of transcripts due to proximal vs distal polyA site usage has been reported to be variable in other diseases or during differentiation^49, 75–77^. The above described alternative splicing and alternative polyadenylation events point towards a novel and previously uninvestigated mechanisms of regulation of the podocyte and slit diaphragm structure and function (**Figure 6C**). Finally, we have demonstrated the ability of chemically-modified SSO designed for steric blocking, with enhanced stability and efficacy^78^, to modulate an alternative splicing event in podocytes, which provides proof-of-concept to treat glomerulopathies via slit diaphragm mRNA regulation. Notably, such technology paved the way to clinical trials and FDA-approved drugs such as nusinersen (SPINSRAZA*^®^*) for exon inclusion in Spinal Muscular Atrophy ^78, 79^.

There were various strengths and limitations to our study. We utilized well-established models of glomerular disease which mimic human minimal change disease and FSGS remarkably well^80^, and identified critical components of the slit diaphragm and filtration barrier that exhibit alternative splicing and alternative polyadenylation events. Further validation of these targets and identification in human biopsies will provide technical and biological confirmation. However, in addition to high cost, these studies require precise probes and fresh samples to capture the differential variants. It is also critical to validate such targets in human samples or cell lines (as we have done for alternative splicing of TJP1 and ITM2B in the present study), as alternative processing events can be highly species specific^81–83^. A critical barrier is the use of bioinformatics approaches to align and stitch segments of exons, junctions and 3’UTRs to identify targets. For example, we identified differential exon usage in *NPHS1*, *NPHS2* and *PODXL* mRNAs, and evidence from other studies suggests that these podocyte genes indeed exist in spliced forms ^84–87^. However, future studies are needed to understand the functional consequences of these events. Another barrier lies in the fact that often multiple alternative processing events occur on the same transcript, but with conventional RNA-seq approaches, it is not possible to link these events. While we utilized effective and proven approaches at high read depth^36, 38^, new technologies such as long-readSeq, performed with sufficient depth, could provide more complete characterization of mRNA isoforms. Another limitation is that bulk glomerular RNASeq is not effective in delineating cell-specific events, in capturing alternative polyadenylation and alternative splicing in lowly-expressed genes, or in precisely mapping polyA sites, which requires 3’-end focused sequencing approaches. Future use of scRNASeq, if done at sufficient depth, will allow assignment of mRNA alternative polyadenylation events to specific cell types, but would likely not reveal many alternative splicing events due to the shortness of the reads^88–91^. Nevertheless, we have successfully deconvoluted the RNASeq data to robust podocyte-metrics and to glomerular cell-types^41,42^. Moreover, spatial transcriptomics using probes unique for a specific mRNA isoform could also help map alternative splicing and alternative polyadenylation events to specific cell types.

In summary, the analyses of mRNA alternative splicing and alternative polyadenylation in minimal change disease and FSGS models in the present study provides novel insights and mechanisms of glomerular pathophysiology and development of novel therapeutic intervention strategies.

## Acknowledgments

We thank The Ohio State University (OSU) Genomics Core Resources for RNA Sequencing Services and the College of Veterinary Medicine for Clinical Pathology Services. We also thank Rachel Cianciolo from Niche Diagnostics for Pathology Services, Neelu Singh for assistance with PAS staining of kidney slides and Research Histology Core at Stony Brook University. We also thank Matthew Sampson, MD and Michelle McNulty at Harvard Medical School for helping us with accessing the GWAS database from their published studies. These studies were partly supported by the Dialysis Clinic Inc. Award, SUNY Seed Grant Award, and NIH R01 DK133440 Award to SA. This project was also partly supported by funding from the Center for Clinical and Translational Science Genomics Shared Resource Voucher Support to SA which was supported by the NIH Clinical and Translational Science award to The Ohio State University (UL1TR002733) from the National Center for Advancing Translational Sciences, United States.

## Author Contributions

MKD performed experiments, analyzed and interpreted the data, prepared figures and tables, and drafted parts of the manuscript. AW and LP performed bioinformatics analyses, prepared figures and tables, and drafted parts of the manuscript. MY, CY and CB performed experiments and prepared figures and tables. RG and CM participated in study design, data interpretation and manuscript editing. SA conceptualized and designed the study, analyzed and interpreted the data, prepared figures and tables, drafted and edited the manuscript, and acquired funding. All the authors approve of the final version of the manuscript. Part of these findings were presented at the 14^th^ International Podocyte Conference, held at University of Pennsylvania, Philadelphia, PA in 2023, at the ASN-Kidney Week held in Philadelphia, PA in 2023, and at the RNA Society Annual Meeting held in Edinburgh, Scotland in 2024.

## Data Sharing

RNA-Seq data has been deposited to the Gene Expression Omnibus (GEO) data repository: GSE179945 and GSE286014

## Supplementary Files

**Supplementary Figure S1**

Serum chemistry of PAN- and adriamycin (ADR)-induced nephropathy animal models

**Supplementary Figure S2**

Ontology enrichment analyses of DEGs in PAN- and adriamycin (ADR)-induced nephropathy models

**Supplementary Figure S3**

DEGs in PAN and adriamycin (ADR) models deconvoluted for podocytes, other glomerular cells and parietal epithelial subtypes

**Supplementary Figure S4**

DEGs in PAN and adriamycin (ADR) models represented in established genes for glomerular disease and podocytopathies

**Supplementary Figure S5**

Dispersion plots depicted good fit for the PAN and adriamycin (ADR) models for exon/intron/junction analyses

**Supplementary Figure S6**

Heatmap of DEGs in the PAN and adriamycin (ADR) models represented in the dataset of a combination of transcription and translation machinery factors

**Supplementary Figure S7**

Manhattan plots depicting multi-population GWAS signals/SNPs localized in the regulatory regions of alternatively spliced genes and in the 3’UTR of alternatively polyadenylated genes in the PAN model

**Supplementary Figure S8**

Genomic organization of *Neph1, Nphs1 and Nphs2* genes and 3’UTR, Poly (A) site usage identified, RACE and 3’PCR primers used and sequence chromatograms obtained from RACE assays

**Supplementary Table 1**

Deconvolution of DEGs in PAN and adriamycin (ADR) models for parietal epithelial subtypes

**Supplementary Table 2**

Deconvolution of alternatively spliced genes in PAN and adriamycin (ADR) models for parietal epithelial subtypes

**Supplementary Table 3**

Alternatively spliced genes in PAN and adriamycin (ADR) models represented in monogenic genes with established roles in glomerular disease

**Supplementary Table 4**

Alternatively spliced genes in PAN and adriamycin (ADR) models representing novel genes in glomerular disease

**Supplementary Table 5**

Alternatively spliced genes reversed with pioglitazone treatment

**Supplementary Table 6**

Alternatively spliced genes reversed with GQ-16 treatment

**Supplementary Table 7**

Deconvolution of alternatively polyadenylated genes in PAN and adriamycin (ADR) models for parietal epithelial subtypes

**Supplementary Table 8**

Alternatively polyadenylated genes in PAN and adriamycin (ADR) models represented in monogenic genes with established roles in glomerular disease

**Supplementary Table 9**

Alternatively polyadenylated genes in PAN and adriamycin (ADR) models representing novel genes in glomerular disease

**Supplementary Table 10**

Alternatively polyadenylated genes reversed with pioglitazone treatment

**Supplementary Table 11**

Alternatively polyadenylated genes reversed with GQ-16 treatment

**Supplementary Table 12**

Dysregulated RBPs and splicing factors in PAN and adriamycin (ADR) models

**Supplementary Table 13**

Dysregulated pre-mRNA 3’ processing and polyadenylation factors in PAN and adriamycin (ADR) models

**Supplementary Table 14**

Dysregulated transcription and translation machinery factors in PAN and adriamycin (ADR) models

