## Supplemental Data for "RNA Alternative Splicing and Polyadenylation and Regulation of the Glomerular Filtration Barrier"

**Table of Contents**

**Supplementary Methods**
*Animal Models*
*RNASeq and Differential Expression of Genes (DEGs)*
*JunctionSeq Analyses*
*rMATS Analyses*
*APATrap Analyses*
*Heatmaps, Pathway Analysis and Ontology Enrichment*
*GWAS Correlation with Alternatively Spliced and Alternatively Polyadenylated Genes*
*Podocyte Culture and Treatments*
*Reverse Transcriptase-Polymerase Chain Reaction (RT-PCR)*
*3' Rapid Amplification of cDNA Ends (RACE) Assay*
*Splice-Switching Oligonucleotide (SSO) Design and Assay*
*Quantitative RT-PCR (SYBR)*
*Western Blot Analysis*
*Immunofluorescence (IF) Staining*

**Supplementary Figures**
Supplementary Figure S1
Supplementary Figure S2
Supplementary Figure S3
Supplementary Figure S4
Supplementary Figure S5
Supplementary Figure S6
Supplementary Figure S7
Supplementary Figure S8

**Supplementary Tables**
Supplementary Table S1
Supplementary Table S2
Supplementary Table S3
Supplementary Table S4
Supplementary Table S5
Supplementary Table S6
Supplementary Table S7
Supplementary Table S8
Supplementary Table S9
Supplementary Table S10
Supplementary Table S11
Supplementary Table S12
Supplementary Table S13
Supplementary Table S14

**Supplementary References**

#### Supplementary Methods

*Animal Models*

The animal studies were conducted under the approval and guidelines of the Institution Animal Care and Use Committee at Nationwide Children's Hospital. Male Wistar rats were injected intravenously (IV) with puromycin aminonucleoside (PAN) (Sigma-Aldrich, St. Louis, MO) (50mg/kg) or adriamycin (Sigma-Aldrich) (7.5mg/kg) on Day 0 to model minimal change disease or focal segmental glomerulosclerosis (FSGS), respectively (n=4/group). Control rats were given saline IV injections. Weights was recorded throughout the study, and spot urine and serum were collected twice a week. The PAN- treated and adriamycin-treated rats were euthanized on Day 11 and 21 respectively, at which time kidneys were harvested and glomeruli isolated using the sequential sieving method, as previously described [1]. Glomeruli were embedded in Histogel™ Specimen Processing Gel (Epredia, Kalamazoo, MI), and routinely processed for paraffin embedding, sectioned at 4 µm thickness and viewed with light microscopy for quality assessment. Glomerular isolation quality was also assessed by directly placing them on a slide and visualizing under a microscope. Total RNA was isolated from rat glomeruli using the mirVana Isolation Kit (Life Technologies Corporation, Carlsbad, CA) according to manufacturer's instructions. Yield and purity were calculated using a nanodrop and a Qubit fluorometer. Kidney cross sections were routinely processed, sectioned at 4 µm thickness, and stained with periodic acid-Schiff method (Sigma-Aldrich). Histology was reviewed by a pathologist blinded to treatments. Equal amounts of urine (5µl) collected from rats throughout the study was resolved using sodium dodecyl sulfate-polyacrylamide gel electrophoresis (SDS-PAGE) on an 8% gel. Albumin bands were visualized after staining with Coomassie Brilliant Blue G-250 (Alfa Aesar, Tewksbury, MA). Urine protein:creatinine ration (UPCR) analyses were performed on final day urine by Antech Diagnostics GLP (Morrisville, NC) to quantify proteinuria. Albumin and cholesterol levels in serum were measured on the Vet Axcel (Alfa Wasserman Diagnostic Technologies, LLC, West Caldwell, NJ) using the ACE albumin and cholesterol reagents (Alfa Wasserman Diagnostic Technologies, LLC, West Caldwell, NJ) at The Ohio State University's College of Veterinary Medicine, Clinical Pathology Services, according to manufacturer's instructions.

*RNASeq and Differential Expression of Genes (DEGs)*

To perform RNA sequencing, mRNA libraries were generated using ~200ng total glomerular RNA (quantified using Qubit Fluorometer) with RIN of >7, using NEBNext Ultra II Directional (stranded) RNA Library Prep Kit for Illumina (NEB #E7760L), NEBNext Poly (A) mRNA Magnetic Isolation Module (NEB #E7490) and NEBNext Multiplex Oligos for Illumina Unique Dual Index Primer Pairs (NEB #6442S/L). Sequencing of the libraries was performed with Illumina NovaSeq 6000 SP flow cell using paired-end 150-bp format (300 cycles = 2x150bp) to at least 17 million passed-filter clusters/sample (equivalent to 34 million reads) and using internal pipeline [2], reads were aligned to Rat genome Rnor6.0 with HISAT2 [3]and counts generated for Rnor 6.0 v101 with featureCounts from the subread package [4]. Post alignment quality check (QC) was assessed with fastqc, RseQC, and picard [5] (https://broadinstitute.github.io/picard). Counts were normalized with voom and differential expression tested with limma (Ritchie et al., 2015). Differentially expressed genes (DEGs) were chosen with adjusted $p<0.05$ and abs(logFC)>1. These data have been deposited to the Gene Expression Omnibus (GEO) data repository: GSE179945 and GSE286014.

These datasets and the RNASeq datasets from the pioglitazone and GQ-16 treated rats from our previously reported study [1] (GSE179945) were utilized for JunctionSeq and APATrap analyses described below.

*JunctionSeq Analyses*

For each sample, the following steps were followed for using JunctionSeq analysis [6], also outlined in http://hartleys.github.io/JunctionSeq/doc/example-walkthrough.pdf.Step 1: QoRTs QC was ran to process each sample (runQoRTsQC.pbs per sample for step 1). Step 2: QoRTs R package was used to generate QC figures and generate size factors. Step 3: QoRTs was run to make flat gff for DEXseq and JunctionSeq and to merge with novel splice sites based on samples. And finally, Step 4: JunctionSeq was used to perform differential splicing analysis and to generate plots. All samples were combined for Steps 2-4 (runJunctionSeq.pbs).

*rMATS Analyses*

After alignment, BAM files and a reference GTF file were used as inputs to rMATS for detecting alternative splicing events across Control, PAN and adriamycin conditions. rMATS-turbo identified five major splicing event types: Skipped Exon (SE), Retained Intron (RI), Alternative 5' Splice Site (A5SS), Alternative 3' Splice Site (A3SS), and Mutually Exclusive Exons (MXE). Statistical analysis was performed groupwise to compare splicing events across all three conditions (Control, PAN and adriamycin) simultaneously. Statistically significant events were selected based on an FDR threshold (≤0.05). The results were analyzed and visualized using pie charts to compare alternative splicing patterns across groups.

*APATrap Analyses*

For each sample, the following steps were followed for using APAtrap analysis [7], also outlined in https://sourceforge.net/p/apatrap/wiki/User%20Manual/. Step 1: make bedgraph (make_bedgraph.pbs). For each sample, trim raw fastq for adapters and length with trimmomatic v0.38 Map to Rnor6.0 with hisat2/2.1.0, convert to bam and sort with samtools/1.10, convert bam to bedgraph with bedtools 2.17.0 using genome CoverageBed. Step 2: identify distal 3'UTR (run_identifyDistal3UTR.pbs). With all sample bedgraphs and reference ensembl rn6 bed from UCSC, identify all 3'UTRs present in data. Step 3: predict alternative polyadenylation (text_for_predictAPA_byCompare). With only the samples needed for a comparison and UTR bed from step 2, estimate coverage of UTRs. Limit samples to only comparison due to run time. Step 4: test differential alternative polyadenylation (run_deAPA.R). Generate statistical test results for each measured alternative polyadenylation for each comparison.

*Heatmaps, Pathway Analysis and Ontology Enrichment*

Heatmaps were generated with ComplexHeatmap in R. Enriched canonical pathways were identified using ontology enrichment performed with MOET – MultiOntology Enrichment Tool [MOET, Ontology Enrichment (mcw.edu)] to identify enriched biological processes, cellular components and molecular functions.

*GWAS Correlation with Alternatively Spliced and Alternatively Polyadenylated Genes*

Genes that were identified in the alternative splicing and alternative polyadenylation dataset in the minimal change disease model were cross-referenced for GWAS SNPs identified from patients with steroid sensitive nephrotic syndrome[8] to generate Manhattan plots. For the alternative splicing dataset, any SNP in the gene region covering the exons, introns, 5' and 3' UTRs and splice site junctions was plotted and for the alternative polyadenylation dataset, any SNP in the 3' UTR region was plotted.

*Podocyte Culture and Treatments*

Immortalized human podocytes (gift from Moin Saleem, University of Bristol) were cultured in a humidified atmosphere with 5% CO2 in proliferating conditions at 33°C in RMPI 1640 (Corning, Tewksbury, MA) with 10% fetal bovine serum (FBS), 1% 100 X Penicillin Streptomycin L-Glutamine (PSG), and 1% Insulin-Transferrin-Selenium-Ethanolamine (ITS-X) (Gibco, Gaithersburg, MD), as described previously[9]. They were differentiated at 37°C for 14 days, and treated with vehicle dimethyl sulfoxide (DMSO) (Sigma-Aldrich), puromycin aminonucleoside (PAN) (Sigma-Aldrich) at 10 or 25 ug/mL and with adriamycin (Sigma-Aldrich) at 5 ug/ml for 48 hours. SSO treatment was performed at 0.5 and 1 µM concentrations with transfection reagent Lipofectamine™ RANiMAX (Life Technologies Corporation) for 48 hours. Total RNA was isolated using the TRIzol reagent and RNAeasy kit (Qiagen) according to manufacturer's instructions and yield was calculated using a Nanodrop lite plus (Life Technologies Corporation).

*Reverse Transcriptase-Polymerase Chain Reaction (RT-PCR)*

Total glomerular and podocyte RNA (500 ng – 1 ug) were treated with DNase (Life Technologies Corporation) at room temperature for 15 min, followed by inactivation by 25 mM EDTA at 65°C for 10 min. RNA was reverse transcribed using the iScript cDNA Synthesis Kit (Bio-Rad, Hercules, CA) according to the manufacturer's instructions. Appropriately diluted cDNA was used for reverse transcription-polymerase chain reaction (RT-PCR) using gene specific and house-keeping primers (**Table 1**). The PCR conditions were: 95°C for 5 min, 35 X (95°C for 30 s, 55°C for 30 s, 72°C for 30 s), followed by extension at 72°C for 10 min. Amplified products were resolved on 2-3% agarose gels and quantified by densitometry using ImageJ (National Institutes of Health, Bethesda, MD).

*3' Rapid Amplification of cDNA Ends (RACE) Assay*

Total RNA (DNase-treated) was reverse transcribed using the Supercript II RT and a Poly A adapter primer with a 20-mer adaptor sequence and 18 thymine residues (5'GGCCACGCGTCGACTAGTACTTTTTTTTTTTTTTTTTT-3'), in accordance with the manufacturer's instructions provided with the 3’ RACE Assay kit (Life Technologies Corporation). PCR reaction was performed using the RACE gene-specific forward primer on the 3'UTR close to the first poly(A) site of the target gene, and the universal Poly A reverse primer (5'-GGCCACGCGTCGACTAGTAC-3'), which is identical to the 20-mer adapter sequence in the PolyA adapter primer (**Table 1**). The PCR was run under the following conditions: 95°C for 5 min, 38-40 X (95°C for 30 s, 50°–60°C annealing temperature for 30 s specific to the gene, and 72°C for 30 s) followed by extension at 72°C for 10 min. The amplified products were separated on an agarose gel, and the expected size products were excised from the gel and sequenced (Genomics Core Facility at SBU). Only the bands correctly identified to the specific gene were quantified by densitometry using ImageJ (National Institutes of Health).

*Splice-Switching Oligonucleotide (SSO) Design and Assay*

An 18-mer Splice-Switching Oligonucleotide (SSO_5’ CTTTCTGTGAAGTGTTTA 3’) was designed and synthesized to sterically bind to the junction region of Intro19 and Exon 20 of *TJP1* gene with chemical modifications (5-methyl-C (C*), phosphorothioate and 2’-O- methoxyethyl group).

*Quantitative RT-PCR (SYBR)*

Quantitative PCR was performed in 20 µL reaction volumes containing 10 µL SYBR Green Master Mix (2×, Applied Biosystems), 0.5 µM each forward/reverse primer (**Table 1**), 4 µL diluted cDNA template (1:10) prepared as above after DNAse digestion, and nuclease-free water. Reactions were run in triplicates on a QuantStudio 3 System (Applied Biosystems) using the following cycling protocol: initial denaturation at 95°C for 10 min; 40 cycles of 95°C for 15 sec (denaturation) and 60°C for 1 min (annealing/extension); followed by melt curve to ensure specific products. Water controls (no template) were included in every experimental run to rule out contamination. The ΔΔCt method [10] was used to normalize with housekeeping gene and to analyze the results as described previously (Bryant et al., 2022).

*Western Blot Analysis*

Differentiated human podocytes (60,000 cells/well) were treated for 48 hours with vehicle (0.1% DMSO), PAN (10 µg/mL or 25 µg/mL), and ADR (5 µg/mL). Cells were harvested and lysed using M-PER™ Reagent (Mammalian Protein Extraction Reagent, Thermo Fisher Scientific), supplemented with protease and phosphatase inhibitors, followed by centrifugation at 12,000 × g for 15 min at 4°C. Protein concentration was determined using the BCA assay. Equal amounts of protein (20–30 µg) were resolved on 8% SDS-PAGE gels, transferred to PVDF membrane using a wet transfer system, and blocked with 5% non-fat milk in TBST for 1 h at room temperature. Membranes were incubated overnight at 4°C with primary antibodies against TJP1 (1:1,000; Thermo Fisher, PA528869), VINCULIN (1:5,000; Thermo Fisher, MA5-16424), GAPDH (1:5,000; Thermo Fisher, 39-8600), and ITM2B (1:1,000; Thermo Fisher,

PA5-31441). After washing with TBST, membranes were incubated with HRP-conjugated secondary antibodies (1:5,000; Thermo Fisher) for 1 h at room temperature, and protein bands were visualized using an Enhanced Chemiluminescence (ECL) Substrate (Millipore) and imaged on an Azure 400 Imaging System. Band intensities were quantified using ImageJ and normalized to housekeeping.

*Immunofluorescence (IF) Staining*

Differentiated podocytes were cultured on 22 mm coverslips (60,000 cells per well in a 6-well plate) and treated for 48 hours with vehicle (0.1% DMSO), PAN (25 μg/mL), or ADR (5 μg/mL). Following treatment, cells were fixed with 4% paraformaldehyde for 30 minutes at room temperature. Blocking was performed using Super Block (Sytek Laboratory, AAA500) supplemented with 0.3% Triton X-100 for 30 minutes, and repeated twice. Cells were then incubated overnight at 4°C with primary antibodies against TJP1 total (1:100; Thermo Fisher, 33-9100), TJP1 α+ form (1:50; OriGene Technologies, AP26402PU), ITM2B total (1:100; Thermo Fisher, PA5-31441), ITM2B Long Form (1:1,000; Santa Cruz, sc-374362), α-TUBULIN (1:10,000; ThermoFisher, A11126) in Super Block containing 0.3% Triton X-100. After three washes with PBST containing 2.5% Sytek blocking buffer, cells were incubated with Alexa Fluor-conjugated secondary antibodies (1:500; Thermo Fisher) for 1 hour at 37°C in dark. Coverslips were mounted using ProLong™ Gold DAPI Antifade Mountant (Thermo Fisher). Fluorescent images were captured using a Nikon Eclipse Ni fluorescence microscope. Rat kidney sections obtained from animal studies were deparaffinized overnight at 64°C in an incubator. Antigen retrieval was performed using the Decloaking Chamber™ NxGen (Biocare Medical, DC2012) with citrate buffer (pH 6.0) at 110°C for 15 min, followed by gradual cooling. Subsequent blocking, staining and microscopy steps were performed as described above.

### Supplementary Figures

#### Supplementary Figure S1

*Serum Chemistry of PAN and Adriamycin-Induced Nephropathy Models*

Serum chemistry profile revealed conditions like hypoalbuminemia and hypercholesterolemia and several other parameters altered in the PAN and adriamycin model of nephropathies.

#### Supplementary Figure S2

*Ontology Enrichment Analysis of Differentially Expressed Genes (DEGs) in PAN and Adriamycin-Induced Nephropathy Models*

Ontology enrichment analysis identified top cellular components (GOCC), biological processes (GOBP), and molecular functions (GOMF) of all the differentially expressed genes (DEGs) in the PAN and adriamycin models **(Ai-iii)** as well as DEGs that were either up- or down-regulated in PAN and adriamycin models **(Bi-iii)**.

#### Supplementary Figure S3

*DEGs in PAN and Adriamycin Models Deconvoluted for Podocytes, other Glomerular Cells and Parietal Epithelial Cell Subtypes*

In order to understand the contribution of all the glomerular cell types in the glomeruli and podocyte specific DEG responses in PAN and adriamycin models of injury, we deconvoluted the RNASeq dataset by first creating a robust podocyte metrics by cross-referencing the DEGs in PAN and adriamycin models with podocyte-specific transcriptomic and proteomic dataset [11] and then with the top 50 most variable genes specifically expressed in these multiple cell types[12]. From the podocyte-specific transcriptomic dataset of 6593 genes (TPM > 10, $q < 0.05$) [11], we identified 1852 and 2534 DEGs (FC > 1, <1, $q < 0.05$) in PAN and adriamycin models, of which 1636 DEGs were common between the two models (**Ai**). From the podocyte-specific iBAQ dataset of 551 proteins ($q < 0.05$) [11], we identified 179 and 199 DEGs (FC > 1, <1, $q < 0.05$) in PAN and adriamycin models, of which 148 DEGs were common between the two models (**Aii**). Podocyte specific genes *Nphs2, Rhpn1, Tmsb4x, Rasl11a, Adm, Cryab, Htra1*, were identified to be altered in both disease models, underlining their crucial role in the podocyte pathophysiology in these disease models (B**i-iii**). Mesangial specific genes *Tpm2, Mgp, Gadd45b, Emd, Map1lc3a,* were altered in both disease models, with additional genes like *Grg11* and *Map3k7cl* being altered uniquely in the adriamycin model. In both PAN and adriamycin models, endothelial specific genes *Cdkn1a, Crip1, Clic1, Ifitm3* were consistently altered. However, adriamycin showed an additional alteration in genes *Ctla2a, Cyp4b1, Emp1, Fxyd5, Igfbp5, Pbx1, B2m*, indicating a more extensive endothelial response in this model. Immune cell specific genes, *Rps4x, Emp3, Ctsz, Tyrobp, Fcer1g, Cd83, Gm2a, Rpl32 were altered* in both disease models, suggesting the involvement of immune responses in both PAN and adriamycin. However, *Rgs2, B2m* and *Msn* were uniquely altered in adriamycin, indicating specific contribution of immune response to adriamycin model. Furthermore, with the recent advancements in the diverse roles of parietal epithelial cell sub-types in the regeneration of podocytes and pathophysiology of glomerular injury, we further deconvoluted the RNASeq data with significant DEGs from these models with the variety of parietal epithelial cell sub-types [13]. A comprehensive cross-referencing with the genes associated with five distinct parietal epithelial cell subpopulations, A1 (136 genes), A2 (39 genes), A3 (90 genes), A4 (277 genes), B (221 genes) identified a multitude of DEGs represented from all the five subpopulations, with the most altered DEGs in the podocyte progenitor parietal epithelial cell-A1 subpopulation. especially in the adriamycin model (C**i and ii, Supplementary Table 1**). In summary, our study suggests a complex interplay and molecular contributions of different glomerular compartments in the PAN and adriamycin injury models.

**Supplementary Figure S4**

*DEGs in PAN and Adriamycin Models Represented in Established Genes Associated with Glomerular Disease and Podocytopathies*

In order to correlate the DEGs identified in the PAN and adriamycin glomerular injury models with the glomerular disease in human patients, we cross-referenced our dataset (*P*adj. <0.05, FC>1, <1) with 80 glomerular genes with established monogenic roles in glomerular disease [14]and 49 genes associated with podocytopathies and steroid resistant nephrotic syndrome [15]. We identified 28 genes DEGs in both PAN and adriamycin models that have established monogenic roles in glomerular disease. as depicted in the heatmaps (**Ai, significant DEGs in PAN; Aii, significant DEGs in Adriamycin, ADR**). These mostly localized within the podocytes (membrane, cytoskeletal scaffold, nuclear proteins and transcription factors, slit diaphragm and mitochondria), endothelial cells or glomerular basement membrane (**Aiii**). Furthermore, upon comparison with genes associated with steroid resistant nephrotic syndrome, we identified 23 and 27 DEGs in PAN and adriamycin models, respectively, with a significant overlap between the two models (**Bi, significant DEGs in adriamycin, ADR**). These are involved in various podocyte functions, such as actin binding and regulation, slit membrane, S1P metabolism, tRNA modification, CoQ biosynthesis and nucleoporins (**Bii**). In summary, our data on glomerular DEGs altered in the two disease models was corroborated with the human glomerular disease data on these overlapping genes.

**Supplementary Figure S5**

*Dispersion Plots Depicted Good Fit for the PAN and Adriamycin models for Exon/Intron/Junction Analyses*

**Supplementary Figure S6**

*Heatmap of DEGs in the PAN and Adriamycin (ADR) Models Represented in the Dataset of a Combination of Transcription and Translation Machinery Factors* (compiled for this study). Evolving research shows that the processes of mRNA transcription and processing via splicing and polyadenylation, and even translation can be intertwined events[16, 17]. Analysis of our glomerular disease DEGs dataset identified several factors, such as RNA polymerase II associated factors, DNA damage response factors, and translation factors that were dysregulated in both PAN (24) and adriamycin (28) models **(Supplementary TableS14)**. **(i)** Cross-referenced with the

DEGs (adj $p$ <0.05, FC>1, <1) in the PAN RNASeq dataset, **(ii)** Cross-referenced with the DEGs (adj $p$ <0.05, FC>1, <1) in the adriamycin (ADR) RNASeq dataset.

**Supplementary Figure S7**

*Manhattan Plots Depicting Multi-Population GWAS Signals/SNPs Localized in the Regulatory Regions of Alternatively Spliced Genes and in the 3'UTR of Alternatively Polyadenylated Genes in the PAN Model.*

Evolving genome wide association studies (GWAS) has identified numerous genetic determinants or single nucleotide polymorphisms (SNPs) utilizing multi-population glomerular disease and healthy cohorts[8, 18]. Most of these genetic variants are noncoding, indicating their potential role in gene regulation. In order to understand their functional and clinical consequences, we cross-referenced the GWAS signals from multi-population study (38,463 participants, including 2440 cases) of SNPs associated with steroid sensitive nephrotic syndrome [8] with the genomic region of 136 alternatively spliced (AS) genes and 3' UTRs of 71 alternatively polyadenylated (APA) genes in the PAN-minimal change disease model dataset from this study. (**A**) Genomic regions (including introns, exons, 5'UTR, 3'UTR, and splice sit junctions) in the 136 alternatively spliced genes identified a few significant SNPs (e.g. *EHMT2, PRRC2A, NPHS1*), and several other SNPs of low significance (e.g. *TJP1, ITM2B, PODXL, MYO1D, COL4A4, NPHS2*). (**B**) 3' UTR regions in the 71 alternatively genes identified a few SNPs of low significance (e.g. *NPHS1, PODXL*). These regions were defined by the grch37 to match the GWAS version. Genes of interest are marked with an arrow. SNPs threshold; High Significance: -log10(P) value of 7.3 = $5X10^{-8}$, above the red line; Low significance: -log10(P) value 1.3 = $5X10^{-2}$, above the blue line. These data suggest the possibility that the SNPs associated with nephrotic syndrome may serve as potential cis-elements for alternative mRNA processing.

**Supplementary Figure S8**

*Genomic Organization of Neph1, Nphs1 and Nphs2 Genes and 3'UTR, Poly (A) Site Usage Identified, RACE and 3'PCR Primers Used and Sequence Chromatograms Obtained from RACE Assays.*

#### Supplementary Tables

**Supplementary Table 1**

Deconvolution of DEGs in PAN and adriamycin models for parietal epithelial cell subtypes

**Supplementary Table 2**

Deconvolution of alternatively spliced (AS) genes in PAN and adriamycin models for parietal epithelial cell subtypes

**Supplementary Table 3**

Alternatively spliced (AS) genes in PAN and adriamycin models represented in monogenic genes with established roles in glomerular disease

**Supplementary Table 4**

Alternatively spliced (AS) genes in PAN and adriamycin models representing novel genes in glomerular disease

**Supplementary Table 5**

Alternatively spliced (AS) genes in PAN model reversed with pioglitazone treatment

**Supplementary Table 6**

Alternatively spliced (AS) genes in PAN model reversed with GQ-16 treatment

**Supplementary Table 7**

Deconvolution of alternatively polyadenylated (APA) genes in PAN and adriamycin models for parietal epithelial cell subtypes

**Supplementary Table 8**

Alternatively polyadenylated (APA) genes in PAN and adriamycin models represented in monogenic genes with established roles in glomerular disease

**Supplementary Table 9**

Alternatively polyadenylated (APA) genes in PAN and adriamycin models representing novel genes in glomerular disease

**Supplementary Table 10**

Alternatively polyadenylated (APA) genes in PAN model reversed with pioglitazone treatment

**Supplementary Table 11**

Alternatively polyadenylated (APA) genes in PAN model reversed with GQ-16 treatment

**Supplementary Table 12**

Dysregulated RBPs and splicing factors in PAN and adriamycin models

**Supplementary Table 13**

Dysregulated pre-mRNA 3’ processing and polyadenylation factors in PAN and adriamycin models

**Supplementary Table 14**

Dysregulated transcription and translation machinery factors in PAN and adriamycin models

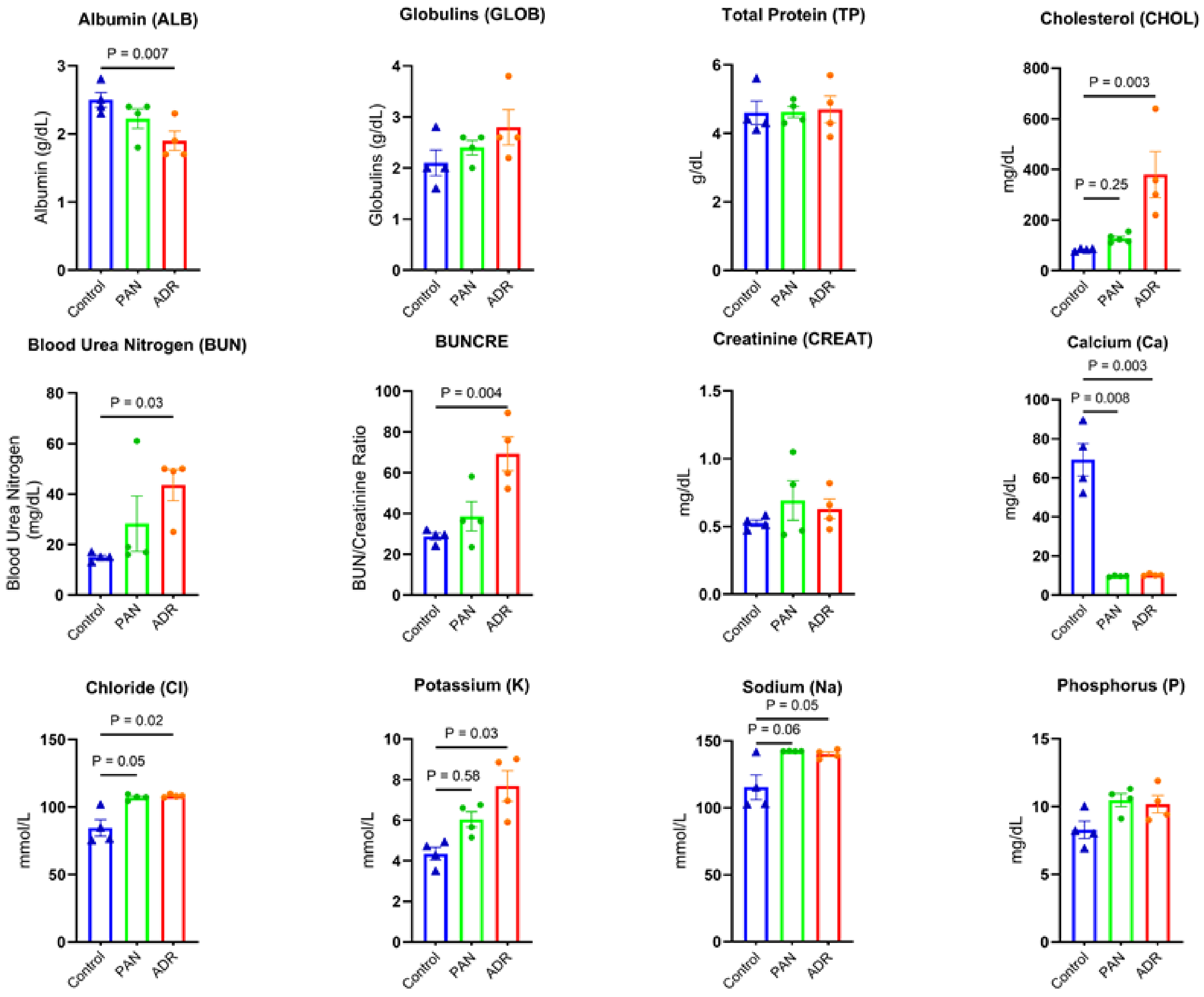

Serum Chemistry of PAN and Adriamycin (ADR)-Induced Nephropathy Models

**Supplementary Figure S1**

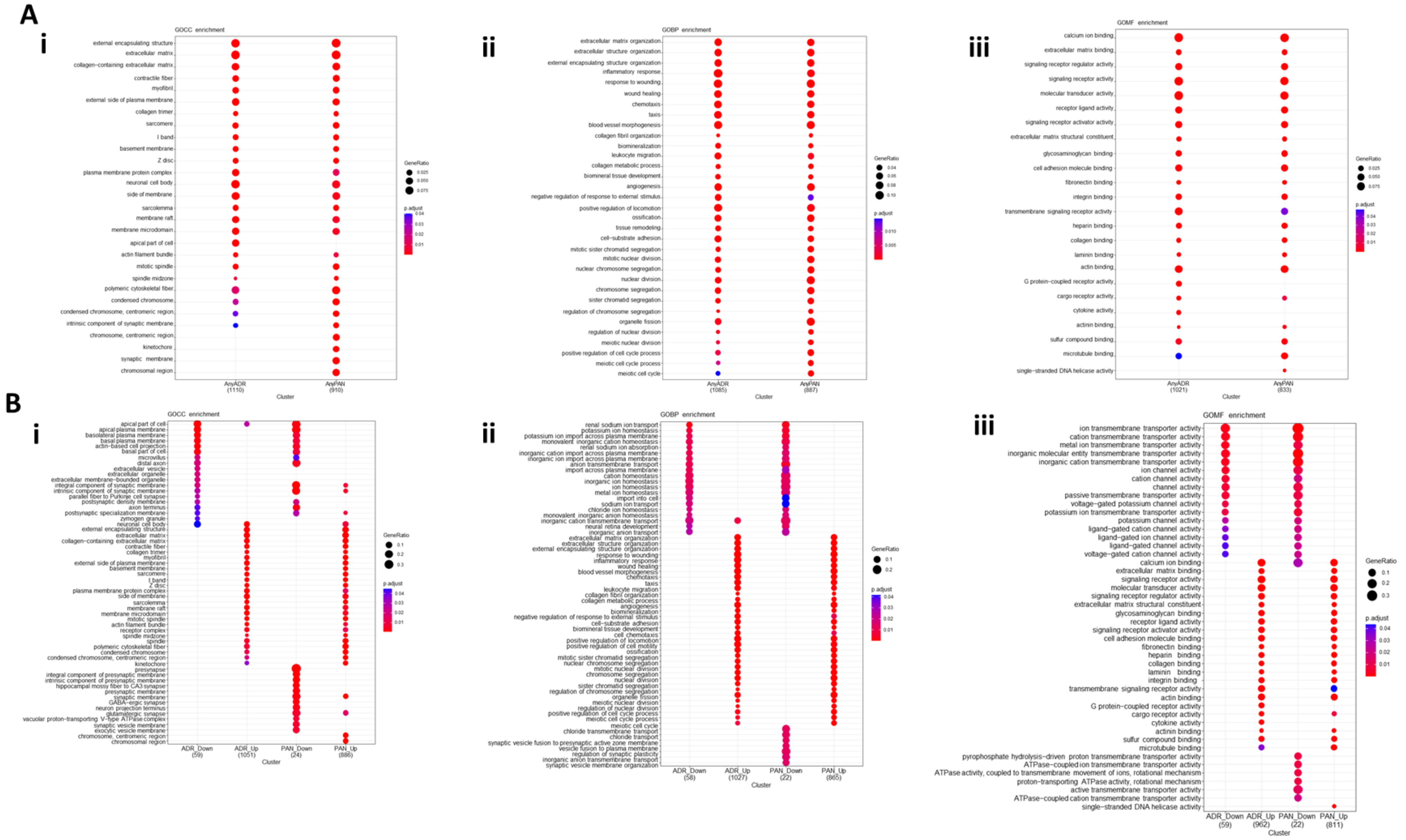

Ontology Enrichment Analysis of Differentially Expressed Genes (DEGs) in PAN and Adriamycin-Induced Nephropathy Models

**Supplementary Figure S2**

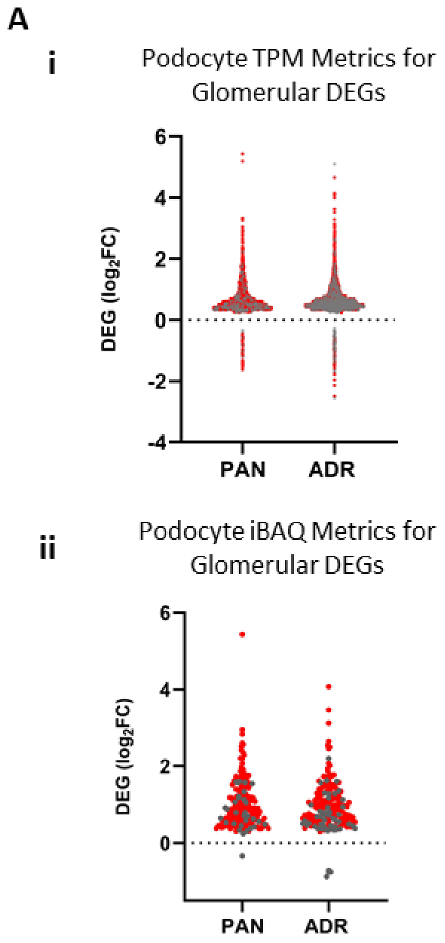

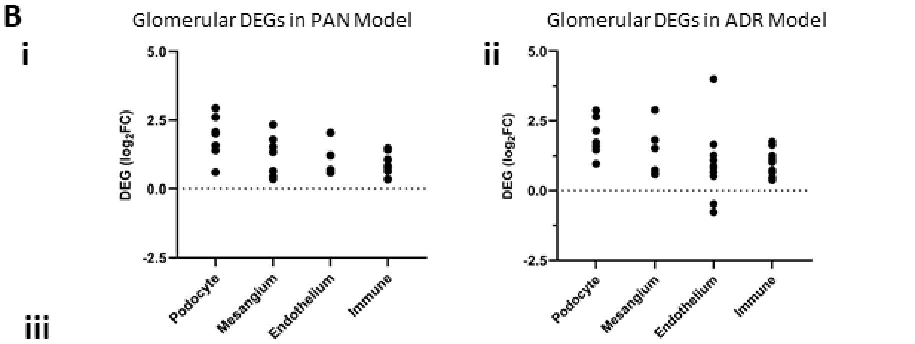

iii

**Deconvolution of DEGs in PAN and ADR Models for Glomerular Cell Population**

| Cell types | PAN | ADR |
|---|---|---|
| **Podocyte** | *Nphs2, Rhpn1, Tmsb4x, Ras111a, Cryab, Adm, Htra1* | *Nphs2, Rhpn1, Tmsb4x, Ras111a, Cryab, Adm, Htra1* |
| **Mesangium** | *Emd, Gadd45b, Map11c3a, Mgp, Tpm2, Usp2* | *Emd, Gadd45b, Gng11, Map11c3a, Map3k7cl, Mgp, Tpm2* |
| **Endothelium** | *Cdkn1a, Crip1, Clic1, Ifitm3* | *Cdkn1a, Crip1, Clic1, Ifitm3, Ctla2a, Cyp4b1, Emp1, Fxyd5, Igfbp5, Pbx1, B2m,* |
| **Immune** | *Rps4x, Emp3, Ctsz, Tyrobp, Fcer1g, Cd83, Gm2a, Rpl32* | *Rps4x, Emp3, Ctsz, Tyrobp, Fcer1g, Cd83, Gm2a, Rpl32, Rgs2, Msn, B2m* |

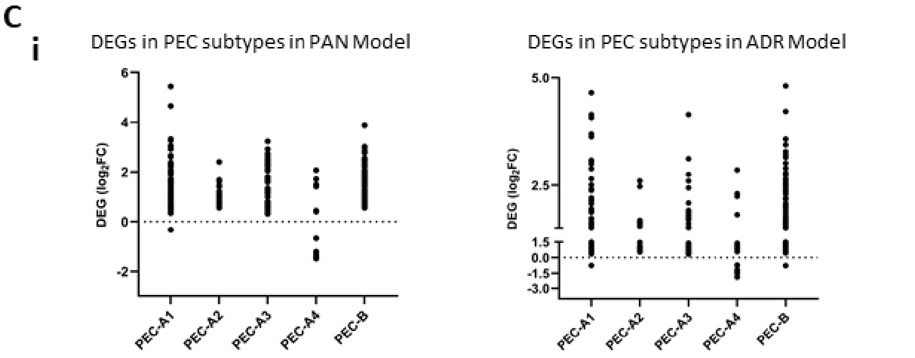

DEGs in PAN and Adriamycin Models Deconvoluted for Podocytes, other Glomerular Cells and Parietal Epithelial Cell Subtypes

**Supplementary Figure S3**

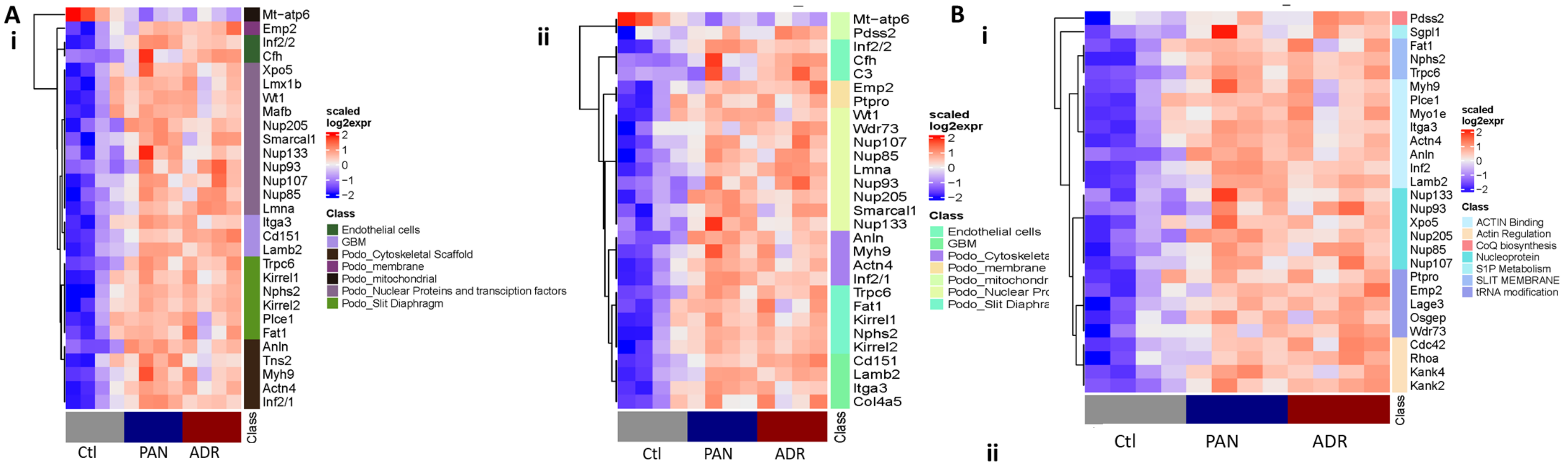

iii

| Monogenic Glomerular Disease Genes Represented in the DEGs in PAN and ADR Models | | | |
|---|---|---|---|
| **Class** | **Glomerulus** | **PAN (MCD)** | **ADR (FSGS)** |
| Podocytes | Membrane | *Emp2* | *Emp2, Ptpro* |
| | Cytoskeletal Scaffold | *Actn4, Anln, Inf2, Myh9, Tns2* | *Actn4, Anln, Inf2, Myh9* |
| | Nucleoporins and transcription factors | *Lmna, Nup107, Nup133, Nup205, Nup85, Nup93, Smarcal1, Wt1, Lmx1b, Mafb, Xpo5* | *Lmna, Nup107, Nup133, Nup205, Nup85, Nup93, Smarcal1, Wt1, Wdr73* |
| | Slit Diaphragm | *Fat1, Kirrel1, Kirrel2, Nphs2 Trpc6, Plce1* | *Fat1, Kirrel1, Kirrel2, Nphs2 Trpc6* |
| | Mitochondrial | *Mt-Atp6* | *Mt-Atp6, Pdss2* |
| **Endothelial** | Endothelial cells | *Cfh, Inf2* | *C3, Cfh, Inf2* |
| **Matrix** | GBM | *Itga3, Lamb2, Cd151* | *Itga3, Lamb2, Cd151, Col4a5* |

ii

| Podocyte Genes Associated with Steroid Resistance Represented in the DEGs in PAN and ADR Models | | |
|---|---|---|
| **Class** | **PAN (MCD)** | **ADR (FSGS)** |
| Actin Binding | *Actn4, Anln, Inf2, Itga3, Plcel, Lamb2, Myh9, Myo1e* | *Actn4, Anln, Inf2, Itga3, Itgb4, Lamb2, Myh9, Myo1e* |
| Slit membrane | *Fat1, Trpc6, Nphs2* | *Fat1, Trpc6, Nphs2* |
| S1P Metabolism | *Sgpl1* | *Sgpl1* |
| tRNA modification | *Emp2, Lage3, Osgep* | *Emp2, Lage3, Osgep, Ptpro, Wdr73* |
| Actin Regulation | *Kank2, Kank4* | *Kank2, Kank4, Cdc42, Rhoa* |
| CoQ biosynthesis | | *Pdss2* |
| Nucleoporin | *Nup107, Nup133, Nup205, Nup85, Nup93, Xpo5* | *Nup107, Nup133, Nup205, Nup85, Nup93* |

DEGs in PAN and Adriamycin Models Represented in Established Genes Associated with Glomerular Disease and Podocytopathies

**Supplementary Figure S4**

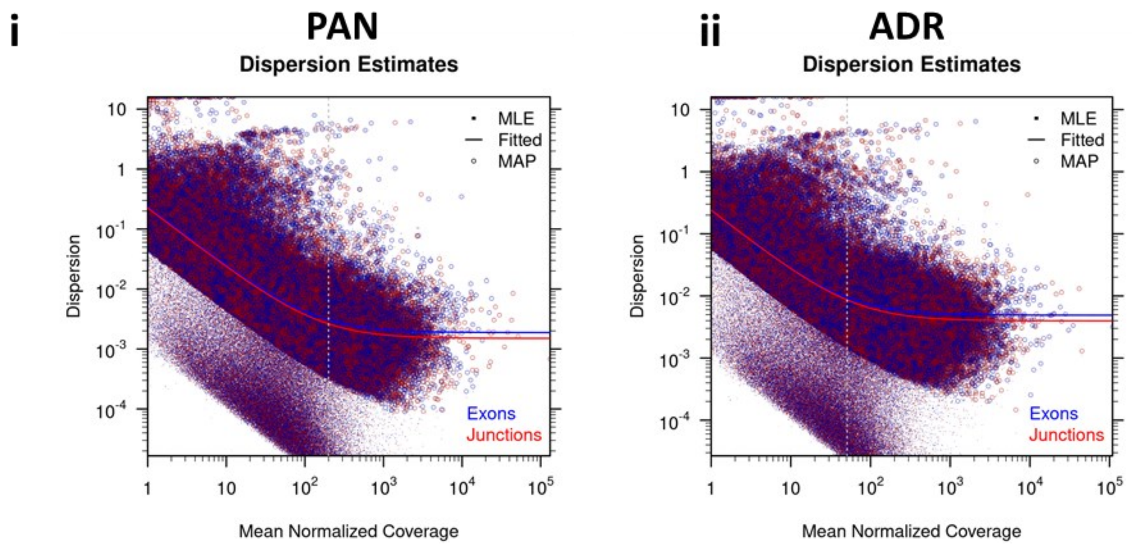

Dispersion Plots Depicted Good Fit for the PAN and Adriamycin (ADR) Models for Exon/Intron/Junction Analyses

**Supplementary Figure S5**

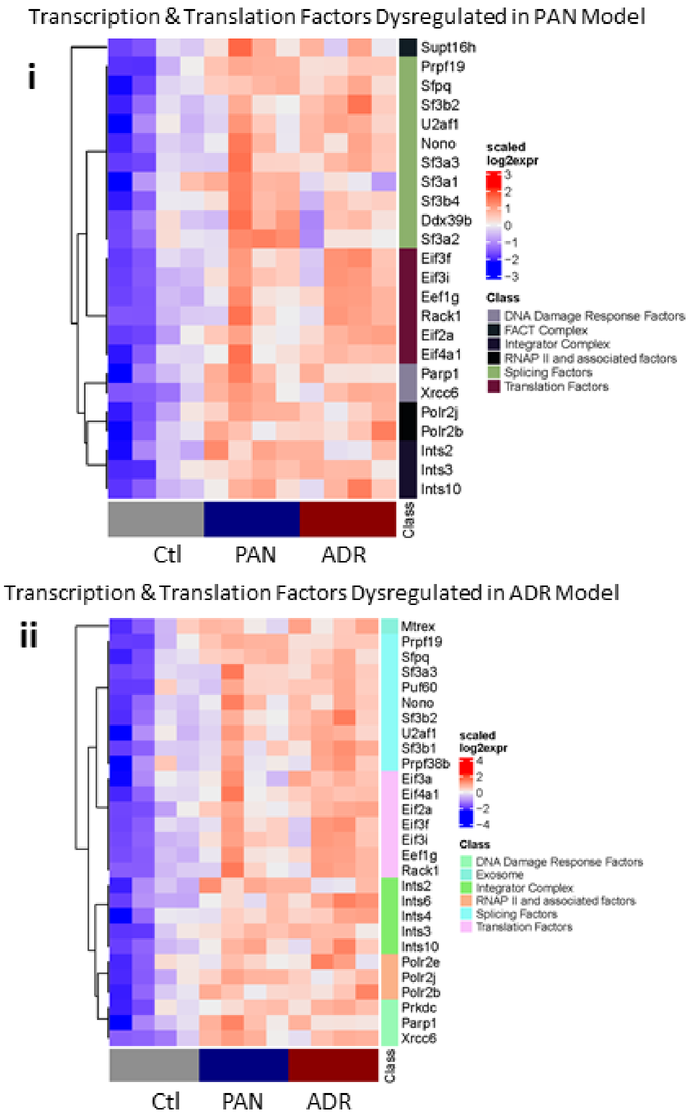

Heatmap of DEGs in the PAN and Adriamycin (ADR) Models Represented in the Dataset of a Combination of Transcription and Translation Machinery Factors

**Supplementary Figure S6**

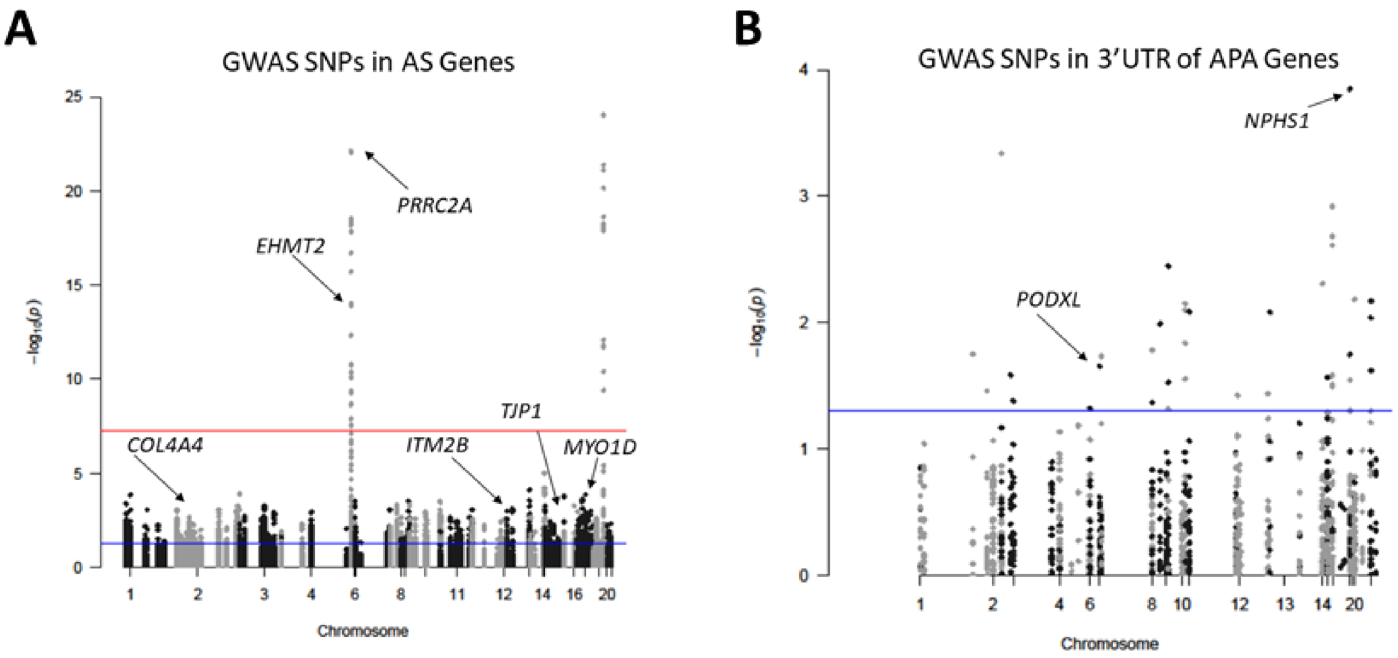

Manhattan Plots Depicting Multi-Population GWAS Signals/SNPs Localized in the Regulatory Regions of AS Genes and in the 3'UTR of APA Genes in the PAN Model

**Supplementary Figure S7**

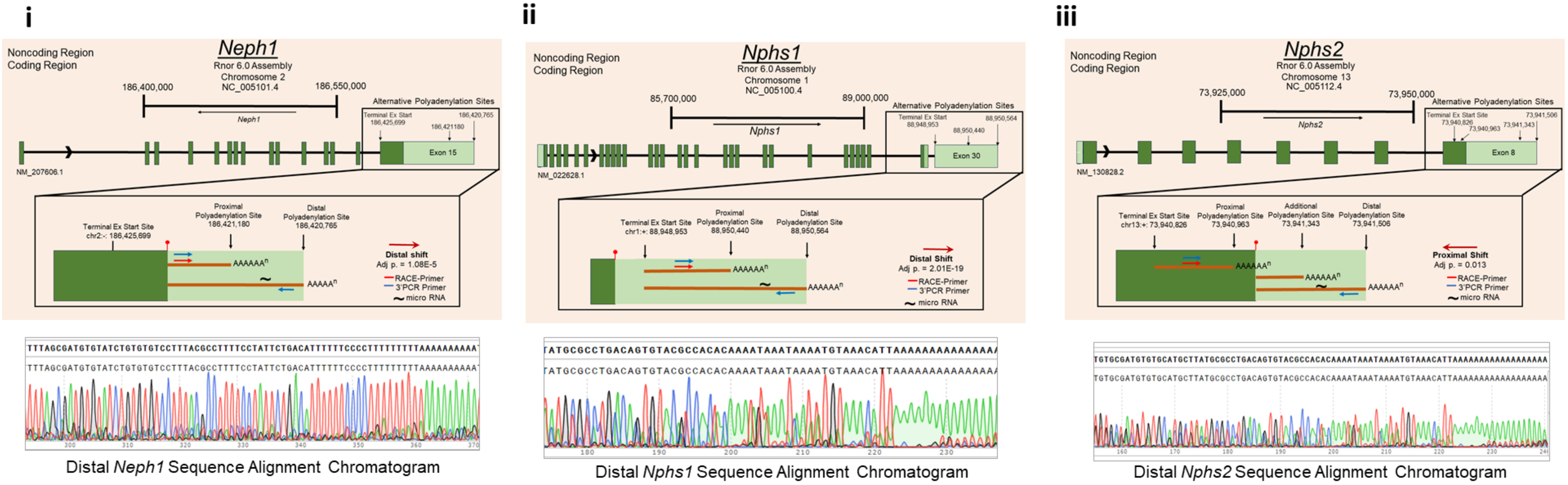

Genomic Organization of *Neph1, Nphs1 and Nphs2* Genes and 3'UTR, Poly (A) Site Usage Identified, RACE and 3'PCR Primers Used and Sequence Chromatograms Obtained from RACE Assays

**Supplementary Figure S8**

Supplementary Table 1: Deconvolution of DEGs in PAN and ADR models for PEC subtypes

| | | PAN | |
|---|---|---|---|
| **Cluster** | **Gene** | **logFC PANvsCtl** | **adj.P.Val PANvsCtl** |
| PEC-A1 | *Ackr3* | 1.54 | 0.04 |
| PEC-A1 | *Actb* | 0.79 | 0.00 |
| PEC-A1 | *Actg1* | 0.45 | 0.01 |
| PEC-A1 | *Actn4* | 1.11 | 0.01 |
| PEC-A1 | *Adm* | 2.94 | 0.00 |
| PEC-A1 | *Angptl2* | 2.21 | 0.01 |
| PEC-A1 | *Anxa1* | 1.31 | 0.03 |
| PEC-A1 | *Cavin1* | 0.95 | 0.00 |
| PEC-A1 | *Cavin3* | 0.59 | 0.01 |
| PEC-A1 | *Ccn1* | 1.59 | 0.00 |
| PEC-A1 | *Ccn2* | 2.38 | 0.01 |
| PEC-A1 | *Cd151* | 0.99 | 0.00 |
| PEC-A1 | *Cd82* | 0.65 | 0.01 |
| PEC-A1 | *Chst1* | 1.14 | 0.04 |
| PEC-A1 | *Cmtm7* | 1.18 | 0.00 |
| PEC-A1 | *Cryab* | 2.01 | 0.01 |
| PEC-A1 | *Des* | 3.27 | 0.00 |
| PEC-A1 | *Dusp3* | 0.43 | 0.04 |
| PEC-A1 | *Efnb1* | 0.92 | 0.04 |
| PEC-A1 | *Eif3m* | 0.77 | 0.01 |
| PEC-A1 | *Eml2* | 0.60 | 0.03 |
| PEC-A1 | *Enpep* | 2.36 | 0.00 |
| PEC-A1 | *Epn3* | 1.60 | 0.02 |
| PEC-A1 | *Fgl2* | 1.61 | 0.00 |
| PEC-A1 | *Gadd45a* | 1.51 | 0.00 |
| PEC-A1 | *Gadd45b* | 1.34 | 0.03 |
| PEC-A1 | *Golim4* | 0.95 | 0.05 |
| PEC-A1 | *Hs3st6* | 2.25 | 0.01 |
| PEC-A1 | *Htra1* | 0.61 | 0.04 |
| PEC-A1 | *Itgb1* | 1.08 | 0.01 |
| PEC-A1 | *Loxl2* | 2.66 | 0.00 |
| PEC-A1 | *Lrrfip1* | 1.09 | 0.00 |
| PEC-A1 | *Lsp1* | 0.68 | 0.05 |
| PEC-A1 | *Mafb* | 1.40 | 0.04 |
| PEC-A1 | *Metrnl* | 1.71 | 0.03 |
| PEC-A1 | *Mustn1* | 1.59 | 0.03 |
| PEC-A1 | *Myl12a* | 1.03 | 0.00 |
| PEC-A1 | *Myl6* | 0.77 | 0.00 |
| PEC-A1 | *Myom2* | 5.44 | 0.00 |
| PEC-A1 | *Nes* | 2.30 | 0.01 |
| PEC-A1 | *Nphs2* | 2.61 | 0.01 |
| PEC-A1 | *Npnt* | 1.58 | 0.04 |
| PEC-A1 | *Nupr1* | 1.90 | 0.01 |
| PEC-A1 | *P3h2* | 1.30 | 0.03 |
| PEC-A1 | *Pak1* | 1.43 | 0.03 |
| PEC-A1 | *Pamr1* | 2.00 | 0.01 |
| PEC-A1 | *Pla2g7* | 3.05 | 0.00 |
| PEC-A1 | *Prss23* | 0.62 | 0.05 |
| PEC-A1 | *Rasl11a* | 2.07 | 0.00 |
| PEC-A1 | *Rhpn1* | 1.58 | 0.00 |
| PEC-A1 | *Schip1* | 0.69 | 0.03 |
| PEC-A1 | *Serpine1* | 2.92 | 0.00 |
| PEC-A1 | *Slc48a1* | 1.92 | 0.01 |
| PEC-A1 | *St3gal6* | -0.33 | 0.04 |
| PEC-A1 | *Tagln* | 3.32 | 0.00 |
| PEC-A1 | *Tmem189* | 0.73 | 0.02 |
| PEC-A1 | *Tmsb4x* | 1.41 | 0.00 |
| PEC-A1 | *Tnfrsf12a* | 2.33 | 0.00 |
| PEC-A1 | *Tnfsf15* | 2.01 | 0.01 |
| PEC-A1 | *Tob1* | 0.66 | 0.01 |
| PEC-A1 | *Vamp5* | 0.57 | 0.03 |
| PEC-A1 | *Vim* | 1.67 | 0.00 |
| PEC-A1 | *Wt1* | 1.53 | 0.03 |

| | | ADR | |
|---|---|---|---|
| **Cluster** | **Gene** | **logFC ADRvsCtl** | **adj.P.Val ADRvsCtl** |
| PEC-A1 | *Ackr3* | 2.1513 | 0.0060 |
| PEC-A1 | *Actb* | 0.5534 | 0.0075 |
| PEC-A1 | *Actg1* | 0.3842 | 0.0138 |
| PEC-A1 | *Actn4* | 1.0159 | 0.0093 |
| PEC-A1 | *Adm* | 2.8843 | 0.0003 |
| PEC-A1 | *Angptl2* | 1.5066 | 0.0239 |
| PEC-A1 | *Anxa1* | 1.8698 | 0.0038 |
| PEC-A1 | *Anxa3* | 1.1892 | 0.0002 |
| PEC-A1 | *Aplp1* | 1.2137 | 0.0046 |
| PEC-A1 | *Bcam* | 1.1351 | 0.0247 |
| PEC-A1 | *Cavin1* | 1.2842 | 0.0004 |
| PEC-A1 | *Cavin3* | 0.8607 | 0.0004 |
| PEC-A1 | *Ccn1* | 2.3776 | 0.0000 |
| PEC-A1 | *Ccn2* | 3.0817 | 0.0010 |
| PEC-A1 | *Cd151* | 1.4048 | 0.0001 |
| PEC-A1 | *Cd82* | 0.7671 | 0.0021 |
| PEC-A1 | *Cdkn1c* | 2.2040 | 0.0110 |
| PEC-A1 | *Cmtm7* | 1.0852 | 0.0004 |
| PEC-A1 | *Cryab* | 2.1365 | 0.0025 |
| PEC-A1 | *Des* | 3.6260 | 0.0000 |
| PEC-A1 | *Dusp3* | 0.4564 | 0.0219 |
| PEC-A1 | *Efnb1* | 0.9874 | 0.0259 |
| PEC-A1 | *Eif3m* | 0.8591 | 0.0044 |
| PEC-A1 | *Eml2* | 0.6191 | 0.0185 |
| PEC-A1 | *Enpep* | 2.5053 | 0.0019 |
| PEC-A1 | *Fgl2* | 1.4962 | 0.0008 |
| PEC-A1 | *Fos* | 1.7295 | 0.0008 |
| PEC-A1 | *Gadd45a* | 1.9106 | 0.0004 |
| PEC-A1 | *Gadd45b* | 1.5182 | 0.0099 |
| PEC-A1 | *Golim4* | 1.0134 | 0.0300 |
| PEC-A1 | *Hs3st6* | 2.1445 | 0.0048 |
| PEC-A1 | *Hspb1* | 0.7264 | 0.0121 |
| PEC-A1 | *Htra1* | 0.9650 | 0.0031 |
| PEC-A1 | *Ier2* | 0.6024 | 0.0060 |
| PEC-A1 | *Igfbp2* | 1.9119 | 0.0346 |
| PEC-A1 | *Itgb1* | 1.3086 | 0.0016 |
| PEC-A1 | *Loxl2* | 2.4111 | 0.0006 |
| PEC-A1 | *Lrrfip1* | 1.0267 | 0.0024 |
| PEC-A1 | *Lsp1* | 1.0013 | 0.0053 |
| PEC-A1 | *Mertk* | 1.3012 | 0.0361 |
| PEC-A1 | *Metrnl* | 1.5676 | 0.0310 |
| PEC-A1 | *Mustn1* | 2.3815 | 0.0023 |
| PEC-A1 | *Myl12a* | 1.3164 | 0.0002 |
| PEC-A1 | *Myl6* | 1.0257 | 0.0005 |
| PEC-A1 | *Myom2* | 4.0759 | 0.0034 |
| PEC-A1 | *Nebl* | 1.1316 | 0.0367 |
| PEC-A1 | *Nes* | 2.0728 | 0.0101 |
| PEC-A1 | *Nphs2* | 2.6483 | 0.0044 |
| PEC-A1 | *Nupr1* | 2.9988 | 0.0003 |
| PEC-A1 | *Pak1* | 1.4044 | 0.0227 |
| PEC-A1 | *Pamr1* | 3.6903 | 0.0001 |
| PEC-A1 | *Pla2g7* | 2.1575 | 0.0042 |
| PEC-A1 | *Plat* | 0.6914 | 0.0392 |
| PEC-A1 | *Plod2* | 1.2929 | 0.0308 |
| PEC-A1 | *Plxdc2* | 1.0753 | 0.0483 |
| PEC-A1 | *Prss23* | 1.9109 | 0.0000 |
| PEC-A1 | *Ptpro* | 1.6254 | 0.0485 |
| PEC-A1 | *Rasl11a* | 1.7270 | 0.0034 |
| PEC-A1 | *Rhpn1* | 1.4643 | 0.0037 |
| PEC-A1 | *Schip1* | 0.7230 | 0.0184 |
| PEC-A1 | *Sdc4* | 1.2823 | 0.0143 |
| PEC-A1 | *Serpinb6b* | -0.7597 | 0.0307 |
| PEC-A1 | *Serpine1* | 4.1478 | 0.0000 |
| PEC-A1 | *Slc31a2* | 1.1045 | 0.0221 |
| PEC-A1 | *Slc48a1* | 1.6465 | 0.0166 |
| PEC-A1 | *Tagln* | 4.6535 | 0.0000 |
| PEC-A1 | *Tmem189* | 0.6585 | 0.0174 |
| PEC-A1 | *Tmod3* | 0.3429 | 0.0436 |
| PEC-A1 | *Tmsb4x* | 1.5954 | 0.0007 |

| | | | |
|---|---|---|---|
| PEC-A1 | *Tnfrsf12a* | 3.0587 | 0.0001 |
| PEC-A1 | *Tnfsf15* | 1.9499 | 0.0072 |
| PEC-A1 | *Tob1* | 0.8719 | 0.0007 |
| PEC-A1 | *Trib2* | 0.5415 | 0.0420 |
| PEC-A1 | *Vamp5* | 0.5056 | 0.0315 |
| PEC-A1 | *Vim* | 1.7217 | 0.0014 |
| PEC-A1 | *Wt1* | 1.2312 | 0.0452 |

| Cluster | Gene | logFC PANvsCtl | adj.P.Val PANvsCtl |
|---|---|---|---|
| PEC-A2 | *Adamts1* | 0.73 | 0.02 |
| PEC-A2 | *Ankrd1* | 0.82 | 0.01 |
| PEC-A2 | *Stc2* | 1.41 | 0.03 |
| PEC-A2 | *Rasl11b* | 0.82 | 0.03 |
| PEC-A2 | *Akap12* | 1.62 | 0.00 |
| PEC-A2 | *Ncam1* | 2.40 | 0.01 |
| PEC-A2 | *Cxcl10* | 0.56 | 0.04 |
| PEC-A2 | *Atf3* | 0.68 | 0.05 |
| PEC-A2 | *Tinagl1* | 1.10 | 0.01 |
| PEC-A2 | *Gstm1* | 1.21 | 0.02 |
| PEC-A2 | *Lsp1* | 1.08 | 0.04 |
| PEC-A2 | *Cpe* | 1.45 | 0.00 |

| Cluster | Gene | logFC ADRvsCtl | adj.P.Val ADRvsCtl |
|---|---|---|---|
| PEC-A2 | *Adamts1* | 0.9035 | 0.0038 |
| PEC-A2 | *Akap12* | 0.8851 | 0.0027 |
| PEC-A2 | *Ankrd1* | 1.5456 | 0.0145 |
| PEC-A2 | *Atf3* | 1.5730 | 0.0004 |
| PEC-A2 | *Bcam* | 1.1351 | 0.0247 |
| PEC-A2 | *C3* | 1.6433 | 0.0059 |
| PEC-A2 | *Clu* | 1.4640 | 0.0020 |
| PEC-A2 | *Cp* | 2.5989 | 0.0011 |
| PEC-A2 | *Cpe* | 1.6101 | 0.0005 |
| PEC-A2 | *Cxcl10* | 2.4712 | 0.0077 |
| PEC-A2 | *Gstm1* | 0.7720 | 0.0051 |
| PEC-A2 | *Insig2* | 0.5454 | 0.0369 |
| PEC-A2 | *Lsp1* | 1.0013 | 0.0053 |
| PEC-A2 | *Ncam1* | 0.9241 | 0.0148 |
| PEC-A2 | *Rasl11b* | 1.6800 | 0.0028 |
| PEC-A2 | *Stc2* | 0.9574 | 0.0499 |
| PEC-A2 | *Tinagl1* | 1.6302 | 0.0001 |
| PEC-A2 | *Tsc22d1* | 0.5821 | 0.0025 |
| PEC-A2 | *Wsb1* | 0.7395 | 0.0011 |

| Cluster | Gene | logFC PANvsCtl | adj.P.Val PANvsCtl |
|---|---|---|---|
| PEC-A3 | *Anxa2* | 0.74 | 0.00 |
| PEC-A3 | *Birc5* | 1.74 | 0.00 |
| PEC-A3 | *Ccdc34* | 0.81 | 0.03 |
| PEC-A3 | *Ccna2* | 2.44 | 0.00 |
| PEC-A3 | *Ccnb1* | 1.80 | 0.01 |
| PEC-A3 | *Cd44* | 1.00 | 0.02 |
| PEC-A3 | *Cd9* | 1.27 | 0.01 |
| PEC-A3 | *Cdc20* | 2.13 | 0.00 |
| PEC-A3 | *Cdca3* | 2.29 | 0.00 |
| PEC-A3 | *Cdca8* | 1.34 | 0.00 |
| PEC-A3 | *Cdk1* | 2.74 | 0.00 |
| PEC-A3 | *Cdkn3* | 2.39 | 0.00 |
| PEC-A3 | *Cenpa* | 2.05 | 0.00 |
| PEC-A3 | *Cks1b* | 1.61 | 0.00 |
| PEC-A3 | *Cks2* | 1.28 | 0.01 |
| PEC-A3 | *Dut* | 0.52 | 0.01 |
| PEC-A3 | *Hmgb2* | 1.24 | 0.00 |
| PEC-A3 | *Hmgn2* | 0.51 | 0.02 |
| PEC-A3 | *Hmmr* | 1.59 | 0.00 |
| PEC-A3 | *Jpt1* | 0.53 | 0.02 |
| PEC-A3 | *Lgals1* | 2.07 | 0.00 |
| PEC-A3 | *Lgals3* | 1.66 | 0.03 |
| PEC-A3 | *Lsm2* | 0.68 | 0.01 |
| PEC-A3 | *Lsm5* | 0.57 | 0.02 |
| PEC-A3 | *Nhp2* | 0.80 | 0.01 |
| PEC-A3 | *Nme1* | 0.56 | 0.02 |
| PEC-A3 | *Npm1* | 0.35 | 0.04 |
| PEC-A3 | *Pbk* | 3.23 | 0.00 |
| PEC-A3 | *Pcna* | 0.76 | 0.00 |
| PEC-A3 | *Pkm* | 0.81 | 0.01 |
| PEC-A3 | *Ppp1r14b* | 0.67 | 0.00 |
| PEC-A3 | *Prc1* | 2.52 | 0.00 |
| PEC-A3 | *Racgap1* | 2.29 | 0.00 |
| PEC-A3 | *Ran* | 0.51 | 0.00 |
| PEC-A3 | *Ranbp1* | 0.45 | 0.01 |
| PEC-A3 | *Rpl12* | 0.34 | 0.05 |
| PEC-A3 | *Rps2* | 0.31 | 0.04 |
| PEC-A3 | *Rrm2* | 1.17 | 0.01 |
| PEC-A3 | *S100a10* | 1.14 | 0.01 |
| PEC-A3 | *S100a6* | 2.24 | 0.01 |
| PEC-A3 | *Selenoh* | 0.33 | 0.05 |
| PEC-A3 | *Serpine1* | 2.92 | 0.00 |

| Cluster | Gene | logFC ADRvsCtl | adj.P.Val ADRvsCtl |
|---|---|---|---|
| PEC-A3 | *Anxa2* | 1.0524 | 0.0001 |
| PEC-A3 | *Ccdc34* | 1.0830 | 0.0048 |
| PEC-A3 | *Ccna2* | 1.7465 | 0.0047 |
| PEC-A3 | *Cd44* | 1.7740 | 0.0003 |
| PEC-A3 | *Cd9* | 1.5214 | 0.0034 |
| PEC-A3 | *Cdc20* | 1.3561 | 0.0073 |
| PEC-A3 | *Cdca3* | 1.7300 | 0.0143 |
| PEC-A3 | *Cdca8* | 0.9745 | 0.0076 |
| PEC-A3 | *Cdk1* | 2.0886 | 0.0033 |
| PEC-A3 | *Cdkn3* | 1.8523 | 0.0029 |
| PEC-A3 | *Cenpa* | 1.1308 | 0.0176 |
| PEC-A3 | *Cks1b* | 1.3809 | 0.0008 |
| PEC-A3 | *Dut* | 0.3837 | 0.0384 |
| PEC-A3 | *Hmga1* | 0.6648 | 0.0044 |
| PEC-A3 | *Hmgb2* | 0.9649 | 0.0034 |
| PEC-A3 | *Hmgn2* | 0.4670 | 0.0154 |
| PEC-A3 | *Hmmr* | 1.1883 | 0.0106 |
| PEC-A3 | *Hnrnpab* | 0.3321 | 0.0367 |
| PEC-A3 | *Jpt1* | 0.6692 | 0.0031 |
| PEC-A3 | *Lgals1* | 2.6014 | 0.0000 |
| PEC-A3 | *Lgals3* | 2.7511 | 0.0012 |
| PEC-A3 | *Lsm2* | 0.5770 | 0.0104 |
| PEC-A3 | *Lsm5* | 0.8138 | 0.0021 |
| PEC-A3 | *Nhp2* | 0.9058 | 0.0026 |
| PEC-A3 | *Nme1* | 0.7543 | 0.0021 |
| PEC-A3 | *Npm1* | 0.5980 | 0.0020 |
| PEC-A3 | *Pbk* | 2.4393 | 0.0027 |
| PEC-A3 | *Pcna* | 0.6467 | 0.0007 |
| PEC-A3 | *Pkm* | 0.9116 | 0.0020 |
| PEC-A3 | *Ppp1r14b* | 0.7883 | 0.0005 |
| PEC-A3 | *Prc1* | 1.8907 | 0.0051 |
| PEC-A3 | *Racgap1* | 1.2914 | 0.0383 |
| PEC-A3 | *Ran* | 0.5929 | 0.0008 |
| PEC-A3 | *Ranbp1* | 0.6094 | 0.0011 |
| PEC-A3 | *Rpl12* | 0.4311 | 0.0119 |
| PEC-A3 | *Rps2* | 0.3174 | 0.0310 |
| PEC-A3 | *S100a10* | 1.6053 | 0.0009 |
| PEC-A3 | *S100a6* | 3.1154 | 0.0010 |
| PEC-A3 | *Serpine1* | 4.1478 | 0.0000 |
| PEC-A3 | *Set* | 0.4138 | 0.0486 |
| PEC-A3 | *Sh3bgrl3* | 0.4297 | 0.0159 |
| PEC-A3 | *Smc2* | 0.4259 | 0.0245 |

| | | | |
|---|---|---|---|
| PEC-A3 | *Smc2* | 0.60 | 0.01 |
| PEC-A3 | *Smc4* | 0.80 | 0.00 |
| PEC-A3 | *Snrpd1* | 0.65 | 0.00 |
| PEC-A3 | *Spc24* | 2.27 | 0.00 |
| PEC-A3 | *Tk1* | 1.59 | 0.00 |
| PEC-A3 | *Top2a* | 2.63 | 0.00 |
| PEC-A3 | *Tuba1b* | 1.13 | 0.01 |
| PEC-A3 | *Tuba1c* | 0.75 | 0.00 |
| PEC-A3 | *Tubb4b* | 0.42 | 0.01 |
| PEC-A3 | *Tubb5* | 0.62 | 0.00 |
| PEC-A3 | *Tubb6* | 1.74 | 0.00 |
| PEC-A3 | *Tyms* | 0.64 | 0.01 |
| PEC-A3 | *Ube2c* | 2.34 | 0.00 |
| PEC-A3 | *Ube2s* | 0.61 | 0.01 |
| PEC-A3 | *Vim* | 1.67 | 0.00 |
| PEC-A3 | *Ywhah* | 0.56 | 0.00 |

| Cluster | Gene | logFC PANvsCtl | adj.P.Val PANvsCtl |
|---|---|---|---|
| PEC-A4 | *Ech1* | 0.40 | 0.04 |
| PEC-A4 | *Gstm5* | 1.41 | 0.02 |
| PEC-A4 | *Hmgcs2* | 1.49 | 0.01 |
| PEC-A4 | *Ldhb* | -0.67 | 0.04 |
| PEC-A4 | *mt-Atp6* | -1.45 | 0.02 |
| PEC-A4 | *mt-Co1* | -1.43 | 0.02 |
| PEC-A4 | *mt-Co2* | -1.31 | 0.03 |
| PEC-A4 | *mt-Co3* | -1.43 | 0.02 |
| PEC-A4 | *mt-Nd1* | -1.43 | 0.02 |
| PEC-A4 | *mt-Nd2* | -1.39 | 0.02 |
| PEC-A4 | *mt-Nd3* | -1.21 | 0.04 |
| PEC-A4 | *mt-Nd4* | -1.46 | 0.02 |
| PEC-A4 | *mt-Nd4l* | -1.49 | 0.01 |
| PEC-A4 | *mt-Nd5* | -1.44 | 0.03 |
| PEC-A4 | *Prxl2a* | 1.71 | 0.01 |
| PEC-A4 | *Slco1a6* | 2.07 | 0.05 |
| PEC-A4 | *Tmbim4* | 0.44 | 0.04 |

| Cluster | Gene | logFC PANvsCtl | adj.P.Val PANvsCtl |
|---|---|---|---|
| PEC-B | *Acta2* | 2.86 | 0.00 |
| PEC-B | *Agtr1a* | 0.92 | 0.01 |
| PEC-B | *Aldh1a2* | 1.19 | 0.01 |
| PEC-B | *Arhgdib* | 0.97 | 0.00 |
| PEC-B | *Arpc1b* | 0.58 | 0.01 |
| PEC-B | *Bgn* | 1.74 | 0.00 |
| PEC-B | *Bok* | 1.17 | 0.00 |
| PEC-B | *C1qtnf2* | 1.69 | 0.00 |
| PEC-B | *C1qtnf6* | 1.21 | 0.00 |
| PEC-B | *Cd302* | 0.88 | 0.01 |
| PEC-B | *Cdh11* | 2.02 | 0.02 |
| PEC-B | *Cebpb* | 1.04 | 0.01 |
| PEC-B | *Cfh* | 1.65 | 0.01 |
| PEC-B | *Ckb* | 0.96 | 0.00 |
| PEC-B | *Clec11a* | 2.42 | 0.00 |
| PEC-B | *Col1a1* | 3.87 | 0.00 |
| PEC-B | *Col4a1* | 1.07 | 0.00 |
| PEC-B | *Col4a2* | 1.03 | 0.00 |
| PEC-B | *Col5a1* | 1.87 | 0.00 |
| PEC-B | *Col5a2* | 1.03 | 0.00 |
| PEC-B | *Col6a2* | 2.47 | 0.03 |
| PEC-B | *Col8a1* | 3.02 | 0.00 |
| PEC-B | *Colec12* | 1.87 | 0.03 |

| | | | |
|---|---|---|---|
| PEC-A3 | *Smc4* | 0.7025 | 0.0018 |
| PEC-A3 | *Snrpd1* | 0.6074 | 0.0020 |
| PEC-A3 | *Snrpe* | 0.6350 | 0.0326 |
| PEC-A3 | *Spc24* | 1.8050 | 0.0011 |
| PEC-A3 | *Tk1* | 1.0646 | 0.0087 |
| PEC-A3 | *Top2a* | 1.9162 | 0.0056 |
| PEC-A3 | *Tuba1b* | 0.9771 | 0.0100 |
| PEC-A3 | *Tuba1c* | 0.5565 | 0.0124 |
| PEC-A3 | *Tubb4b* | 0.3289 | 0.0166 |
| PEC-A3 | *Tubb5* | 0.5391 | 0.0045 |
| PEC-A3 | *Tubb6* | 1.5078 | 0.0000 |
| PEC-A3 | *Tyms* | 0.7153 | 0.0029 |
| PEC-A3 | *Ube2c* | 1.6978 | 0.0045 |
| PEC-A3 | *Ube2s* | 0.6370 | 0.0035 |
| PEC-A3 | *Vim* | 1.7217 | 0.0014 |
| PEC-A3 | *Ywhah* | 0.7762 | 0.0002 |

| Cluster | Gene | logFC ADRvsCtl | adj.P.Val ADRvsCtl |
|---|---|---|---|
| PEC-A4 | *Asl* | 0.7529 | 0.0418 |
| PEC-A4 | *Ass1* | -1.1729 | 0.0424 |
| PEC-A4 | *Cat* | -0.7537 | 0.0309 |
| PEC-A4 | *Cd74* | 0.8873 | 0.0101 |
| PEC-A4 | *Cgref1* | 2.3039 | 0.0009 |
| PEC-A4 | *Cox7c* | 0.5696 | 0.0252 |
| PEC-A4 | *Cyp4b1* | 1.2654 | 0.0022 |
| PEC-A4 | *Dnase1* | -1.8719 | 0.0226 |
| PEC-A4 | *Ech1* | 0.7788 | 0.0006 |
| PEC-A4 | *Gcnt1* | 1.1670 | 0.0004 |
| PEC-A4 | *Gstm5* | 1.3801 | 0.0198 |
| PEC-A4 | *Hmgcs2* | 2.2428 | 0.0006 |
| PEC-A4 | *Ldhb* | -0.7109 | 0.0237 |
| PEC-A4 | *Me1* | 0.9461 | 0.0054 |
| PEC-A4 | *mt-Atp6* | -1.4064 | 0.0160 |
| PEC-A4 | *mt-Co1* | -1.4103 | 0.0121 |
| PEC-A4 | *mt-Co2* | -1.1775 | 0.0395 |
| PEC-A4 | *mt-Co3* | -1.4586 | 0.0122 |
| PEC-A4 | *mt-Nd1* | -1.3667 | 0.0199 |
| PEC-A4 | *mt-Nd2* | -1.2786 | 0.0202 |
| PEC-A4 | *mt-Nd4* | -1.3970 | 0.0141 |
| PEC-A4 | *mt-Nd4l* | -1.4590 | 0.0097 |
| PEC-A4 | *mt-Nd5* | -1.5159 | 0.0148 |
| PEC-A4 | *Prxl2a* | 1.8133 | 0.0042 |
| PEC-A4 | *Ptgds* | -1.8303 | 0.0045 |
| PEC-A4 | *Rbm47* | -1.1723 | 0.0181 |
| PEC-A4 | *Slc22a12* | 1.1365 | 0.0298 |
| PEC-A4 | *Slco1a6* | 2.8555 | 0.0059 |
| PEC-A4 | *Tmbim4* | 0.7517 | 0.0015 |

| Cluster | Gene | logFC ADRvsCtl | adj.P.Val ADRvsCtl |
|---|---|---|---|
| PEC-B | *Abca8a* | 3.5724 | 0.0141 |
| PEC-B | *Acta2* | 3.1511 | 0.0001 |
| PEC-B | *Agtr1a* | 0.8082 | 0.0069 |
| PEC-B | *Aldh1a2* | 1.6331 | 0.0011 |
| PEC-B | *Angptl4* | 3.2139 | 0.0036 |
| PEC-B | *Arhgdib* | 1.0893 | 0.0005 |
| PEC-B | *Arpc1b* | 0.7017 | 0.0010 |
| PEC-B | *B2m* | 0.6588 | 0.0037 |
| PEC-B | *Bgn* | 1.7377 | 0.0006 |
| PEC-B | *Bok* | 1.5650 | 0.0000 |
| PEC-B | *C1qtnf2* | 1.6823 | 0.0003 |
| PEC-B | *C1qtnf6* | 0.7026 | 0.0033 |
| PEC-B | *Ccl6* | 1.6228 | 0.0022 |
| PEC-B | *Cd302* | 1.2290 | 0.0008 |
| PEC-B | *Cdh11* | 2.3848 | 0.0066 |
| PEC-B | *Cebpb* | 1.5783 | 0.0004 |
| PEC-B | *Cebpd* | 1.1279 | 0.0039 |
| PEC-B | *Cfh* | 2.1798 | 0.0017 |
| PEC-B | *Ckb* | 1.2341 | 0.0001 |
| PEC-B | *Clec11a* | 1.9980 | 0.0008 |
| PEC-B | *Col15a1* | 2.6522 | 0.0281 |
| PEC-B | *Col1a1* | 4.2187 | 0.0005 |
| PEC-B | *Col3a1* | 4.8122 | 0.0179 |

| | | | |
|---|---|---|---|
| PEC-B | *Cped1* | 0.79 | 0.04 |
| PEC-B | *Cst3* | 2.77 | 0.00 |
| PEC-B | *Ctsk* | 1.60 | 0.00 |
| PEC-B | *Cxcl12* | 0.90 | 0.02 |
| PEC-B | *Cygb* | 1.75 | 0.00 |
| PEC-B | *Dapk2* | 1.11 | 0.01 |
| PEC-B | *Ddr2* | 1.84 | 0.01 |
| PEC-B | *Dkk2* | 0.97 | 0.01 |
| PEC-B | *Ecm1* | 2.21 | 0.00 |
| PEC-B | *Efemp2* | 1.43 | 0.00 |
| PEC-B | *Eln* | 1.26 | 0.01 |
| PEC-B | *Emilin1* | 1.40 | 0.00 |
| PEC-B | *Eng* | 0.68 | 0.02 |
| PEC-B | *Eva1b* | 0.73 | 0.00 |
| PEC-B | *F2r* | 1.26 | 0.00 |
| PEC-B | *Fblim1* | 2.56 | 0.01 |
| PEC-B | *Fbln5* | 1.45 | 0.00 |
| PEC-B | *Fbln7* | 1.76 | 0.02 |
| PEC-B | *Fbn1* | 1.94 | 0.00 |
| PEC-B | *Fhl2* | 2.06 | 0.00 |
| PEC-B | *Fkbp7* | 1.21 | 0.01 |
| PEC-B | *Frzb* | 1.51 | 0.01 |
| PEC-B | *Fstl1* | 1.49 | 0.00 |
| PEC-B | *Gpm6b* | 1.26 | 0.01 |
| PEC-B | *Gucy1a1* | 1.14 | 0.01 |
| PEC-B | *Gucy1b1* | 1.52 | 0.00 |
| PEC-B | *Ifitm1* | 0.68 | 0.01 |
| PEC-B | *Ifitm2* | 0.56 | 0.01 |
| PEC-B | *Ifitm3* | 0.70 | 0.03 |
| PEC-B | *Igfbp3* | 1.24 | 0.04 |
| PEC-B | *Igfbp4* | 1.05 | 0.00 |
| PEC-B | *Il6st* | 0.79 | 0.02 |
| PEC-B | *Lbh* | 0.62 | 0.02 |
| PEC-B | *Lgals1* | 2.07 | 0.00 |
| PEC-B | *Lgals3bp* | 1.09 | 0.01 |
| PEC-B | *Lgals9* | 0.66 | 0.05 |
| PEC-B | *Loxl1* | 1.79 | 0.00 |
| PEC-B | *Lpl* | 0.74 | 0.01 |
| PEC-B | *Lrp1* | 1.29 | 0.00 |
| PEC-B | *Marcks* | 0.70 | 0.01 |
| PEC-B | *Marcksl1* | 0.70 | 0.02 |
| PEC-B | *Mfge8* | 1.45 | 0.00 |
| PEC-B | *Mgp* | 1.79 | 0.00 |
| PEC-B | *Mmd* | 0.70 | 0.02 |
| PEC-B | *Mmp14* | 1.48 | 0.00 |
| PEC-B | *Mrc2* | 1.86 | 0.01 |
| PEC-B | *Mxra8* | 1.10 | 0.00 |
| PEC-B | *Myl9* | 1.98 | 0.00 |
| PEC-B | *Myo1b* | 0.70 | 0.02 |
| PEC-B | *Ndufa4l2* | 1.20 | 0.01 |
| PEC-B | *Nid1* | 1.68 | 0.00 |
| PEC-B | *Npc2* | 0.81 | 0.01 |
| PEC-B | *Nt5dc2* | 1.17 | 0.00 |
| PEC-B | *Oaf* | 0.67 | 0.02 |
| PEC-B | *Olfml3* | 0.94 | 0.00 |
| PEC-B | *P3h3* | 1.44 | 0.00 |
| PEC-B | *Pcolce* | 2.52 | 0.00 |
| PEC-B | *Pdgfrb* | 1.61 | 0.00 |
| PEC-B | *Pdlim1* | 1.55 | 0.00 |
| PEC-B | *Plvap* | 1.40 | 0.00 |
| PEC-B | *Pmepa1* | 1.01 | 0.01 |
| PEC-B | *Ppib* | 0.60 | 0.01 |
| PEC-B | *Ppic* | 2.07 | 0.00 |
| PEC-B | *Ppp1r14a* | 1.97 | 0.00 |
| PEC-B | *Pros1* | 0.72 | 0.01 |
| PEC-B | *Rarres2* | 1.22 | 0.02 |
| PEC-B | *Rasgrp2* | 1.14 | 0.00 |
| PEC-B | *Rcn3* | 1.95 | 0.00 |
| PEC-B | *Rem1* | 1.16 | 0.00 |
| PEC-B | *Rerg* | 1.63 | 0.01 |
| PEC-B | *Scarf2* | 1.86 | 0.00 |
| PEC-B | *Scn1b* | 0.88 | 0.01 |

| | | | |
|---|---|---|---|
| PEC-B | *Col4a1* | 0.8217 | 0.0027 |
| PEC-B | *Col4a2* | 0.7796 | 0.0037 |
| PEC-B | *Col5a1* | 1.6387 | 0.0010 |
| PEC-B | *Col5a2* | 0.7771 | 0.0055 |
| PEC-B | *Col6a1* | 3.5742 | 0.0176 |
| PEC-B | *Col6a2* | 2.7842 | 0.0123 |
| PEC-B | *Col8a1* | 3.4395 | 0.0001 |
| PEC-B | *Colec12* | 2.5569 | 0.0042 |
| PEC-B | *Cped1* | 1.2863 | 0.0021 |
| PEC-B | *Cst3* | 2.7847 | 0.0014 |
| PEC-B | *Ctsk* | 1.6876 | 0.0005 |
| PEC-B | *Cxcl12* | 0.7826 | 0.0306 |
| PEC-B | *Cxcl16* | 0.4352 | 0.0427 |
| PEC-B | *Cygb* | 1.5933 | 0.0021 |
| PEC-B | *Dapk2* | 1.4962 | 0.0016 |
| PEC-B | *Dkk2* | 1.1766 | 0.0018 |
| PEC-B | *Ecm1* | 2.4959 | 0.0003 |
| PEC-B | *Efemp2* | 1.5208 | 0.0004 |
| PEC-B | *Eln* | 1.3646 | 0.0036 |
| PEC-B | *Emilin1* | 1.2100 | 0.0025 |
| PEC-B | *Emp1* | 0.5127 | 0.0119 |
| PEC-B | *Eng* | 0.6257 | 0.0164 |
| PEC-B | *Eva1b* | 0.6302 | 0.0058 |
| PEC-B | *F2r* | 1.1871 | 0.0010 |
| PEC-B | *Fblim1* | 3.0116 | 0.0028 |
| PEC-B | *Fbln5* | 1.4459 | 0.0006 |
| PEC-B | *Fbln7* | 1.6102 | 0.0232 |
| PEC-B | *Fbn1* | 1.6667 | 0.0004 |
| PEC-B | *Fhl2* | 2.0203 | 0.0004 |
| PEC-B | *Fkbp7* | 1.2120 | 0.0036 |
| PEC-B | *Frzb* | 2.1732 | 0.0004 |
| PEC-B | *Fstl1* | 1.3632 | 0.0008 |
| PEC-B | *Fxyd5* | 1.0959 | 0.0047 |
| PEC-B | *Gng11* | 0.6284 | 0.0100 |
| PEC-B | *Gpm6b* | 1.6560 | 0.0005 |
| PEC-B | *Gucy1a1* | 1.1074 | 0.0042 |
| PEC-B | *Gucy1b1* | 1.5003 | 0.0021 |
| PEC-B | *Ifitm1* | 0.7539 | 0.0054 |
| PEC-B | *Ifitm2* | 0.6545 | 0.0033 |
| PEC-B | *Ifitm3* | 0.8964 | 0.0056 |
| PEC-B | *Igfbp3* | 1.7364 | 0.0048 |
| PEC-B | *Igfbp4* | 0.8490 | 0.0065 |
| PEC-B | *Igfbp5* | -0.7687 | 0.0085 |
| PEC-B | *Il11ra1* | 0.5241 | 0.0235 |
| PEC-B | *Il6st* | 0.8424 | 0.0078 |
| PEC-B | *Lbh* | 0.5569 | 0.0255 |
| PEC-B | *Lgals1* | 2.6014 | 0.0000 |
| PEC-B | *Lgals3bp* | 1.2423 | 0.0016 |
| PEC-B | *Loxl1* | 1.7208 | 0.0005 |
| PEC-B | *Lpar1* | 1.6050 | 0.0324 |
| PEC-B | *Lpl* | 1.0535 | 0.0004 |
| PEC-B | *Lrp1* | 1.1495 | 0.0020 |
| PEC-B | *Marcks* | 0.6001 | 0.0134 |
| PEC-B | *Mfge8* | 1.5614 | 0.0001 |
| PEC-B | *Mgp* | 1.8149 | 0.0009 |
| PEC-B | *Mmd* | 0.8347 | 0.0055 |
| PEC-B | *Mmp14* | 1.5311 | 0.0006 |
| PEC-B | *Mrc2* | 1.7158 | 0.0082 |
| PEC-B | *Mt1* | 1.6662 | 0.0345 |
| PEC-B | *Mxra8* | 0.8466 | 0.0049 |
| PEC-B | *Myl9* | 2.2943 | 0.0000 |
| PEC-B | *Myo1b* | 0.5948 | 0.0278 |
| PEC-B | *Ndufa4l2* | 1.1554 | 0.0042 |
| PEC-B | *Nid1* | 1.3727 | 0.0012 |
| PEC-B | *Npc2* | 1.0981 | 0.0004 |
| PEC-B | *Nt5dc2* | 1.0643 | 0.0046 |
| PEC-B | *Oaf* | 0.6177 | 0.0242 |
| PEC-B | *Olfml3* | 0.9492 | 0.0028 |
| PEC-B | *Osmr* | 0.7432 | 0.0462 |
| PEC-B | *P3h3* | 1.4304 | 0.0007 |
| PEC-B | *Pcolce* | 2.4631 | 0.0014 |
| PEC-B | *Pdgfrb* | 1.3334 | 0.0021 |

| | | | |
|---|---|---|---|
| PEC-B | *Selenom* | 0.81 | 0.00 |
| PEC-B | *Serpine2* | 1.73 | 0.00 |
| PEC-B | *Serping1* | 1.65 | 0.01 |
| PEC-B | *Serpinh1* | 0.81 | 0.00 |
| PEC-B | *Slco2b1* | 2.52 | 0.00 |
| PEC-B | *Smoc2* | 2.99 | 0.00 |
| PEC-B | *Sparc* | 1.24 | 0.01 |
| PEC-B | *Srpx* | 1.86 | 0.00 |
| PEC-B | *Stbd1* | 0.57 | 0.05 |
| PEC-B | *Tcf21* | 2.27 | 0.01 |
| PEC-B | *Tgfb1* | 0.81 | 0.00 |
| PEC-B | *Tgfbi* | 1.16 | 0.01 |
| PEC-B | *Timp1* | 1.26 | 0.01 |
| PEC-B | *Tmem176a* | 1.04 | 0.01 |
| PEC-B | *Tpm2* | 2.33 | 0.00 |
| PEC-B | *Tril* | 0.94 | 0.04 |
| PEC-B | *Vegfd* | 2.48 | 0.00 |
| PEC-B | *Vstm4* | 1.90 | 0.00 |
| PEC-B | *Zeb2* | 0.57 | 0.04 |

| | | | |
|---|---|---|---|
| PEC-B | *Pdlim1* | 2.0216 | 0.0002 |
| PEC-B | *Pik3r1* | 0.4576 | 0.0370 |
| PEC-B | *Plvap* | 1.3185 | 0.0002 |
| PEC-B | *Pmepa1* | 0.8158 | 0.0125 |
| PEC-B | *Ppib* | 0.7804 | 0.0011 |
| PEC-B | *Ppic* | 2.4082 | 0.0001 |
| PEC-B | *Ppp1r14a* | 2.5208 | 0.0000 |
| PEC-B | *Prelp* | 1.3913 | 0.0426 |
| PEC-B | *Pros1* | 1.1396 | 0.0004 |
| PEC-B | *Rarres2* | 1.4341 | 0.0054 |
| PEC-B | *Rasgrp2* | 0.9280 | 0.0014 |
| PEC-B | *Rcn3* | 2.0752 | 0.0007 |
| PEC-B | *Rem1* | 0.8750 | 0.0130 |
| PEC-B | *Rerg* | 1.8806 | 0.0020 |
| PEC-B | *Scarf2* | 1.6085 | 0.0003 |
| PEC-B | *Scn1b* | 1.2476 | 0.0003 |
| PEC-B | *Selenom* | 0.9560 | 0.0005 |
| PEC-B | *Serpine2* | 2.1849 | 0.0000 |
| PEC-B | *Serping1* | 1.9386 | 0.0017 |
| PEC-B | *Serpinh1* | 0.8279 | 0.0010 |
| PEC-B | *Slco2b1* | 2.7406 | 0.0014 |
| PEC-B | *Smoc2* | 3.2837 | 0.0007 |
| PEC-B | *Sparc* | 1.2253 | 0.0048 |
| PEC-B | *Srpx* | 1.7620 | 0.0014 |
| PEC-B | *Stbd1* | 0.9533 | 0.0024 |
| PEC-B | *Tcf21* | 2.0110 | 0.0152 |
| PEC-B | *Tgfb1* | 0.7104 | 0.0030 |
| PEC-B | *Tgfbi* | 1.2180 | 0.0045 |
| PEC-B | *Timp1* | 1.7362 | 0.0009 |
| PEC-B | *Tmem176a* | 1.3427 | 0.0020 |
| PEC-B | *Tnfrsf1a* | 0.6422 | 0.0234 |
| PEC-B | *Tpm2* | 2.8958 | 0.0000 |
| PEC-B | *Vegfd* | 2.3416 | 0.0000 |
| PEC-B | *Vstm4* | 2.1977 | 0.0000 |
| PEC-B | *Zeb2* | 0.5621 | 0.0292 |

Supplementary Table 2: Deconvolution of AS genes in PAN and ADR models for PEC subtypes

| | **PAN** | |
|---|---|---|
| **Cluster** | **Gene** | **most Sig *P* -adj.** |
| PEC-A1 | *Nphs2* | 0.0000 |
| PEC-A1 | *Itgb1* | 0.0001 |
| PEC-A1 | *Nphs1* | 0.0001 |
| PEC-A1 | *Podxl* | 0.0002 |
| PEC-A1 | *Aif1l* | 0.0002 |
| PEC-A1 | *Enpep* | 0.0004 |
| PEC-A1 | *Lrrfip1* | 0.0014 |
| PEC-A1 | *Schip1* | 0.0014 |
| PEC-A1 | *Efnb1* | 0.0035 |
| PEC-A1 | *Actb* | 0.0037 |
| PEC-A1 | *Dpp4* | 0.0038 |
| PEC-A1 | *Clic5* | 0.0046 |
| PEC-A1 | *Col4a4* | 0.0068 |
| PEC-A1 | *Rgs3* | 0.0096 |

| | **ADR** | |
|---|---|---|
| **Cluster** | **Gene** | **most Sig *P* -adj.** |
| PEC-A1 | *Itgb1* | 0.0000 |
| PEC-A1 | *Nphs1* | 0.0000 |
| PEC-A1 | *Podxl* | 0.0000 |
| PEC-A1 | *Clic5* | 0.0000 |
| PEC-A1 | *Golim4* | 0.0000 |
| PEC-A1 | *Slc9a3r2* | 0.0000 |
| PEC-A1 | *Tspan13* | 0.0001 |
| PEC-A1 | *Sema3g* | 0.0001 |
| PEC-A1 | *Dpysl2* | 0.0002 |
| PEC-A1 | *Sdc2* | 0.0002 |
| PEC-A1 | *Crim1* | 0.0003 |
| PEC-A1 | *Klf2* | 0.0003 |
| PEC-A1 | *Cd82* | 0.0005 |
| PEC-A1 | *Dpp4* | 0.0007 |
| PEC-A1 | *Col4a3* | 0.0008 |
| PEC-A1 | *Col4a4* | 0.0010 |
| PEC-A1 | *Lrrfip1* | 0.0012 |
| PEC-A1 | *Rhpn1* | 0.0016 |
| PEC-A1 | *P3h2* | 0.0018 |
| PEC-A1 | *Pth1r* | 0.0020 |
| PEC-A1 | *Tmsb4x* | 0.0023 |
| PEC-A1 | *Myl12a* | 0.0023 |
| PEC-A1 | *Cpq* | 0.0023 |
| PEC-A1 | *Rgs3* | 0.0024 |
| PEC-A1 | *Anxa3* | 0.0029 |
| PEC-A1 | *Nsf* | 0.0036 |
| PEC-A1 | *Schip1* | 0.0036 |
| PEC-A1 | *Cavin1* | 0.0037 |
| PEC-A1 | *Plxdc2* | 0.0039 |
| PEC-A1 | *Eif3m* | 0.0057 |
| PEC-A1 | *Ccn2* | 0.0074 |
| PEC-A1 | *Lsp1* | 0.0093 |

| Cluster | Gene | most Sig *P* -adj. |
|---|---|---|
| PEC-A2 | - | |

| Cluster | Gene | most Sig *P* -adj. |
|---|---|---|
| PEC-A2 | *Cpe* | 0.0036 |
| PEC-A2 | *Clu* | 0.0000 |
| PEC-A2 | *Tsc22d1* | 0.0000 |
| PEC-A2 | *Cdh16* | 0.0002 |
| PEC-A2 | *Akap12* | 0.0002 |
| PEC-A2 | *Pax8* | 0.0035 |
| PEC-A2 | *Adgrg1* | 0.0043 |
| PEC-A2 | *Wsb1* | 0.0049 |
| PEC-A2 | *Tinagl1* | 0.0053 |
| PEC-A2 | *Lsp1* | 0.0073 |
| PEC-A2 | *Adamts1* | 0.0080 |

| Cluster | Gene | most Sig *P* -adj. |
|---|---|---|
| PEC-A3 | - | |

| Cluster | Gene | most Sig *P* -adj. |
|---|---|---|
| PEC-A3 | *Smc4* | 0.0007 |
| PEC-A3 | *Ran* | 0.0009 |
| PEC-A3 | *Tmpo* | 0.0031 |
| PEC-A3 | *Tubb5* | 0.0031 |
| PEC-A3 | *Npm1* | 0.0034 |
| PEC-A3 | *Hspd1* | 0.0036 |
| PEC-A3 | *Rpsa* | 0.0037 |
| PEC-A3 | *Gapdh* | 0.0039 |
| PEC-A3 | *Cd9* | 0.0053 |
| PEC-A3 | *Txn1* | 0.0066 |
| PEC-A3 | *Tubb4b* | 0.0067 |
| PEC-A3 | *Hmgn2* | 0.0078 |

| | | |
|---|---|---|
| PEC-A3 | *Rpl12* | 0.0090 |
| PEC-A3 | *Anxa2* | 0.0091 |
| PEC-A3 | *Krt8* | 0.0093 |

| Cluster | Gene | most Sig *P* -adj. |
|---|---|---|
| PEC-A4 | *Aldob* | 0.0045 |
| PEC-A4 | *Atp1a1* | 0.0072 |
| PEC-A4 | *Cycs* | 0.0085 |
| PEC-A4 | *Ddt* | 0.0008 |

| Cluster | Gene | most Sig *P* -adj. |
|---|---|---|
| PEC-A4 | *Aadat* | 0.0027 |
| PEC-A4 | *Abhd14b* | 0.0040 |
| PEC-A4 | *Acat1* | 0.0088 |
| PEC-A4 | *Acox1* | 0.0007 |
| PEC-A4 | *Acsm2* | 0.0001 |
| PEC-A4 | *Akr1a1* | 0.0002 |
| PEC-A4 | *Aldob* | 0.0025 |
| PEC-A4 | *Amacr* | 0.0095 |
| PEC-A4 | *Aqp1* | 0.0007 |
| PEC-A4 | *Ass1* | 0.0001 |
| PEC-A4 | *Atp1a1* | 0.0000 |
| PEC-A4 | *Bdh2* | 0.0023 |
| PEC-A4 | *Bnip3* | 0.0058 |
| PEC-A4 | *Bphl* | 0.0007 |
| PEC-A4 | *Calml4* | 0.0058 |
| PEC-A4 | *Car2* | 0.0006 |
| PEC-A4 | *Cd74* | 0.0000 |
| PEC-A4 | *Cisd1* | 0.0042 |
| PEC-A4 | *Cldn10* | 0.0001 |
| PEC-A4 | *Cltrn* | 0.0020 |
| PEC-A4 | *Cmbl* | 0.0035 |
| PEC-A4 | *Cndp2* | 0.0000 |
| PEC-A4 | *Cox5a* | 0.0084 |
| PEC-A4 | *Cyc1* | 0.0015 |
| PEC-A4 | *Cycs* | 0.0012 |
| PEC-A4 | *Dab2* | 0.0004 |
| PEC-A4 | *Dbi* | 0.0000 |
| PEC-A4 | *Dcxr* | 0.0029 |
| PEC-A4 | *Ddah1* | 0.0007 |
| PEC-A4 | *Ddt* | 0.0030 |
| PEC-A4 | *Ech1* | 0.0021 |
| PEC-A4 | *Ephx2* | 0.0051 |
| PEC-A4 | *Etfa* | 0.0021 |
| PEC-A4 | *Ethe1* | 0.0017 |
| PEC-A4 | *Fbp1* | 0.0035 |
| PEC-A4 | *Fmo1* | 0.0017 |
| PEC-A4 | *Folr1* | 0.0049 |
| PEC-A4 | *Gapdh* | 0.0031 |
| PEC-A4 | *Gatm* | 0.0018 |
| PEC-A4 | *Gclm* | 0.0085 |
| PEC-A4 | *Gpd1* | 0.0030 |
| PEC-A4 | *Grhpr* | 0.0001 |
| PEC-A4 | *Gsr* | 0.0007 |
| PEC-A4 | *Gstt2* | 0.0046 |
| PEC-A4 | *Hagh* | 0.0095 |
| PEC-A4 | *Hao2* | 0.0029 |
| PEC-A4 | *Hgd* | 0.0001 |
| PEC-A4 | *Hibadh* | 0.0031 |
| PEC-A4 | *Hnf4a* | 0.0012 |
| PEC-A4 | *Hpn* | 0.0054 |
| PEC-A4 | *Idh1* | 0.0000 |
| PEC-A4 | *Ivns1abp* | 0.0000 |
| PEC-A4 | *Khk* | 0.0098 |
| PEC-A4 | *Krt8* | 0.0034 |
| PEC-A4 | *Ldhb* | 0.0000 |
| PEC-A4 | *Ldhd* | 0.0000 |
| PEC-A4 | *Lgmn* | 0.0050 |
| PEC-A4 | *Lrp2* | 0.0019 |
| PEC-A4 | *Mcrip2* | 0.0049 |

| | | |
|---|---|---|
| PEC-A4 | *Mep1a* | 0.0007 |
| PEC-A4 | *Mif* | 0.0030 |
| PEC-A4 | *Mpc2* | 0.0020 |
| PEC-A4 | *Mrpl12* | 0.0061 |
| PEC-A4 | *Napsa* | 0.0008 |
| PEC-A4 | *Ndrg1* | 0.0000 |
| PEC-A4 | *Ndufa6* | 0.0001 |
| PEC-A4 | *Ndufv3* | 0.0065 |
| PEC-A4 | *Nipsnap1* | 0.0001 |
| PEC-A4 | *Pah* | 0.0038 |
| PEC-A4 | *Park7* | 0.0003 |
| PEC-A4 | *Pck1* | 0.0000 |
| PEC-A4 | *Pdzk1ip1* | 0.0072 |
| PEC-A4 | *Phyh* | 0.0006 |
| PEC-A4 | *Plscr2* | 0.0000 |
| PEC-A4 | *Qdpr* | 0.0085 |
| PEC-A4 | *Rab3ip* | 0.0100 |
| PEC-A4 | *Slc13a1* | 0.0001 |
| PEC-A4 | *Slc13a3* | 0.0001 |
| PEC-A4 | *Slc17a3* | 0.0009 |
| PEC-A4 | *Slc22a18* | 0.0002 |
| PEC-A4 | *Slc22a6* | 0.0047 |
| PEC-A4 | *Slc25a10* | 0.0011 |
| PEC-A4 | *Slc25a39* | 0.0001 |
| PEC-A4 | *Slc27a2* | 0.0005 |
| PEC-A4 | *Slc34a1* | 0.0053 |
| PEC-A4 | *Slc47a1* | 0.0031 |
| PEC-A4 | *Slc7a7* | 0.0088 |
| PEC-A4 | *Slc7a9* | 0.0033 |
| PEC-A4 | *Slc9a3r1* | 0.0077 |
| PEC-A4 | *Sod2* | 0.0043 |
| PEC-A4 | *Spink1* | 0.0062 |
| PEC-A4 | *Suclg1* | 0.0006 |
| PEC-A4 | *Tcn2* | 0.0012 |
| PEC-A4 | *Thnsl2* | 0.0054 |
| PEC-A4 | *Tkfc* | 0.0022 |
| PEC-A4 | *Tmbim4* | 0.0058 |
| PEC-A4 | *Ttc36* | 0.0030 |
| PEC-A4 | *Uqcr11* | 0.0008 |
| PEC-A4 | *Uqcrfs1* | 0.0000 |
| PEC-A4 | *Uqcrq* | 0.0003 |

| Cluster | Gene | most Sig *P* -adj. |
|---|---|---|
| PEC-B | *Col4a1* | 0.0002 |
| PEC-B | *Eng* | 0.0014 |
| PEC-B | *Epas1* | 0.0000 |
| PEC-B | *Mfge8* | 0.0057 |
| PEC-B | *Plpp1* | 0.0006 |

| Cluster | Gene | most Sig *P* -adj. |
|---|---|---|
| PEC-B | *Adamtsl2* | 0.0027 |
| PEC-B | *Agtr1a* | 0.0001 |
| PEC-B | *Arhgdib* | 0.0073 |
| PEC-B | *B2m* | 0.0084 |
| PEC-B | *Bst2* | 0.0024 |
| PEC-B | *Camk2n1* | 0.0026 |
| PEC-B | *Ckb* | 0.0031 |
| PEC-B | *Col4a1* | 0.0000 |
| PEC-B | *Col4a2* | 0.0000 |
| PEC-B | *Col5a1* | 0.0093 |
| PEC-B | *Col5a2* | 0.0009 |
| PEC-B | *Cped1* | 0.0091 |
| PEC-B | *Dkk2* | 0.0030 |
| PEC-B | *Emilin1* | 0.0090 |
| PEC-B | *Emp1* | 0.0001 |
| PEC-B | *Epas1* | 0.0000 |
| PEC-B | *Fxyd5* | 0.0018 |
| PEC-B | *G0s2* | 0.0002 |
| PEC-B | *Hexa* | 0.0057 |
| PEC-B | *Ifitm1* | 0.0013 |
| PEC-B | *Igfbp5* | 0.0003 |

| | | |
|---|---|---|
| PEC-B | *Il6st* | 0.0071 |
| PEC-B | *Lgals3bp* | 0.0040 |
| PEC-B | *Lgals9* | 0.0046 |
| PEC-B | *Lpl* | 0.0025 |
| PEC-B | *Lrp1* | 0.0028 |
| PEC-B | *Man2a1* | 0.0011 |
| PEC-B | *Mfge8* | 0.0000 |
| PEC-B | *Nid1* | 0.0000 |
| PEC-B | *Npc2* | 0.0009 |
| PEC-B | *Pi16* | 0.0051 |
| PEC-B | *Plpp1* | 0.0009 |
| PEC-B | *Ppib* | 0.0032 |
| PEC-B | *Pros1* | 0.0032 |
| PEC-B | *Rbp1* | 0.0001 |
| PEC-B | *Scarb2* | 0.0096 |
| PEC-B | *Sfrp1* | 0.0039 |
| PEC-B | *Slc29a1* | 0.0000 |
| PEC-B | *Slc43a3* | 0.0041 |
| PEC-B | *Slfn5* | 0.0055 |
| PEC-B | *Tmem176a* | 0.0055 |

**Supplementary Table 3:** AS genes in PAN and ADR models represented in monogenic genes with established roles in glomerular disease

| Gene | Protein | Class | *P* adj. PAN | *P* adj. ADR | Function | Podocyte/Kidney Role |
|---|---|---|---|---|---|---|
| *Coq2* | CoenzymeQ2 / Polypolyprenyl transferase | Podocyte Mitochondria | | 3.77e-4 | It catalyzes a key step in the synthesis of CoQ (Ubiquinone), which plays a crucial role in the electron transport chain within mitochondria [1]. | Pathogenic COQ2 variants have been shown to be a primary cause of CoQ10 deficiency, which is associated with nephrotic syndrome [2]. |
| *Pdss2* | Decaprenyl Diphosphate Synthase Subunit 2 | Podocyte Mitochondria | | 4.76e-3 | A podocyte specific gene that encodes an enzyme responsible for synthesizing CoQ (Ubiquinone). CoQ is involved in the electron transport chain of mitochondria [3]. | Mutations in this gene lead to CoQ biosynthesis dysfunction, which can lead to kidney failure due to the loss of podocytes. Knockout of this gene in podocytes has been shown to cause nephrotic syndrome [4]. |
| *Ankfy1* | Ankyrin repeat and FYVE domain-containing protein 1 | Podocyte Metabolic | | 1.99e-4 | ANKFY1 acts as an effector for RAB5 (Ras-related protein in brain 5), a GTPase. In its active form, RAB5 recruits effectors like ANKFY1 to endosomes, regulating the sorting of internalized materials during endocytosis. [5]. | Mutations in ANKFY1 that enhance its binding to RAB5, leading to overactivity, have been linked to nephrotic syndrome [6]. |
| *Dlc1* | Rho GTPase-activating protein 7 | Podocyte Cytoskeletal Scaffold | | 1.09e-4 | A tumor suppressor, involved in cell polarity, proliferation, migration, and survival, and found to be downregulated in different cancer types [7]. | Mutations have been identified in patients with nephrotic syndrome. Knockdown of this gene in podocytes leads to reduced cell migration and can impact the filtration barrier [8]. |
| *Myh9* | Myosin-9 | Podocyte Cytoskeletal Scaffold | | 5.05e-3 | Subunit for the larger myosin heavy chain IIA protein, responsible for various processes such as cell movement, maintenance of cell shape and cytokinesis [9]. | Mutations have been implicated with nephropathy and proteinuria. SNPs in the *MYH9* gene have been associated with CKD, specifically glomerulopathies, among African Americans [10]. |
| *Tns2* | Tensin 2 | Podocyte Cytoskeletal Scaffold | 2.73e-3 | 2.26e-4 | A tyrosine-protein phosphatase that helps to regulate cell motility, proliferation, and links integrins to the actin cytoskeleton [11] | Predominantly expressed in podocytes in the kidneys. *Tns2*-deficient mice develop nephrotic syndrome and proteinuria [11]. |
| *Inf2* | Inverted Formin 2 | Podocyte Cytoskeletal Scaffold | 3.59e-3 | 3.45e-4 | Actin-regulating protein that influences a variety of cellular functions such as cell migration and vesicular transport [12]. | Mutations have been shown to play a role in the development of thin glomerular basement membrane and FSGS [13]. |
| *Itsn2* | Intersectin 2 | Podocyte Cytoskeletal Scaffold | 5.68e-3 | | Regulates the formation of clathrin-coated vesicles [14]. | Mutations can cause nephrotic syndrome and has been implicated in hereditary renal failure [14]. It functions as podocytic guanine nucleotide exchange factor for Cdc42 and *Itsn2-L* knockout mice recapitulate the mild nephrotic syndrome phenotype [8]. |

| | | | | | | |
|---|---|---|---|---|---|---|
| *Podxyl* | Podocalyxin | Podocyte Membrane | 6.76e-3 | 3.24e-4 | Highly expressed on the surface of podocytes and plays an essential role in the formation and maintenance of podocyte foot processes. It also plays a role in tissue remodeling and development [15]. | Mutations in this gene are linked to proteinuria-related kidney disease, with some causing anuria (failure to produce urine). Podocalyxin shedding in urine is a common biomarker of nephrotic syndrome. [16]. |
| *Scarb2* | Scavenger receptor class B member 2 or lysosomal integral membrane protein-2 (LIMP-2). | Podocyte Lysosomal | | 9.63e-3 | Found on lysosomal membrane and is required for the biogenesis and maintenance of lysosomes and endosomes [17]. | Mutations associated with action myoclonus-renal failure syndrome (SCARB2-AMRF). It is a disorder that results in progressive neurological disease and steroid-resistant nephrotic syndrome. Mice deficient in this protein show kidneys with glomerular lesions and effacement of the foot processes [18]. |
| *Lmna* | Lamin A and C | Podocyte Nuclear Proteins | 5.59e-3 | 2.35e-3 | Forms part of the nuclear lamina, a two-dimensional matrix of proteins located next to the inner nuclear membrane [19] . | Specific variations of this gene have been linked to atypical myopathy and proteinuric nephropathy [20]. |
| *Fat1* | FAT1 cadherin protein | Podocyte Slit Diaphragm | | 2.16e-4 | Activates several signaling pathways through protein-protein interactions- particularly pathways that relate to cell migration, adhesion, proliferation, and invasion [21]. | Mutations have been shown to cause glomerulotubular nephropathy. Podocyte-specific deletion of *Fat1* in mice causes abnormal glomerular filtration barrier development, leading to podocyte foot process effacement [22]. |
| *Nphs1* | Nephrin | Podocyte Slit Diaphragm | 9.62e-3 | 1.99e-4 | Podocyte cell surface protein, essential for the formation and maintenance of the slit diaphragm [23]. | Mutations in this gene are linked to nephrotic syndrome, proteinuria, glomerular hypertrophy, FSGS and eventually end-stage renal failure. *Nphs1* knockout mice don't recover from foot process effacement [24]. |
| *Nphs2* | Podocin | Podocyte Slit Diaphragm | 3.48e-3 | | Podocin is almost exclusively expressed as a membrane protein in podocytes that plays a key role in the structural integrity of the slit diaphragm [25]. | Mutations cause proteinuria and congenital nephrotic syndrome. Podocin interacts with nephrin and other proteins in the slit diaphragm. Podocin knockout mice show proteinuria at birth [26]. |
| *Myo1e* | Myosin 1e | Podocyte Actin Regulation | | 2.92e-4 | Involved in several different cellular processes, such as remodeling of cellular membrane during endocytosis and exocytosis, maintaining the glomerular filtration barrier, and controlling the movement of cytoplasmic vesicles [27]. | Regulates endocytosis in podocytes and F-actin cytoskeleton and plays an important role in renal filtration. Mutations in this gene are associated with FSGS and nephrotic syndrome [28]. |
| *Rac1* | Rac Family Small GTPase 1 | Podocyte Actin Regulation | | 6.50e-03 | A member of the Rho family of GTPases, it is located at the leading edge of migrating cells. It regulates various signaling pathways, including those involved in cell proliferation, transcription, and cytoskeleton organization [29]. | Hyperactivation of Rac1 in glomerular podocytes has been linked to proteinuric kidney diseases, while a deficiency in Rac1 activity can impair post-natal development of the renal medulla [30]. |

| | | | | | | |
|---|---|---|---|---|---|---|
| *Cdc42* | Cell division control protein42 | Podocyte Actin Regulation | | 1.22e-03 | A small GTPase that regulates various signaling pathways involved in cell morphology, cell migration, filopodia formation, endocytosis, adhesion and cytoskeletal formation [31]. | Deficiency in infants has been linked with proteinuria, and kidney failure. It is essential for kidney development and proper function, and its deletion results in cystic kidney disease. Deletion in nephron progenitors in mice formed hypoplastic kidneys with a reduced nephrogenic zone [32]. |
| *Cubn* | Cubilin | Podocyte tRNA modification | | 5.21e-03 | An extracellular protein that is expressed in both renal proximal tubular cells and podocytes. Serves many functions such as; intestinal absorption of B12, reabsorption of albumin, and transferrin, binding and uptake of several nephrotoxic proteins including light chains, myoglobin and hemoglobin [33]. | Mutations can cause structural abnormalities in the cubilin protein in the kidneys, which can lead to proteinuria [34]. |
| *Lage3* | L antigen family member 3 | Podocyte tRNA modification | | 4.63e-04 | Regulates occurrence and invasion of tumors and has been shown to promote cell proliferation, migration, and invasion [35]. | LAGE3 gene is associated with clear cell renal cell carcinoma, which is a type of kidney tumor [36]. Mutations in this gene can cause Galloway-Mowat Syndrome, which involves steroid-resistant nephrotic syndrome and microcephaly with brain anomalies [37]. |
| *Col4a3* | Collagen IV alpha 3 | Podocyte Matrix GBM | | 8.43e-05 | The α3 chain of collagen IV combines with the α4 and α5 chains to make α345(IV) collagen molecules, which attach together in complex protein networks needed to form basement membranes of several organs, including the kidneys [38]. | Variants in this gene have been associated with several glomerular basement membrane nephropathies; such as Alport Syndrome due to impairments in the formation of the GBM [39]. |
| *Col4a4* | Collagen IV alpha 4 | Podocyte Matrix GBM | 1.71e-04 | 2.09e-04 | The α4 chain of collagen IV combines with the α3 and α5 chains to make α345(IV) collagen molecules which attach together in complex protein networks needed to form basement membranes of several organs, including the kidneys [40]. | Variants in this gene have been associated with several glomerular basement membrane nephropathies; such as Alport Syndrome due to impairments in the formation of the GBM [39]. |
| *Col4a5* | Collagen IV alpha 5 | Podocyte Matrix GBM | | 9.37e-03 | The α5 chain of collagen IV combines with the α3 and α4 chains to make α345(IV) collagen molecules which attach together in complex protein networks needed to form basement membranes of several organs, including the kidneys [41]. | Variants in this gene have been associated with several glomerular basement membrane nephropathies; such as Alport Syndrome due to impairments in the formation of the GBM [41]. |
| *Lama5* | Laminin-a5 | Podocyte Matrix GBM | | 5.73e-06 | Promotes cell adhesion and migration, and maintenance and function of the extracellular matrix in several tissues, including the GBM [42]. | Mutations have been linked thinning of the GBM, proteinuria and FSGS, and development of kidney cysts and kidney dysfunction [43]. |

| | | | | | | |
|---|---|---|---|---|---|---|
| *Lamb2* | laminin-b2 | Podocyte Matrix GBM | 2.88e-03 | 2.17e-03 | Maintenance and function of the extracellular matrix in several tissues, including the ocular structures and the GBM [44]. | Mutations have been linked to proteinuria, podocyte foot effacement, Pierson syndrome, nephrotic syndrome and renal failure [45]. |

**Supplementary Table 4:** AS genes in PAN and ADR models representing novel genes in glomerular disease

| Gene | Protein | *P* adj. PAN | *P* adj. ADR | Function | Podocyte/Kidney Role | Known Splice Variants |
|---|---|---|---|---|---|---|
| *Itm2b* | Integral Transmembrane Protein 2B | 1.05e-4 | 6.2e-26 | ITM2B is mainly expressed in the brain and retina. In the brain, it inhibits amyloid β oligomerization by interacting with amyloid precursor protein. Mutation (c.782A>C, p.Glu261Ala) is linked to retinal dystrophy [1]. | It has been recently identified in the interactome of the slit diaphragm complex [2]. | *ITM2B* variant Long: Canonical variant containing 6 exons, encoding 266 aa, mostly expressed in the brain and retina.<br>*ITM2B* variant Short: Reported to be expressed as a 210 aa protein in the human retina [1]. |
| *Adgrg1* | Adhesion G Protein-Coupled Receptor G1 | --- | 5.01e-19 | GPR56/ADGRG1 is an adhesion G protein-coupled receptor with key roles in the nervous, reproductive, muscular, and immune systems, and blood cell production[3]. It promotes myelin sheath and oligodendrocyte formation in the cerebral cortex, aids in hematopoietic stem cell development, induces adipogenesis, and regulates immune cell function [4]. | Expressed in the tubules, however, no specific role has been proposed [5]. | *ADGRG1* WT variant- The canonical (predominant) form of *ADGRG1* with 14 exons and 693aa. Multiple AS variants in humans:<br>S1: 6 aa deleted in Ex10. No known role.<br>S2: 6 aa deleted in Ex10, however, an additional 5 aa added in Ex2. No known role.<br>S3: Partial deletion of the PLL and GAIN domains. No known role.<br>S4: Lacks the entire PPL domain and is involved in microglia-mediated synaptic pruning. Complete GAIN domain [3]. |
| *Polr1a* | RNA Polymerase I Subunit A | --- | 3.96e-15 | Subunit A is a catalytic subunit of the RNA Polymerase 1 Complex, responsible for mRNA transcription. Defects in this gene are linked to Acrofacial Dysostosis Cincinnati Type [6]. | No specific role to the kidney-has broad expression across various organs. | *POLR1A* variant 1: Canonical variant of *POLR1A* consists of 34 exons in humans (NCBI). No experimentally found splice variants. |
| *Tas1r2* | Taste receptor type 1 member 2 | --- | 6.42e-11 | Sweet taste receptor and G-protein-coupled-receptor. Dimerizes with the TAS1R3 protein. TAS1R2 provides the binding site for the sweet stimuli while the TAS1R3 protein is responsible for signal transduction and amplification [7]. | Experimentally shown expression of TAS1R2/TAS1R3 heterodimers in the kidneys. Potential renal role towards regulating amino acid and energy homeostasis [8]. | *TAS1R2* variant 1: Canonical variant of *TAS1R2* consists of 6 exons in humans (NCBI).<br>*Tas1r2* variant 2: Novel splice variant (ex4 deleted), found in mouse taste buds. Causes a decrease in the perception of sweet taste responses [9]. |
| *Tacc2* | Transforming Acidic Coiled-Coil Containing Protein 2 | 4.32e-3 | 1.66e-10 | TACC2 is associated with the centrosome-spindle apparatus that is used during the metaphase portion of mitosis. It is crucial for the proper segregation of chromosomes and thus, it is necessary for accurate cell division [10]. | Expressed in human kidney, however, no known role has been established [10]. | *TACC2* variant 1: Canonical variant; 22 exons in rats, 23 in humans. Additional AS variants identified (NCBI).<br>*Tacc2* isoform 3 is the shorter variant with 17 exons with different open reading frame shown to be expressed in developmental stages of rats, whereas variant 1 is distributed mainly in adult rat tissues [11]. |

| | | | | | | |
|---|---|---|---|---|---|---|
| *Rgs3* | Regulator of G Protein Signaling 3 | 1.33e-7 | 7.18e-10 | RGS3 plays a crucial role in regulating G protein-coupled receptor (GPCR) signaling by binding and inhibiting the activated form of Gα$_{11}$, and reducing cytoplasmic calcium levels [12]. | RGS3 is suggested to modulate tubular functions during renal development and in the adult kidney [13]. | *RGS3* variant 1: Canonical variant with 1,086 aa and 23 exons in humans (NCBI). Thirteen additional known splice variants in humans. *RGS3s* variant: Rapidly activated when calcium binds to its EF domain, causing rapid desensitization of GPCR signaling. *RGS3ss* variant: Lacks the first 125aa that are present in RGS3s. Slower activation due to lack of EF domain, causing slower desensitization of GPCR signaling [14]. |
| *Cab39* | Calcium Binding Protein 39 | --- | 2.82e-3 | Plays a role in cell signaling and regulation of protein kinases. It is a binding partner and an enhancer of SPAK/OSR1 and WNK4 activity- helping to facilitate kinase autoactivation [15]. | WNK4 and SPAK/OSR1 have been shown to play an important role in renal function. CAB39 is involved in the activation of these pathways and thus suggesting a link with kidney function [15]. | *CAB39* variant 1: Canonical isoform with 9 exons in humans. There are 2 additional known slice variants in humans (NCBI). |
| *Tjp1* | Tight Junction Protein 1 | 1.38e-4 | 2.74e-3 | TJP1/ZO1 is essential for tight junctions, which connect adjacent cells. It contains PDZ, SH3, and GuK domains, and unique motifs. It coordinates protein and actin binding for proper tight junction function [16]. | TJP1 is crucial for developing the podocyte filtration barrier. Downregulation of TJP1 impairs podocyte foot process formation and slit diaphragm creation, leading to glomerular dysfunction. It integrates epithelial junction components with newly synthesized podocyte-specific elements. [17]. | *TJP1* Variant 1: Canonical form containing 28 Exons and 1,841 aa in humans. Referred to as TJP1/ZO1 α+. Eight additional known AS variants in humans *TJP*1 Variant 2: Referred to as TJP1/ZO1 α-, and is found to be expressed in tumor tissues. This variant lacks Ex20 [18]. |
| *Lima1* | LIM Domain and Actin Binding 1 | --- | 6.41e-9 | LIMA1 organizes the actin cytoskeleton by binding and crosslinking actin filaments, promoting stability and regulating actin dynamics at the leading edge of migrating cells. It is highly expressed at adherens junctions, is essential for epithelial cell polarity [19]. | It has been shown that there is biased LIMA1 expression in the kidneys. It also modulates platelet-derived growth factor-mediated adhesion and motility of mesangial cells [20]. | *LIMA1a and b* variants: Two variants consisting of 11 exons or exons 4-11, resulting from transcription from 2 distinct promoters. The variants encode for proteins with 600 and 759 aa [19]. |
| *Ampd2* | Adenosine Monophosphate Deaminase 2 | --- | 8.27e-9 | AMPD2 catalyzes the deamination of AMP to IMP, a key step in the purine nucleotide cycle, helping to regulate the balance of purine nucleotides in the cell. [21]. | *Ampd* knockout mice show a significant increase in proteinuria, particularly albumin, suggesting that AMPD2 may play a role in glomerular filtration [22]. | *Ampd2* variant 1: Canonical variant found with 19 exons encoding for 825 amino acids in humans. Four additional AS variants known (NCBI). |

| | | | | | | |
|---|---|---|---|---|---|---|
| *Macf1* | Microtubule-actin crosslinking factor 1 | 9.91e-5 | 1.79e-8 | The MACF1 protein stabilizes the cytoskeleton by linking microtubules to actin filaments, coordinating their crosslinking and organization. It is involved in cell shape, division, intracellular transport, adhesion, and movement [23]. | No specific role to the kidney has been described. It has broad expression across various organs. | *MACF1A*: Canonical variant. Humans: 93 exons encoding 5,430 aa. Mice: makes a protein of 600 kD with 7,263 aa and 114 exons.<br>*Mac1fb*: Encodes a large (~800 kD) protein with extra plakin repeats between the plakin domain and spectrin repeats, different from the original *Macf1a*.<br>*Macf1c*: Similar to *Macf1a* but lacks the N-terminal actin-binding domain.<br>*Macf1a1*: Contains the actin-binding domain, but has a unique 5' UTR.<br>*Macf1a2*: Contains the same actin-binding domain as *Macf1a1* but a different 5' UTR.<br>*Macf1a3*: Contains only half of the actin-binding domain and a unique 5' UTR from *Macf1a1*. Contains a longer N-terminal sequence [24]. |
| *Cwc27* | Spliceosome Associated Cyclophilin | --- | 2.34e-8 | CWC27 is part of the spliceosome. As a cyclophilin, it has peptidyl prolyl isomerase activity, aiding in protein folding and stabilizing the spliceosome, contributing to its proper assembly. [25]. | No specific role to the kidney has been described. It has broad expression across various organs. | *CWC27* variant 1: Canonical variant found with 14 exons encoding 472 aa in humans (NCBI). Four additional variants known.<br>*CWC27* variant 2: Splicing variant at exon 6/intron 7 junction was observed in individuals presenting with developmental abnormalities. This variant has an altered reading frame with a premature stop codon [26]. |

*P* adj. denotes the most significant AS event observed in the gene

Supplementary Table 5: AS genes reversed with pioglitazone treatment

| Gene | most Sig *P*- adj. |
|---|---|
| *Septin 4* | 0.00000 |
| *Tmem127* | 0.00000 |
| *Col4a4* | 0.00000 |
| *Eif4g2* | 0.00001 |
| *Srpra* | 0.00001 |
| *Timp2* | 0.00001 |
| *Rpl37a* | 0.00015 |
| *Anxa2* | 0.00028 |
| *Ube2s* | 0.00031 |
| *Fstl1* | 0.00031 |
| *Clasp2* | 0.00048 |
| *Tsc22d1* | 0.00072 |
| *Ywhaq* | 0.00072 |
| *Plod1* | 0.00086 |
| *Atp6v1c1* | 0.00086 |
| *Slfn5* | 0.00100 |
| *Rps29* | 0.00116 |
| *Ap2m1* | 0.00127 |
| *Adgrl4* | 0.00137 |
| *Grina* | 0.00143 |
| *Nedd4* | 0.00143 |
| *Gask1b* | 0.00157 |
| *Rps3a* | 0.00164 |
| *Scpep1* | 0.00168 |
| *Ndufs2* | 0.00177 |
| *Abce1* | 0.00182 |
| *Myof* | 0.00196 |
| *Eef1g* | 0.00294 |
| *Mink1* | 0.00308 |
| *Atp6v1d* | 0.00328 |
| *Tjp1* | 0.00388 |
| *Fam20b* | 0.00403 |
| *Tspan8* | 0.00403 |
| *Wdr81* | 0.00535 |
| *Cd55* | 0.00535 |
| *Ncl* | 0.00535 |
| *Ctsb* | 0.00601 |
| *Col4a1* | 0.00601 |
| *Snx5* | 0.00601 |
| *Tab2* | 0.00639 |
| *Myo18a* | 0.00639 |
| *Col1a1* | 0.00639 |
| *Vim* | 0.00639 |

| | |
|---|---|
| *Actn1* | 0.00639 |
| *Mtarc1* | 0.00687 |
| *Spp1* | 0.00687 |
| *Lgalsl* | 0.00687 |
| *Rragc* | 0.00687 |
| *Ctsh* | 0.00696 |
| *Ptprm* | 0.00696 |
| *Ptprk* | 0.00704 |
| *Rgs3* | 0.00704 |
| *Aurkaip1* | 0.00883 |
| *Mylk* | 0.00888 |
| *Eif4g1* | 0.00888 |
| *Babam1* | 0.00888 |
| *Washc2c* | 0.00912 |
| *Tie1* | 0.00917 |
| *Mllt1* | 0.00917 |
| *Hnrnpk* | 0.00993 |

Supplementary Table 6: AS genes reversed with GQ-16 treatment

| **Gene** | **most Sig *P*- adj.** |
|---|---|
| *Tcea1* | 0.0000000 |
| *Bcl2l2* | 0.0000000 |
| *LOC103689975* | 0.0000022 |
| *Tns3* | 0.0000155 |
| *Mgam* | 0.0001400 |
| *Kcmf1* | 0.0009240 |
| *Icoslg* | 0.0036800 |
| *Wdr54* | 0.0038000 |
| *LOC100911837* | 0.0043100 |
| *LOC103689996* | 0.0043100 |
| *Ankrd24* | 0.0043100 |
| *Pdcd5* | 0.0088800 |
| *LOC108348065* | 0.0088800 |
| *LOC108348136* | 0.0096800 |
| *Asb2* | 0.0099500 |

Supplementary Table 7: Deconvolution of APA genes in PAN and ADR models for PEC subtypes

| | | PAN | |
|---|---|---|---|
| **Cluster** | **Gene** | **% APA Difference** | **r#** |
| PEC-A1 | *Aif1l* | 16.68 | 0.22 |
| PEC-A1 | *Cd151* | 6.15 | 0.08 |
| PEC-A1 | *Golim4* | 10.29 | 0.08 |
| PEC-A1 | *Itgb1* | 5.68 | 0.06 |
| PEC-A1 | *Nphs1* | 12.75 | 0.14 |
| PEC-A1 | *Nphs2* | 4.43 | -0.04 |
| PEC-A1 | *Podxl* | 5.05 | -0.05 |
| PEC-A1 | *Rhpn1* | 4.51 | 0.06 |
| PEC-A1 | *Sema3g* | 3.64 | 0.04 |

| | | ADR | |
|---|---|---|---|
| **Cluster** | **Gene** | **% APA Difference** | **r** |
| PEC-A1 | *Actb* | 8.529 | 0.055 |
| PEC-A1 | *Actg1* | 16.160 | 0.184 |
| PEC-A1 | *Aif1l* | 12.440 | 0.154 |
| PEC-A1 | *Cd151* | 12.249 | 0.170 |
| PEC-A1 | *Clic5* | 7.446 | 0.073 |
| PEC-A1 | *Crim1* | 13.120 | 0.096 |
| PEC-A1 | *Dpysl2* | 19.496 | 0.248 |
| PEC-A1 | *Dusp3* | 25.779 | 0.245 |
| PEC-A1 | *Efnb1* | 9.148 | 0.093 |
| PEC-A1 | *Fxyd6* | 9.932 | 0.118 |
| PEC-A1 | *Golim4* | 10.680 | 0.077 |
| PEC-A1 | *Itgb1* | 11.400 | 0.129 |
| PEC-A1 | *Klf2* | 11.878 | 0.144 |
| PEC-A1 | *Nes* | 7.095 | 0.061 |
| PEC-A1 | *Nphs1* | 24.074 | 0.248 |
| PEC-A1 | *Nphs2* | 9.179 | -0.047 |
| PEC-A1 | *Plat* | 7.149 | 0.075 |
| PEC-A1 | *Podxl* | 7.678 | -0.091 |
| PEC-A1 | *Prss23* | 19.624 | 0.175 |
| PEC-A1 | *Sdc2* | 12.089 | 0.109 |
| PEC-A1 | *Sema3g* | 9.305 | 0.105 |
| PEC-A1 | *Slc9a3r2* | 4.775 | 0.043 |
| PEC-A1 | *Trib2* | 19.211 | 0.158 |
| PEC-A1 | *Tspan13* | 11.317 | 0.089 |
| PEC-A1 | *Wt1* | 10.310 | 0.118 |

| Cluster | Gene | % APA Difference | r |
|---|---|---|---|
| PEC-A2 | N/A | 0.00 | |

| Cluster | Gene | % APA Difference | r |
|---|---|---|---|
| PEC-A2 | N/A | 0 | |

| Cluster | Gene | % APA Difference | r |
|---|---|---|---|
| PEC-A3 | *Tuba1c* | 4.82 | -0.07 |
| PEC-A3 | *Spc25* | 49.97 | -0.49 |

| Cluster | Gene | % APA Difference | r |
|---|---|---|---|
| PEC-A3 | *Ube2s* | 11.452 | -0.119 |
| PEC-A3 | *Tuba1c* | 6.259 | -0.068 |
| PEC-A3 | *Ywhah* | 11.416 | 0.111 |
| PEC-A3 | *Hmgn2* | 7.874 | 0.111 |
| PEC-A3 | *Pkm* | 6.088 | 0.061 |

| Cluster | Gene | % APA Difference | r |
|---|---|---|---|
| N/A | | 0.00 | |

| Cluster | Gene | % APA Difference | r |
|---|---|---|---|
| PEC-A4 | *Aldh9a1* | 22.487 | -0.182 |
| PEC-A4 | *Aldob* | 3.213 | 0.033 |
| PEC-A4 | *Aqp1* | 6.608 | 0.099 |
| PEC-A4 | *Atp1a1* | 4.297 | 0.052 |
| PEC-A4 | *Cat* | 9.359 | 0.098 |
| PEC-A4 | *Cd74* | 7.894 | 0.087 |
| PEC-A4 | *Cldn2* | 22.477 | 0.145 |
| PEC-A4 | *Cox5a* | 14.373 | -0.215 |
| PEC-A4 | *Dab2* | 11.203 | 0.115 |
| PEC-A4 | *Fth1* | 4.469 | 0.048 |
| PEC-A4 | *Gatm* | 5.253 | 0.045 |
| PEC-A4 | *Gpx1* | 3.679 | 0.042 |
| PEC-A4 | *Gpx3* | 9.089 | 0.086 |
| PEC-A4 | *Hao2* | 6.643 | 0.081 |
| PEC-A4 | *Id2* | 14.629 | 0.146 |
| PEC-A4 | *Kcnj15* | 19.135 | 0.205 |
| PEC-A4 | *Lrp2* | 12.690 | 0.128 |
| PEC-A4 | *Mep1a* | 16.236 | 0.182 |
| PEC-A4 | *Ndrg1* | 5.834 | 0.085 |
| PEC-A4 | *Pck1* | 10.915 | 0.132 |
| PEC-A4 | *Rida* | 4.995 | 0.055 |
| PEC-A4 | *Slc13a3* | 13.353 | 0.137 |
| PEC-A4 | *Slc34a1* | 3.402 | 0.036 |
| PEC-A4 | *Slc4a4* | 23.658 | 0.231 |
| PEC-A4 | *Tpi1* | 8.286 | 0.095 |

| Cluster | Gene | % APA Difference | r |
|---|---|---|---|
| PEC-B | *Col4a1* | 8.99 | 0.09 |
| PEC-B | *Igfbp5* | 8.12 | 0.11 |
| PEC-B | *Epas1* | 5.71 | 0.07 |
| PEC-B | *Cst3* | 2.90 | -0.02 |

| Cluster | Gene | % APA Difference | r |
|---|---|---|---|
| PEC-B | *Apoe* | 12.323 | 0.150 |
| PEC-B | *Bok* | 23.466 | 0.280 |
| PEC-B | *Col4a1* | 7.557 | 0.083 |
| PEC-B | *Cx3cl1* | 30.029 | 0.439 |
| PEC-B | *Dkk2* | 11.552 | 0.131 |
| PEC-B | *Emid1* | 10.839 | 0.118 |
| PEC-B | *Epas1* | 14.055 | 0.156 |
| PEC-B | *Errfi1* | 16.184 | 0.162 |
| PEC-B | *F2r* | 18.115 | 0.135 |
| PEC-B | *Fn1* | 16.529 | 0.207 |
| PEC-B | *Fstl1* | 6.201 | 0.083 |
| PEC-B | *Igfbp5* | 14.413 | 0.105 |
| PEC-B | *Il6st* | 14.064 | 0.186 |
| PEC-B | *Lbh* | 10.660 | 0.234 |
| PEC-B | *Lpl* | 13.083 | 0.137 |
| PEC-B | *Lrp1* | 17.180 | 0.218 |
| PEC-B | *Man2a1* | 24.289 | 0.271 |
| PEC-B | *Marcksl1* | 14.970 | 0.219 |
| PEC-B | *Mfge8* | 8.606 | 0.077 |
| PEC-B | *Mgp* | 3.161 | 0.031 |
| PEC-B | *Mmd* | 26.095 | 0.423 |
| PEC-B | *Myo1b* | 13.062 | 0.161 |
| PEC-B | *Nrp1* | 21.741 | 0.183 |
| PEC-B | *Oaf* | 15.748 | 0.206 |
| PEC-B | *Scarb2* | 19.952 | 0.137 |
| PEC-B | *Serpine2* | 14.243 | 0.185 |
| PEC-B | *Tril* | 32.300 | 0.292 |

*[#] '+ r value denotes distal shift and - r value denotes proximal shift*

**Supplementary Table 8:** APA genes in PAN and ADR models represented in monogenic genes with established roles in glomerular disease

| Gene | Protein | APA (%) | | DEG (Log2FC) | | Function | Podocyte/Kidney Role | Known APA | Targeting miRs# (score) | miR Sequence |
|---|---|---|---|---|---|---|---|---|---|---|
| | | PAN | ADR | PAN | ADR | | | | | |
| *Podxl* | Podocalyxin | 50.45 (P) | 7.68 (P) | 1.03 | 1.56 | It is highly expressed on the surface of podocytes and plays a crucial role in the formation and maintenance of podocyte foot processes. Additionally, it contributes to tissue remodeling and development [1]. | Mutations in this gene are linked to proteinuria-related kidney disease, with some leading to anuria (failure to produce urine). Podocalyxin shedding in urine is a common biomarker for nephrotic syndrome [2]. | --- | hsa-miR-6748-3p (93)<br>hsa-miR-3617-5p (87)<br>hsa-miR-4319 (87)<br>hsa-miR-641 (87)<br>hsa-miR-6759-3p (87)<br>hsa-miR-4651 (85)<br>hsa-miR-608 (85)<br>hsa-miR-3925-3p (84)<br>hsa-miR-3613-3p (84)<br>hsa-miR-517-5p (83) | uccugucccugucuccuacag<br>aaagacauaguugcaagauggg<br>ucccugagcaaagccac<br>aaagacauaggauagagucaccuc<br>ugaccuuugccucuccccucag<br>cggggugggugaggucgggc<br>agggguggguguugggacagcuccgu<br>acuccaguuuuaguucucuug<br>acaaaaaaaaaagcccaacccuuc<br>ccucuagauggaagcacugucu |
| *Myh9* | Myosin-9 | 4.39 (D) | 10.94 (D) | 0.67 | 0.90 | A subunit of myosin IIA, crucial for cell movement, shape, and cytokinesis [3]. | Mutations are linked to nephropathy and proteinuria. SNPs in the *MYH9* gene are associated with CKD, particularly glomerulopathies, in African Americans [4]. | *a* [5] | hsa-miR-107 (96)<br>hsa-miR-103a-3p (96)<br>hsa-miR-6825-5p (96)<br>hsa-miR-3714 (96)<br>hsa-miR-3120-3p (95)<br>hsa-miR-10526-3p (95)<br>hsa-miR-4753-3p (94)<br>hsa-miR-492 (93)<br>hsa-miR-542-3p (92)<br>hsa-miR-433-3p (92) | agcagcauuguacagggcuauca<br>agcagcauuguacagggcuauga<br>uggggaggugugggagucagcau<br>gaaggcagcagugcuccccugu<br>cacagcaaguguagacaggca<br>aaaagggggcugagguggag<br>uucucuuucuuuagccuugugu<br>aggaccugcgggacaagauucuu<br>ugugacagauugauaacugaaa<br>aucaugaugggcuccucggugu |
| *Dlc1* | Rho GTPase-activating Protein 7 | | 13.41 (D) | -0.08 | 0.03 | A tumor suppressor involved in cell polarity, proliferation, migration, and survival, often downregulated in various cancer types [6]. | Mutations in this gene have been identified in nephrotic syndrome. Knockdown in podocytes reduces migration and affects the filtration barrier, while dexamethasone blocks RhoA activation in *DLC1* knockdown [7] | --- | hsa-miR-141-5p (98)<br>hsa-miR-4500 (97)<br>hsa-miR-202-3p (97)<br>hsa-let-7f-5p (96)<br>hsa-miR-4789-3p (96)<br>hsa-miR-98-5p (96)<br>hsa-miR-7162-3p (96)<br>hsa-let-7b-5p (96)<br>hsa-let-7g-5p (96)<br>hsa-miR-141-3p (96) | caucuuccaguacaguguugga<br>ugagguaguaguuucuu<br>agagguauagggcaugggaa<br>ugagguaguagauuguauaguu<br>cacacauagcagguguauaua<br>ugagguaguaaguuguauuguu<br>ucugagguggaacagcagc<br>ugagguaguagguugugugguu<br>ugagguaguaguuuguacaguu<br>uaacacugucugguaaagaugg |
| *Scarb2* | Scavenger Receptor Class B Member 2 | | 19.95 (D) | 0.03 | 0.20 | Located on the lysosomal membrane, it is essential for the biogenesis and maintenance of lysosomes and endosomes [8]. | Mutations in *SCARB2* cause action myoclonus-renal failure syndrome, leading to neurological disease and steroid-resistant nephrotic syndrome. Deficient mice show glomerular lesions | b [10] | hsa-miR-8054 (95)<br>hsa-miR-548k (95)<br>hsa-miR-548av-5p (95)<br>hsa-miR-4796-3p (91)<br>hsa-miR-340-5p (91)<br>hsa-miR-642a-3p (88)<br>hsa-miR-642b-3p (88)<br>hsa-miR-1912-5p (87) | gaaaguacagaucggaugggu<br>aaaaguacuugcggauuuugcu<br>aaaaguacuugcggauuu<br>uaaaguggcagaguauagacac<br>uuauaaagcaaugagacugauu<br>agacacauuuggagagggaacc<br>agacacauuuggagagggaccc<br>cucauugcaugggcuguguaua |

| | | | | | | | | | | |
|---|---|---|---|---|---|---|---|---|---|---|
| | | | | | | | and foot process effacement [9]. | | hsa-miR-6755-5p (86)<br>hsa-miR-339-5p (85) | uaggguagacacugacaacguu<br>ucccuguccuccaggagcucacg |
| *Wt1* | Wilm's Tumor 1 | | 10.31 (D) | 1.53 | 1.23 | WT1 is a tumor suppressor and transcription factor. Mutations, found in 10% of Wilm's tumors and other cancers, affect gene transcription, mRNA processing, and can cause congenital abnormalities[11]. | *Wt1* knockout mice show delayed kidney development. Low WT1 expression causes glomerulonephritis, mesangial sclerosis, and downregulation of podocalyxin and Nephrin [12]. | c [13] | hsa-miR-8485 (100)<br>hsa-miR-29a-5p (97)<br>hsa-miR-661 (95)<br>hsa-miR-522-3p (94)<br>hsa-miR-224-3p (94)<br>hsa-miR-5585-3p (91)<br>hsa-miR-498-5p (91)<br>hsa-miR-1910-5p (91)<br>hsa-miR-153-5p (90)<br>hsa-miR-4672 (90) | cacacacacacacacacguau<br>acugauuucuuuugguguucag<br>ugccugggucucuggccugcgcgu<br>aaaugguucccuuuagagugu<br>aaaauggugcccuagugacuaca<br>cugaauagcugggacuacaggu<br>uuucaagccaggggcguuuuuc<br>ccaguccugugccugccgccu<br>ucauuuuugugauguugcagcu<br>uuacacagcuggacagaggca |
| *Kirrel1* | Kirre Like Nephrin Family Adhesion Molecule 1 | 13.88 (D) | 13.31 (D) | 1.69 | 1.66 | KIRREL1, structurally similar to nephrin, forms the slit diaphragm in the glomerular filtration barrier with nephrin and podocin. It may regulate the HIPPO pathway and suppress YAP activity [14]. | KIRREL1, with nephrin and podocin, maintains slit diaphragm integrity and filtering function. Its deletion in mice causes podocyte foot process effacement [15]. | --- | hsa-miR-4795-3p (100)<br>hsa-miR-5011-5p (100)<br>hsa-miR-190a-3p (100)<br>hsa-miR-1277-5p (99)<br>hsa-miR-126-5p (99)<br>hsa-miR-5692c (98)<br>hsa-miR-5692b (98)<br>hsa-miR-765 (96)<br>hsa-miR-3140-5p (96)<br>hsa-miR-11181-3p (95) | auauuauuagccacuucuggau<br>uauauauacagccaugcacuc<br>cuauauaucaaacauauuccu<br>aaauauauauauauauguacguau<br>cauuauuacuuuugguacgcg<br>aauaauaucacaguagguguac<br>aauaauaucacaguaggugu<br>uggaggagaaggaaggugaug<br>accugaauuaccaaaagcuuu<br>aggaggaggaggucaggc |
| *Nphs1* | Nephrin | 12.75 (D) | 24.07 (D) | 1.18 | 0.87 | Podocyte surface protein, crucial for slit diaphragm formation and maintenance [16]. | Mutations in this gene cause nephrotic syndrome, proteinuria, and end-stage renal failure. Low nephrin leads to proteinuria, glomerular hypertrophy, and FSGS. *Nphs*1 knockout mice can't recover from foot process effacement [17]. | *d* [18] | hsa-miR-4774-3p (85)<br>hsa-miR-10524-5p (83)<br>hsa-miR-4436b-5p (83)<br>hsa-miR-4516 (80)<br>hsa-miR-4729 (78)<br>hsa-miR-1286 (78)<br>hsa-miR-4795-3p (78)<br>hsa-miR-5703 (77)<br>hsa-miR-4434 (77)<br>hsa-miR-205-3p (77) | auugccuaacaugugccagaa<br>caggaugccagcauagu<br>guccacuucugccugcccugcc<br>gggagaagggucggggc<br>ucauuuaucuguugggaagcua<br>ugcaggaccaagaugagcccu<br>auauuauuagccacuucuggau<br>aggagaagucgggaaggu<br>aggagaaguaaaguagaa<br>gauuucaguggagugaaguuc |
| *Nphs2* | Podocin | 4.43 (P) | 9.18 (P) | 2.61 | 2.65 | Podocin is a membrane protein almost exclusively expressed in podocytes, essential for maintaining slit diaphragm integrity [19]. | Mutations cause proteinuria and congenital nephrotic syndrome. Podocin interacts with nephrin in the slit diaphragm. Knockout mice | e [19] | hsa-miR-196a-3p (90)<br>hsa-miR-6823-5p (90)<br>hsa-miR-494-3p (88)<br>hsa-miR-3915 (87)<br>hsa-miR-4306 (85)<br>hsa-miR-6805-3p (82)<br>hsa-miR-5691 (82)<br>hsa-miR-3164 (82)<br>hsa-miR-6820-3p (82)<br>hsa-miR-146a-3p (78) | cggcaacaagaaacugccugag<br>ucaggguugguaggggguugcu<br>ugaaacauacacgggaaaccuc<br>uugaggaaaagauggucuuauu<br>uggagagaaaggcagua<br>uugcucugcucccccgcccccag<br>uugcucugagcuccgagaaagc<br>ugugacuuuaagggaaauggcg<br>ugugacuucuccccugccacag<br>ccucugaaauucaguucuucag |

| | | | | | | | | | | |
|---|---|---|---|---|---|---|---|---|---|---|
| | | | | | | | show proteinuria at birth [20]. | | | |
| *Cd151* | Cluster of Differentiation 151 | 6.15 (D) | 12.25 (D) | 0.99 | 1.40 | CD151, a tetraspanin family protein, is highly expressed on cell surfaces and vesicles. It forms webs with membrane receptors and immunoglobulins, acting as a molecular facilitator [21]. | CD151 promotes podocyte adhesion to the GBM, preventing glomerular disease under high capillary pressure. Mice lacking *Cd151* develop renal injury after severe hypertension. [22]. | --- | hsa-miR-506-3p (93)<br>hsa-miR-124-3p (93)<br>hsa-miR-1207-5p (91)<br>hsa-miR-3663-3p (91)<br>hsa-miR-4763-3p (88)<br>hsa-miR-4797-3p (82)<br>hsa-miR-199b-3p (78)<br>hsa-miR-199a-3p (78)<br>hsa-miR-199a-3p (78)<br>hsa-miR-4731-5p (76) | uaaggcacccuucugaguaga<br>uaaggcacgcggugaaugccaa<br>uggcagggaggcugggagggg<br>ugagcaccacacaggccgggcgc<br>aggcaggggcuggugcugggcggg<br>ucucaguaaguggcacucugu<br>acaguagucugcacauugguua<br>acaguagucugcacauugguua<br>gcaguaguguagagauugguuu<br>ugcugggggccacaugagugug |
| *Thbd* | Thrombomodulin | | 14.17 (D) | -0.24 | 0.09 | It has anticoagulant and anti-inflammatory properties [23]. | Thrombomodulin protects the glomerular filtration barrier by enabling communication between glomerular endothelium and podocytes. Its loss in the glomerulus leads to glomerulopathies [24]. | --- | hsa-miR-9902 (95)<br>hsa-let-7c-3p (91)<br>hsa-miR-1225-3p (91)<br>hsa-miR-550a-3p (91)<br>hsa-miR-200c-5p (90)<br>hsa-miR-7159-3p (90)<br>hsa-miR-4299 (89)<br>hsa-miR-122b-5p (89)<br>hsa-miR-7106-5p (87)<br>hsa-miR-5692a (86) | cccagaaaucugguaugccagc<br>cuguacaaccuucuagcuuucc<br>ugagccccugugccgcccccag<br>ugucuuacucccucaggcacau<br>cgucuuacccagcaguguuugg<br>uuucuauguuaguuggaag<br>gcuggugacaugagaggc<br>uuuagugugauaauggcguuuga<br>ugggaggaggggaucuuggg<br>caaauaauaccacagugggugu |
| *Lmx1b* | LIM Homeobox Transcription Factor 1 Beta | 10.18 (D) | | 1.59 | 1.32 | LMX1B, a LIM-homeodomain transcription factor, regulates body patterning. It's expressed in dorsal mesenchyme, and knockout mice lack dorsal limb structures, duplicating ventral ones [25]. | LMX1B is crucial for maintaining actin cytoskeletal integrity in podocytes. Podocyte-specific deletion in mice causes post-natal death around 14 days after birth [26]. | --- | hsa-miR-6867-5p (100)<br>hsa-miR-6752-3p (95)<br>hsa-miR-4731-5p (91)<br>hsa-miR-4505 (91)<br>hsa-miR-3085-3p (91)<br>hsa-miR-5787 (90)<br>hsa-miR-4288 (90)<br>hsa-miR-3064-5p (89)<br>hsa-miR-6504-5p (89)<br>hsa-miR-6721-5p (86) | ugugugugguagaggaagaaggga<br>ucccugcccccauacucccag<br>ugcugggggccacaugagugug<br>aggcugggcugggacgga<br>ucuggcugcuauggcccccuc<br>gggcuggggcgcggggaggu<br>uugucugcugaguuucc<br>ucuggcuguuguggugugcaa<br>ucuggcugugcuguaaugcag<br>ugggcaggggcuuauuguaggag |
| *Rac1* | Rac Family Small GTPase 1 | | 5.86 (D) | 0.21 | 0.24 | RAC1, a Rho family GTPase, is found at the leading edge of migrating cells and regulates pathways involved in cell proliferation, transcription, and cytoskeleton organization [27]. | Hyperactivation of Rac1 in glomerular podocytes is linked to proteinuric kidney diseases, while Rac1 deficiency can impair post-natal development of the renal medulla [28]. | f [29] | hsa-miR-4715-5p (96)<br>hsa-miR-101-3p (96)<br>hsa-miR-6835-3p (95)<br>hsa-miR-3688-3p (94)<br>hsa-miR-142-3p (93)<br>hsa-miR-3686 (92)<br>hsa-miR-3148 (92)<br>hsa-miR-2909 (91)<br>hsa-miR-659-3p (90)<br>hsa-miR-548x-3p (89) | aaguuggcugcaguuaaggugg<br>uacaguacugugauaacugaa<br>aaaagcacuuuucugucucccag<br>uauggaaagacuuugccacucu<br>uguaguguuuccuacuuuaugga<br>aucuguaagagaaaguaaauga<br>uggaaaaaacuggugugugcuu<br>guuagggccaacaucucuugg<br>cuugguucagggagggucccca<br>uaaaaacugcaauuacuuuc |

| | | | | | | | | | | |
|---|---|---|---|---|---|---|---|---|---|---|
| *Sgpl1* | Sphingo sine-1-phosph ate Lyase 1 | | 17.66 (D) | 0.52 | 0.50 | SGPL1 encodes sphingosine-1-phosphate lyase, an enzyme in the endoplasmic reticulum responsible for the final step of sphingolipid breakdown, irreversibly cleaving sphingosine-1-phosphate [30]. | Biallelic mutations in *SGPL1* cause a severe form of steroid-resistant nephrotic syndrome. *SGPL1* knockdown downregulates nephrin and WT1, potentially leading to podocyte foot process effacement and albuminuria. [31]. | g [32] | hsa-miR-8485 (96)<br>hsa-miR-544b (6)<br>hsa-miR-4319 (95)<br>hsa-miR-4478 (95)<br>hsa-miR-4747-5p (94)<br>hsa-miR-6867-5p (94)<br>hsa-miR-5196-5p (94)<br>hsa-miR-6847-5p (93)<br>hsa-miR-125a-5p (93)<br>hsa-miR-125b-5p (93) | cacacacacacacacacguau<br>accugagguugugcauuucuaa<br>ucccugagcaaagccac<br>gaggcugagcugaggag<br>agggaaggaggcuuggucuuag<br>ugugugguguagaggaagaaggga<br>agggaaggggacgaggguuggg<br>acagaggacaguggagugugagc<br>ucccugagacccuuuaaccuguga<br>ucccugagacccuaacuuguga |

*a; MYH9* is associated with APA in the 3'UTR of fibroblasts in quiescence after 7 days of contact inhibition when compared to proliferating fibroblasts [5]

b; APA was found in the isolated human homolog of rat *Limp2* in the 3'UTR [10]

c; Shortened isoforms of genes caused by APA events have related to tumor formation [13]

d; *NPHS1* as well as its antisense transcript have both shown multiple polyadenylation sites [18]

e; *NPHS2* contains an atypical polyadenylation signal that is 13 base pairs upstream of where the poly(A) tail attaches. There are two other overlapping polyadenylation signals in the expressed sequence tag AI672038, 44 base pairs downstream of this signal [19]

f; Rac1 exists as two different mRNA isoforms through two common poly (A) signals in the 3'UTR, in differentiated oligodendrocyte. During development, the longer mRNA isoform expression increases, and it localizes in the neurites of primary cortical neurons. Knockdown of the longer mRNA led to changes in their morphology [29]

g; SGPL1 along with CREG1 contributes to tumor progression in clear cell renal cell carcinoma via the binding of poly(A) binding protein nuclear 1 (PABPN1) and differential usage of distal vs proximal poly (A) site in its 3'UTR [32]

#; For each APA gene, predicted miRNA sites were identified from miRNA database (miRDB.org). A target score of 74 or above was selected (listed in parenthesis), and at maximum ten miRs with the highest target scores are listed for each gene. For each miR, its sequence is listed as well.

(P) denotes proximal shift and (D) denotes distal shift in the usage of poly(A) site

<u>References:</u>

7. Ashraf S, Kudo H, Rao J, Kikuchi A, Widmeier E, Lawson JA, Tan WZ, Hermle T, Warejko JK, Shril S, Airik M, Jobst-Schwan T, Lovric S, Braun DA, Gee HY, Schapiro D, Majmundar AJ, Sadowski CE, Pabst WL, Daga A, van der Ven AT, Schmidt JM, Low BC, Gupta AB, Tripathi BK, Wong J, Campbell K, Metcalfe K, Schanze D, Niihori T, Kaito H, Nozu K, Tsukaguchi H, Tanaka R, Hamahira K, Kobayashi Y, Takizawa T, Funayama R, Nakayama K, Aoki Y, Kumagai N, Iijima K, Fehrenbach H, Kari JA, El Desoky S, Jalalah S, Bogdanovic R, Stajic N, Zappel H, Rakhmetova A, Wassmer SR, Jungraithmayr T, Strehlau J, Kumar AS, Bagga A, Soliman NA, Mane SM, Kaufman L, Lowy DR, Jairajpuri MA, Lifton RP, Pei Y, Zenker M, Kure S, Hildebrandt F. Mutations in six nephrosis genes delineate a pathogenic pathway amenable to treatment. Nature Communications. 2018;9. doi: ARTN 196010.1038/s41467-018-04193-w. PubMed PMID: WOS:000432343700002.
8. Gonzalez A, Valeiras M, Sidransky E, Tayebi N. Lysosomal integral membrane protein-2: a new player in lysosome-related pathology. Mol Genet Metab. 2014;111(2):84-91. Epub 20131211. doi: 10.1016/j.ymgme.2013.12.005. PubMed PMID: 24389070; PMCID: PMC3924958.
9. Lowik MM, Groenen PJ, Levtchenko EN, Monnens LA, van den Heuvel LP. Molecular genetic analysis of podocyte genes in focal segmental glomerulosclerosis--a review. Eur J Pediatr. 2009;168(11):1291-304. Epub 20090627. doi: 10.1007/s00431-009-1017-x. PubMed PMID: 19562370; PMCID: PMC2745545.
10. Heybrock S, Kanerva K, Meng Y, Ing C, Liang A, Xiong ZJ, Weng XL, Kim YA, Collins R, Trimble W, Pomès R, Privé GG, Annaert W, Schwake M, Heeren J, Lüllmann-Rauch R, Grinstein S, Ikonen E, Saftig P, Neculai D. Lysosomal integral membrane protein-2 (LIMP-2/SCARB2) is involved in lysosomal cholesterol export. Nature Communications. 2019;10. doi: ARTN 352110.1038/s41467-019-11425-0. PubMed PMID: WOS:000478867500006.
11. Scharnhorst V, van der Eb AJ, Jochemsen AG. WT1 proteins: functions in growth and differentiation. Gene. 2001;273(2):141-61. doi: 10.1016/s0378-1119(01)00593-5. PubMed PMID: 11595161.
12. Guo JK, Menke AL, Gubler MC, Clarke AR, Harrison D, Hammes A, Hastie ND, Schedl A. WT1 is a key regulator of podocyte function: reduced expression levels cause crescentic glomerulonephritis and mesangial sclerosis. Hum Mol Genet. 2002;11(6):651-9. doi: DOI 10.1093/hmg/11.6.651. PubMed PMID: WOS:000174654300005.
13. Xue Z, Warren RL, Gibb EA, MacMillan D, Wong J, Chiu R, Hammond SA, Yang C, Nip KM, Ennis CA, Hahn A, Reynolds S, Birol I. Recurrent tumor-specific regulation of alternative polyadenylation of cancer-related genes. BMC Genomics. 2018;19(1):536. Epub 20180713. doi: 10.1186/s12864-018-4903-7. PubMed PMID: 30005633; PMCID: PMC6045855.
14. Paul A, Annunziato S, Lu B, Sun T, Evrova O, Planas-Paz L, Orsini V, Terracciano LM, Charlat O, Loureiro ZY, Ji L, Zamponi R, Sigoillot F, Lei H, Lindeman A, Russ C, Reece-Hoyes JS, Nicholson TB, Tchorz JS, Cong F. Cell adhesion molecule KIRREL1 is a feedback regulator of Hippo signaling recruiting SAV1 to cell-cell contact sites. Nat Commun. 2022;13(1):930. Epub 20220217. doi: 10.1038/s41467-022-28567-3. PubMed PMID: 35177623; PMCID: PMC8854406.
15. Solanki AK, Widmeier E, Arif E, Sharma S, Daga A, Srivastava P, Kwon SH, Hugo H, Nakayama M, Mann N, Majmundar AJ, Tan W, Gee HY, Sadowski CE, Rinat C, Becker-Cohen R, Bergmann C, Rosen S, Somers M, Shril S, Huber TB, Mane S, Hildebrandt F, Nihalani D. Mutations in KIRREL1, a slit diaphragm component, cause steroid-resistant nephrotic syndrome. Kidney Int. 2019;96(4):883-9. Epub 20190710. doi: 10.1016/j.kint.2019.06.016. PubMed PMID: 31472902; PMCID: PMC6756928.
16. Martin CE, Jones N. Nephrin Signaling in the Podocyte: An Updated view of Signal Regulation at the Slit Diaphragm and Beyond. Front Endocrinol. 2018;9. doi: ARTN 30210.3389/fendo.2018.00302. PubMed PMID: WOS:000434129600001.
17. Zhuo L, Huang LL, Yang ZL, Li GS, Wang L. A comprehensive analysis of NPHS1 gene mutations in patients with sporadic focal segmental glomerulosclerosis. Bmc Medical Genetics. 2019;20. doi: ARTN 11110.1186/s12881-019-0845-4. PubMed PMID: WOS:000472141600001.
18. Ihalmo P, Rinta-Valkama J, Mai P, Astrom E, Palmen T, Pham TT, Floss T, Holthofer H. Molecular cloning and characterization of an endogenous antisense transcript of Nphs1. Genomics. 2004;83(6):1134-40. doi: 10.1016/j.ygeno.2004.01.001. PubMed PMID: 15177566.
19. Boute N, Gribouval O, Roselli S, Benessy F, Lee H, Fuchshuber A, Dahan K, Gubler MC, Niaudet P, Antignac C. , encoding the glomerular protein podocin, is mutated in autosomal recessive steroid-resistant nephrotic syndrome. Nat Genet. 2000;24(4):349-54. doi: Doi 10.1038/74166. PubMed PMID: WOS:000086192800011.
20. Franceschini N, North KE, Kopp JB, Mckenzie L, Winkler C. gene, nephrotic syndrome and focal segmental glomerulosclerosis:: A HuGE review. Genetics in Medicine. 2006;8(2):63-75. doi: 10.1097/01.gim.0000200947.09626.1c. PubMed PMID: WOS:000235504400001.
21. Kumari S, Devi Gt, Badana A, Dasari VR, Malla RR. CD151-A Striking Marker for Cancer Therapy. Biomark Cancer. 2015;7:7-11. Epub 20150322. doi: 10.4137/BIC.S21847. PubMed PMID: 25861224; PMCID: PMC4372031.
22. Pozzi A, Zent R. Hold tight or you'll fall off: CD151 helps podocytes stick in high-pressure situations. J Clin Invest. 2012;122(1):13-6. Epub 20111227. doi: 10.1172/JCI61858. PubMed PMID: 22201676; PMCID: PMC3248315.
23. Okamoto T, Tanigami H, Suzuki K, Shimaoka M. Thrombomodulin: a bifunctional modulator of inflammation and coagulation in sepsis. Crit Care Res Pract. 2012;2012:614545. Epub 20120228. doi: 10.1155/2012/614545. PubMed PMID: 22482044; PMCID: PMC3299293.

**Supplementary Table 9:** APA genes in PAN and ADR models representing novel genes in glomerular disease

| Genes | Protein | APA (%) | | DEG (Log2FC) | | Function | Podocyte/Kidney Role | Known APA | Targeting miRs# (score) | miR Sequence |
|---|---|---|---|---|---|---|---|---|---|---|
| | | PAN | ADR | PAN | ADR | | | | | |
| *Hsd17b14* \|1\|+ | Hydroxysteroid (17-beta) Dehydrogenase 14 | | 45.3 (P) | -0.18 | -0.34 | A 17-beta-hydroxysteroid dehydrogenase enzyme, catalyzing NADPH-dependent reactions on androgens and estrogens, highly expressed in brain and kidney, with a possible role in retinoid metabolism [1]. | Low expression in the glomeruli but high in proximal tubules, which decreases in diabetic kidney disease and mouse models. Genome-wide SNP analysis shows a protective allele against end-stage kidney disease in Type 1 Diabetes [2]. | --- | hsa-miR-378g (74) | acugggcuuggagucagaag |
| *Dph2* \|5\|- | Diphthamide Biosynthesis 2 | | 41.4 (D) | 0.16 | 0.13 | DPH1 is involved in diphthamide biosynthesis in elongation factor 2. DPH1 deficiency causes diphthamide-deficiency syndrome, leading to developmental delays, abnormal head size, short stature, and congenital heart disease [3]. | Diphthamide biosynthesis genes, including *DPH2*, have been shown to regulate podocyte adhesion. Knockdown or knockout of *DPH1*, *DPH2*, *DPH3*, and *DPH4* increases podocyte adhesion and causes spreading defects, which may indicate a failure in process formation at the leading edge [4]. | --- | hsa-miR-7113-3p (95)<br>hsa-miR-4319 (93)<br>hsa-miR-597-3p (92)<br>hsa-miR-125b-5p (91)<br>hsa-miR-125a-5p (91)<br>hsa-miR-539-5p (88)<br>hsa-miR-29b-1-5p (88)<br>hsa-miR-874-5p (86)<br>hsa-miR-3679-3p (82)<br>hsa-miR-134-5p (80) | ccucccugcccgccucucugcag<br>ucccugagcaaagccac<br>ugguucucuuguggcucaagcgu<br>ucccugagacccuaacuuguga<br>ucccugagacccuuuaaccuguga<br>ggagaaauuauccuuggugugu<br>gcugguuucauauggugguuuaga<br>cggccccacgcaccagguaaga<br>cuuccccccaguaaucuucauc<br>ugugacugguugaccagagggg |
| *Dvl3* \|11\|- | Disheveled Segment Polarity Protein 3 | | 40.6 (D) | 0.41 | -0.02 | *DVL3*, along with other Disheveled genes *DVL1* and *DVL2*, functions as a key component of the Wnt signaling pathway. These genes regulate various cellular processes, including cell proliferation, survival, differentiation, and polarity [5] | Dvl3 is a key component of the Wnt signaling pathway, which plays a critical role in kidney development. Wnt signaling has been implicated in the development of renal fibrosis, and alterations in this pathway may also be associated with diabetic nephropathy [6]. | --- | hsa-miR-4731-5p (97)<br>hsa-miR-4311 (96)<br>hsa-miR-6742-5p (95)<br>hsa-miR-3188 (93)<br>hsa-miR-1294 (93)<br>hsa-miR-9986 (93)<br>hsa-miR-1275 (92)<br>hsa-miR-2467-5p (92)<br>hsa-miR-4651 (92)<br>hsa-miR-608 (92) | ugcuggggccacaugagugu<br>gaaagagagcugagugug<br>aguggggugggacccagcuguu<br>agaggcuuugugcggauacgggg<br>ugugagguuggcauuguugucu<br>ugugagguugucaugccugc<br>gugggggagaggcuguc<br>ugaggcucuguuagccuuggcuc<br>cggggugggugaggucgggc<br>aggggugguguugggacagcuccgu |
| *Ccr5* \|8\|+ | C-C Motif Chemokine Receptor 5 | | 39.5 (D) | 0.95 | 1.12 | CCR5 is a receptor for the C-C chemokine RANTES, regulating the trafficking and functions of memory/effector T-lymphocytes, macrophages, and immature dendritic cells. It also acts as a coreceptor for R5 strains of HIV-1 and HIV-2 [7]. | CCR5-positive cells are elevated in patients with impaired renal function, seen in chronic glomerulonephritis, interstitial nephritis, transplant rejection, lupus nephritis, and membranoproliferative glomerulonephritis. [8]. | --- | hsa-miR-320a-5p (92)<br>hsa-miR-3179 (92)<br>hsa-miR-498-5p (91)<br>hsa-miR-670-5p (87)<br>hsa-miR-5193 (85)<br>hsa-miR-1202 (85)<br>hsa-miR-3972 (85)<br>hsa-miR-27a-5p (83)<br>hsa-miR-4319 (82)<br>hsa-miR-629-3p (76) | gccuucucuucccgguucuucc<br>agaaggggugaaauuuaaacgu<br>uuucaagccaggggcguuuuuc<br>gucccugaguguauguggug<br>uccuccucuaccucaucccagu<br>gugccagcugcaguggggggag<br>cugccagccccguuccagggca<br>agggcuuagcugcuugugagca<br>ucccugagcaaagccac<br>guucucccaacguaagcccagc |
| *Foxn3* \|6\|- | Forkhead Box N3 | 36.6 (D) | 39.5 (D) | 0.16 | 0.08 | FOXN3, a forkhead/winged helix transcription factor, regulates the cell cycle and tumorigenesis. It acts as a tumor suppressor, with alterations linked to cancers | FOXN3 overexpression is reported to repress the AKT/MDM2/p53 signaling pathway in human glioma cells. This pathway is involved in fibroblast growth factor 21 | a [11] | hsa-miR-3119 (100)<br>hsa-miR-6867-5p (100)<br>hsa-miR-450b-5p (100)<br>hsa-miR-135b-5p (98)<br>hsa-miR-135a-5p (98)<br>hsa-miR-548au-3p (96)<br>hsa-miR-651-3p (96)<br>hsa-miR-3064-3p (95) | uggcuuuuaacuuugauggc<br>ugugugugagaggaagaaggga<br>uuuugcaauauguuccugaaua<br>uauggcuuuucauuccuauguga<br>uauggcuuuuuauuccuauguga<br>uggcaguuacuuuugcaccag<br>aaaggaaaguguauccuaaaag<br>uugccacacugcaacaccuuaca |

| | | | | | | | | | | |
|---|---|---|---|---|---|---|---|---|---|---|
| | | | | | | like melanoma and osteosarcoma [9]. | (FGF21) mediated anti-fibrotic effects in kidney disease [10]. | | hsa-miR-7-5p (95)<br>hsa-miR-3680-3p (94) | uggaagacuagugauuuuguuguu<br>uuuugcaugacccugggaguagg |
| *Ndrg2* \|15\|- | NDRG Family Member 2 | | 39.2 (D) | 0.11 | 0.18 | NDRG2 is a tumor suppressor playing a key role in inhibiting tumor cell metastasis and proliferation. It is also involved in cellular stress responses [12]. | NDRG2 knockdown promotes renal fibrosis by increasing epithelial-mesenchymal transition markers and extracellular matrix components, regulated through the Smad3 pathway in renal tubular disease [13]. | --- | hsa-miR-5701 (92)<br>hsa-miR-139-5p (91)<br>hsa-miR-6755-5p (91)<br>hsa-miR-3059-5p (86)<br>hsa-miR-3974 (84)<br>hsa-miR-6801-5p (84)<br>hsa-miR-10393-3p (84)<br>hsa-miR-298 (84)<br>hsa-miR-4301 (80)<br>hsa-miR-4723-5p (78) | uuauugucacguucugauu<br>ucuacagugcacgugucuccagu<br>uaggguagacacugacaacguu<br>uuuccucucugccccauagggugu<br>aaaggucauuguaagguuaaugc<br>uggucagaggcagcaggaaauga<br>uggucagauuugaacucuucaa<br>agcagaagcagggagguucuccca<br>ucccacuacuucacuuguga<br>ugggggagccaugagauaagagca |
| *Slc25a24* \|2\|+) | Solute Carrier Family 25 Member 24 | | 39.1 (D) | 0.47 | 0.56 | *SLC25A24* encodes a carrier protein that transports ATP-Mg in exchange for phosphate and manages adenine nucleotide flow across the mitochondrial inner membrane. It protects cells from oxidative stress-induced death and buffers calcium in the mitochondrial matrix [14]. | Medial calcification in chronic kidney disease is linked to phosphate and calcium overload in vascular smooth muscle cells. Phosphate is delivered by mitochondrial transporters like SLC25A24. | --- | hsa-miR-3658 (98)<br>hsa-miR-5011-5p (98)<br>hsa-miR-607 (97)<br>hsa-miR-153-5p (97)<br>hsa-miR-3671 (94)<br>hsa-miR-6124 (94)<br>hsa-miR-10527-5p (92)<br>hsa-miR-659-3p (92)<br>hsa-miR-652-5p (91)<br>hsa-miR-5583-5p (91) | uuuaagaaaacaccauggagau<br>uauauauacagccaugcacuc<br>guucaaauccagaucuauaac<br>ucauuuuugugauguugcagcu<br>aucaaauaaggacuagucugca<br>gggaaaaggaagggggagga<br>aaagcaaauguugggugaacggc<br>cuugguucagggagggucccca<br>caacccuaggagagggugccauuca<br>aaacuaauauacccauauucug |
| *Nup50* \|7\|+ | Nucleoporin 50 | | 39.0 (D) | 0.15 | 0.02 | NUP50 is a component of the nuclear pore complex and an FG-repeat nucleoporin. It functions as a nuclear basket nucleoporin, involved in mRNA export and links the nuclear pore complex to transcription machinery [15]. | Differentially expressed in the glomeruli of patients with diabetic nephropathy [16]. | b [17] | hsa-miR-15b-5p (99)<br>hsa-miR-497-5p (99)<br>hsa-miR-16-5p (99)<br>hsa-miR-6838-5p (99)<br>hsa-miR-15a-5p (99)<br>hsa-miR-195-5p (99)<br>hsa-miR-424-5p (99)<br>hsa-miR-23a-3p (98)<br>hsa-miR-141-5p (98)<br>hsa-miR-23b-3p (98) | uagcagcacaucaugguuuaca<br>cagcagcacacugugguuugu<br>uagcagcacguaaauauuggcg<br>aagcagcaguggcaagacuccu<br>uagcagcacauaaugguuugug<br>uagcagcacagaaauauuggc<br>cagcagcaauucauguuuugaa<br>aucacauugccagggauuucc<br>caucuuccaguacaguguugga<br>aucacauugccagggauuaccac |
| *Fgfr1op* 3\|1\|- | Fibroblast Growth Factor Receptor 1 Oncogene Partner | | 38.8 (D) | 0.55 | 0.67 | FGFR1OP, also known as Centrosomal Protein 43 or FOP, is a hydrophilic centrosomal protein required for anchoring microtubules to subcellular structures. Fusion of CEP43 with FGFR1 has been found in cases of myeloproliferative disorder [18]. | Embryonic mouse kidneys homozygous for gene-trapped *FOP* were found to have absent cilia in the renal tubules [19]. | c [20] | hsa-miR-95-5p (99)<br>hsa-miR-9851-3p (96)<br>hsa-miR-593-5p (96)<br>hsa-miR-3662 (95)<br>hsa-miR-203a-3p (95)<br>hsa-miR-1231 (94)<br>hsa-miR-4437 (94)<br>hsa-miR-4708-3p (93)<br>hsa-miR-4738-3p (90)<br>hsa-miR-8057 (89) | ucaauaaaugucuguugaauu<br>uggcaccagcacuggcgguguc<br>aggcaccagccaggcauugcucagc<br>gaaaaugaugaguagugacugaug<br>gugaaauguuuaggaccacuag<br>gugucugggcggacagcugc<br>ugggcucaggguacaaagguu<br>agcaaggcggcaucucucugau<br>ugaaacuggagcgccuggagga<br>guggcucuguaguaagaugga |
| *Mgat3* \|7\|+ | Beta-1,4-mannosyl-glycoprotein 4-beta-N-acetylglucos aminyltransf erase | | 38.4 (D) | -0.6 | -0.84 | This enzyme synthesizes triacylglycerol, highly expressed in the small intestine, aiding chylomicron formation. MGAT3 expression increases lipid droplet size and number [21]. | In rat renal macrophages, Mgat3 was differentially expressed in response to a 5% Hydroxy-L-proline diet, indicating that oxalate may affect macrophage phagocytic activity and response to kidney crystals [22]. | --- | hsa-miR-4755-5p (98)<br>hsa-miR-211-5p (98)<br>hsa-miR-204-5p (98)<br>hsa-miR-5006-3p (98)<br>hsa-miR-324-5p (95)<br>hsa-miR-4795-3p (93)<br>hsa-miR-4286 (93)<br>hsa-miR-126-5p (93)<br>hsa-miR-8084 (91)<br>hsa-miR-1295b-5p (91) | uuucccuucagagccuggcuuu<br>uucccuuugucauccuucgccu<br>uucccuuugucauccuaugccu<br>uuucccuuuccauccuggcag<br>cgcauccccuagggcauuggug<br>auauuauuagccacuucuggau<br>accccacuccugguacc<br>cauuauuacuuuugguacgcg<br>gaauacuaaguaaaaaaucagua<br>cacccagaucugcggccuaau |

a; APA of *FOXN3* has been reported that leads to shortened mRNA lacking the tumor suppressive function of the full-length protein [11].

b; APA of *NUP50* at the 3'UTR has been reported to lead to three alternate transcripts that are 2kb and additional 0.8 kb and 3.2 kb longer transcripts [17].
c; APA of *FGFR1OP* has been reported at its 3'UTR in proliferating versus inactive fibroblasts [20].
#; For each APA gene, predicted miRNA sites were identified from miRNA database (miRDB.org). A target score of 74 or above was selected (listed In parenthesis), and at maximum ten miRs with the highest target scores are listed for each gene. For each miR, its sequence is listed as well.
(P) denotes proximal shift and (D) denotes distal shift in the usage of poly(A) site

Supplementary Table 10: APA genes reversed with pioglitazone treatment

| **Gene** | **% APA Difference** | ***P* adj.** | **r#** |
|---|---|---|---|
| *Ube2s\|1\|+* | 32.60 | 0.000 | -0.367 |
| *Fndc3b\|2\|-* | 29.64 | 0.000 | -0.279 |
| *Lyc2\|7\|+* | 29.30 | 0.000 | 0.344 |
| *Mmd\|10\|+* | 25.58 | 0.014 | -0.368 |
| *Cd47\|11\|-* | 25.09 | 0.011 | -0.285 |
| *Lrch3\|11\|-* | 23.22 | 0.001 | -0.318 |
| *Slc30a4\|3\|-* | 22.56 | 0.003 | -0.300 |
| *rnf141\|1\|-* | 22.52 | 0.001 | -0.312 |
| *Sgpl1\|20\|-* | 22.41 | 0.019 | -0.291 |
| *Pi4k2a\|1\|+* | 22.37 | 0.022 | -0.332 |
| *Arrdc3\|2\|+* | 21.90 | 0.036 | -0.266 |
| *Serpine1\|12\|+* | 21.36 | 0.013 | -0.264 |
| *Retreg1\|2\|+* | 21.33 | 0.000 | -0.282 |
| *Mcc\|18\|-* | 17.25 | 0.002 | 0.240 |
| *Armcx1\|X\|+* | 16.94 | 0.028 | -0.191 |
| *Fam120a\|17\|+* | 16.61 | 0.001 | -0.286 |
| *Ncoa4\|16\|+* | 16.15 | 0.047 | -0.186 |
| *Golim4\|2\|-* | 15.41 | 0.000 | -0.076 |
| *Vps52\|20\|-* | 14.65 | 0.045 | 0.165 |
| *Atp11a\|16\|-* | 14.26 | 0.045 | -0.196 |
| *Lpl\|16\|-* | 14.14 | 0.046 | -0.156 |
| *Ugcg\|5\|+* | 13.30 | 0.009 | -0.199 |
| *Lgals3bp\|10\|-* | 13.09 | 0.046 | -0.187 |
| *Bcl2l2\|15\|+* | 12.76 | 0.011 | -0.168 |
| *Slc6a6\|4\|-* | 12.75 | 0.000 | -0.234 |
| *Fam117b\|9\|+* | 12.49 | 0.000 | -0.186 |
| *Ube2h\|4\|-* | 11.88 | 0.009 | -0.220 |
| *Sptbn1\|14\|-* | 11.04 | 0.000 | -0.147 |
| *Aldh9a1\|13\|+* | 10.81 | 0.011 | -0.092 |
| *St3gal1\|7\|-* | 10.78 | 0.000 | -0.238 |
| *Cox5a\|8\|+* | 10.74 | 0.000 | -0.155 |
| *Dusp3\|10\|-* | 10.48 | 0.006 | -0.142 |
| *Des\|9\|+* | 10.46 | 0.000 | 0.096 |
| *Calm3\|1\|-* | 10.10 | 0.042 | -0.106 |
| *Kirrel1\|2\|-* | 9.70 | 0.002 | -0.163 |
| *Rbm39\|3\|-* | 9.44 | 0.036 | 0.103 |
| *Tns3\|14\|-* | 8.50 | 0.000 | -0.167 |
| *Akap2\|5\|+* | 8.48 | 0.047 | -0.142 |
| *Scpep1\|10\|-* | 8.35 | 0.021 | 0.083 |
| *Rn45s\|14\|+* | 7.66 | 0.000 | 0.089 |
| *Msn\|X\|+* | 7.49 | 0.000 | -0.109 |
| *Adamts1\|11\|-* | 7.49 | 0.026 | -0.089 |
| *Clic4\|5\|-* | 6.98 | 0.042 | -0.115 |
| *Npr3\|2\|-* | 6.98 | 0.013 | -0.171 |

| | | | |
|---|---|---|---|
| *Aif1l\|3\|-* | 6.80 | 0.007 | -0.109 |
| *Tgfbr2\|8\|-* | 6.66 | 0.044 | -0.035 |
| *Prkar1a\|10\|+* | 6.58 | 0.044 | -0.069 |
| *Igfbp5\|9\|-* | 6.54 | 0.000 | -0.093 |
| *Synpo\|18\|-* | 6.47 | 0.016 | -0.084 |
| *Actg1\|3\|+* | 6.20 | 0.003 | -0.067 |
| *Itgb1\|19\|+* | 5.64 | 0.013 | -0.065 |
| *Calr\|19\|-* | 4.73 | 0.001 | -0.049 |
| *Sparc\|10\|-* | 4.16 | 0.000 | -0.035 |
| *Podxl\|4\|-* | 3.94 | 0.029 | 0.048 |
| *Vasn\|10\|-* | 2.71 | 0.005 | -0.125 |
| *Actb\|12\|+* | 2.52 | 0.015 | -0.025 |

[#] *'+ r value denotes distal shift and - r value denotes proximal shift*

Supplementary Table 11: APA genes reversed with GQ-16 treatment

| **Gene** | **% APA Difference** | ***P* adj.** | **r#** |
|---|---|---|---|
| *Nrbp2\|7\|-* | 30.06 | 0.006 | 0.337 |
| *Lyc2\|7\|+* | 19.20 | 0.000 | 0.218 |
| *Hhip\|19\|+* | 17.49 | 0.013 | -0.215 |
| *Lima1\|7\|-* | 16.78 | 0.001 | 0.167 |
| *Rpl12\|3\|+* | 16.09 | 0.001 | 0.162 |
| *Kcnj16\|10\|+* | 15.26 | 0.018 | -0.105 |
| *Srsf4\|5\|+* | 14.51 | 0.039 | 0.152 |
| *Aif1l\|3\|-* | 12.78 | 0.000 | -0.181 |
| *Fam81a\|8\|-* | 11.08 | 0.023 | -0.110 |
| *Kirrel1\|2\|-* | 11.05 | 0.000 | -0.180 |
| *Igfbp5\|9\|-* | 9.72 | 0.000 | 0.103 |
| *Tgfbr2\|8\|-* | 7.45 | 0.003 | 0.057 |
| *Epas1\|6\|+* | 5.50 | 0.013 | 0.070 |
| *Sema3g\|16\|+* | 5.32 | 0.000 | -0.059 |
| *Tuba1b\|7\|-* | 5.06 | 0.002 | -0.048 |
| *Calr\|19\|-* | 4.89 | 0.018 | -0.049 |
| *Tns3\|14\|-* | 4.57 | 0.003 | -0.130 |
| *Tmsb4x\|X\|+* | 4.28 | 0.000 | 0.044 |
| *Gpx3\|10\|+* | 3.12 | 0.000 | 0.031 |
| *Eef1a1\|8\|-* | 1.33 | 0.039 | -0.032 |

### *'+ r value denotes distal shift and - r value denotes proximal shift*

Supplementary Table 12: Dysregulated splicing factors in PAN and ADR models

| | | PAN | |
|---|---|---|---|
| **Gene** | **Class/family** | **logFC PANvsCtl** | **adj.P.Val PANvsCtl** |
| *Phf5a* | 17S U2 snRNP | 0.38 | 0.03 |
| *Sf3a1* | 17S U2 snRNP | 0.27 | 0.05 |
| *Sf3a2* | 17S U2 snRNP | 0.39 | 0.02 |
| *Sf3a3* | 17S U2 snRNP | 0.39 | 0.03 |
| *Sf3b2* | 17S U2 snRNP | 0.30 | 0.05 |
| *Sf3b3* | 17S U2 snRNP | 0.40 | 0.01 |
| *Sf3b4* | 17S U2 snRNP | 0.41 | 0.02 |
| *Sf3b5* | 17S U2 snRNP | 0.53 | 0.03 |
| *Sf3b6* | 17S U2 snRNP | 0.38 | 0.04 |
| *Snrpa1* | 17S U2 snRNP | 0.47 | 0.03 |
| *Snrpb2* | 17S U2 snRNP | 0.49 | 0.01 |
| *Cherp* | 17S U2 snRNPassociated | 0.37 | 0.04 |
| *Rbm17* | 17S U2 snRNPassociated | 0.40 | 0.01 |
| *Smndc1* | 17S U2 snRNPassociated | 0.29 | 0.05 |
| *U2af1* | 17S U2 snRNPassociated | 0.39 | 0.04 |
| *Elavl1* | Alternative Splicing Factors | 0.31 | 0.03 |
| *Mbnl2* | Alternative Splicing Factors | 0.43 | 0.03 |
| *Mbnl3* | Alternative Splicing Factors | 1.39 | 0.01 |
| *Khdrbs3* | Alternative Splicing Factors | 1.25 | 0.01 |
| *Celf2* | Alternative Splicing Factors | 0.73 | 0.04 |
| *Ddx50* | Detected in Bact complex | 0.42 | 0.03 |
| *Mov10* | Detected in Bact complex | 0.38 | 0.04 |
| *Trim24* | Detected in Bact complex | 0.61 | 0.02 |
| *Toe1* | Detected in C complex | 0.52 | 0.01 |
| *Alyref* | EJC/mRNP | 0.40 | 0.02 |
| *Eif4a3* | EJC/mRNP | 0.30 | 0.04 |
| *Magoh* | EJC/mRNP | 0.43 | 0.02 |
| *Nxt1* | EJC/mRNP | 0.89 | 0.00 |
| *Rbm8a* | EJC/mRNP | 0.40 | 0.02 |
| *Rnps1* | EJC/mRNP | 0.42 | 0.01 |
| *Sap18* | EJC/mRNP | 0.49 | 0.01 |
| *Hnrnpa1* | hnRNP | 0.46 | 0.01 |
| *Hnrnpa3* | hnRNP | 0.31 | 0.04 |
| *Hnrnph2* | hnRNP | 0.29 | 0.04 |
| *Hnrnpk* | hnRNP | 0.34 | 0.03 |
| *Hnrnpl* | hnRNP | 0.41 | 0.02 |
| *Hnrnpr* | hnRNP | 0.35 | 0.04 |
| *Raly* | hnRNP | 0.39 | 0.02 |
| *Rbmx* | hnRNP | 0.37 | 0.05 |
| *Syncrip* | hnRNP | 0.32 | 0.05 |
| *Nono* | linked to splicing | 0.35 | 0.02 |
| *Sfpq* | linked to splicing | 0.40 | 0.02 |
| *Srpk1* | linked to splicing | 0.26 | 0.05 |
| *Lsm1* | LSm | 0.46 | 0.03 |
| *Lsm2* | LSm | 0.68 | 0.01 |
| *Lsm3* | LSm | 0.49 | 0.01 |
| *Lsm4* | LSm | 0.54 | 0.01 |
| *Lsm5* | LSm | 0.57 | 0.02 |
| *Lsm6* | LSm | 0.49 | 0.02 |
| *Lsm7* | LSm | 0.46 | 0.03 |
| *Naa38* | LSm | 0.60 | 0.01 |
| *Bcas2* | PRP19 complex | 0.36 | 0.05 |
| *Ctnnbl1* | PRP19 complex | 0.30 | 0.04 |
| *Plrg1* | PRP19 complex | 0.26 | 0.04 |
| *Pqbp1* | PRP19 complex | 0.47 | 0.02 |
| *Prpf19* | PRP19 complex | 0.68 | 0.01 |
| *Wbp11* | PRP19 complex | 0.37 | 0.02 |
| *Bud31* | PRP19-related | 0.94 | 0.00 |
| *Isy1* | PRP19-related | 0.50 | 0.01 |
| *Ppie* | PRP19-related | 0.54 | 0.03 |
| *Ppil1* | PRP19-related | 0.59 | 0.00 |
| *Rbm22* | PRP19-related | 0.40 | 0.02 |
| *Champ1* | RBD-containing | 0.26 | 0.04 |
| *Ddx18* | RBD-containing | 0.32 | 0.04 |
| *Ddx27* | RBD-containing | 0.30 | 0.04 |
| *Ddx39b* | RBD-containing | 0.48 | 0.04 |
| *Rbm3* | RBD-containing | 0.69 | 0.02 |
| *Rbmxl1* | RBD-containing | 0.46 | 0.01 |
| *Zc3h18* | RBD-containing | 0.32 | 0.04 |
| *Zcrb1* | RBD-containing | 0.55 | 0.02 |
| *Ewsr1* | RBP | 0.44 | 0.03 |
| *Ncbp2* | RBP | 0.38 | 0.05 |
| *Pabpc1* | RBP | 0.70 | 0.01 |
| *Pabpc4* | RBP | 0.49 | 0.04 |
| *Prpf38a* | Recruited at B complex | 0.34 | 0.02 |
| *Zmat2* | Recruited at B complex | 0.39 | 0.02 |
| *Cwc25* | Recruited at Bact complex | 0.34 | 0.05 |
| *Gpatch1* | Recruited at Bact complex | 0.44 | 0.02 |
| *Ppil2* | Recruited at Bact complex | 0.44 | 0.02 |
| *Yju2* | Recruited at Bact complex | 0.47 | 0.01 |
| *Ddx41* | Recruited at C complex | 0.36 | 0.04 |
| *Dhx35* | Recruited at C complex | 0.46 | 0.02 |
| *Nosip* | Recruited at C complex | 0.39 | 0.04 |
| *Ppil3* | Recruited at C complex | 0.65 | 0.00 |
| *Ppwd1* | Recruited at C complex | 0.41 | 0.03 |
| *Dhx38* | Second step factors | 0.37 | 0.02 |
| *Dhx8* | Second step factors | 0.30 | 0.04 |
| *Snrpb* | Sm | 0.31 | 0.04 |
| *Snrpd1* | Sm | 0.65 | 0.00 |
| *Snrpd2* | Sm | 0.46 | 0.01 |
| *Snrpd3* | Sm | 0.44 | 0.01 |
| *Snrpf* | Sm | 0.43 | 0.02 |
| *Srsf2* | SR protein | 0.42 | 0.05 |
| *Srsf3* | SR protein | 0.40 | 0.02 |
| *Srsf4* | SR protein | 0.41 | 0.02 |
| *Srsf6* | SR protein | 0.40 | 0.05 |
| *Srsf7* | SR protein | 0.36 | 0.04 |
| *Srsf9* | SR protein | 0.52 | 0.01 |
| *Tra2b* | SR protein | 0.37 | 0.02 |
| *Thoc3* | TREX | 0.38 | 0.03 |
| *Thoc5* | TREX | 0.46 | 0.02 |
| *Thoc6* | TREX | 0.72 | 0.01 |
| *Thoc7* | TREX | 0.65 | 0.02 |
| *Usp39* | tri-snRNP | 0.36 | 0.02 |
| *Snrpa* | U1 snRNP | 0.66 | 0.00 |
| *Snrpc* | U1 snRNP | 0.50 | 0.00 |

| | | ADR | |
|---|---|---|---|
| **Gene** | **Class/family** | **logFC ADRvsCtl** | **adj.P.Val ADRvsCtl** |
| *Phf5a* | 17S U2 snRNP | 0.4887 | 0.0063 |
| *Sf3a3* | 17S U2 snRNP | 0.4134 | 0.0147 |
| *Sf3b1* | 17S U2 snRNP | 0.5080 | 0.0077 |
| *Sf3b2* | 17S U2 snRNP | 0.4549 | 0.0043 |
| *Sf3b3* | 17S U2 snRNP | 0.3942 | 0.0085 |
| *Sf3b5* | 17S U2 snRNP | 0.6746 | 0.0051 |
| *Sf3b6* | 17S U2 snRNP | 0.4777 | 0.0093 |
| *Snrpa1* | 17S U2 snRNP | 0.5067 | 0.0133 |
| *Snrpb2* | 17S U2 snRNP | 0.6575 | 0.0011 |
| *Dhx15* | 17S U2 snRNPassociated | 0.3310 | 0.0133 |
| *Htatsf1* | 17S U2 snRNPassociated | 0.5496 | 0.0117 |
| *Puf60* | 17S U2 snRNPassociated | 0.3118 | 0.0331 |
| *Rbm17* | 17S U2 snRNPassociated | 0.4972 | 0.0021 |
| *Smndc1* | 17S U2 snRNPassociated | 0.3926 | 0.0081 |
| *U2af1* | 17S U2 snRNPassociated | 0.5091 | 0.0070 |
| *Khdrbs3* | Alternative Splicing Factors | 1.6470 | 0.0008 |
| *Elavl1* | Alternative Splicing Factors | 0.3193 | 0.0194 |
| *Mbnl2* | Alternative Splicing Factors | 0.5283 | 0.0075 |
| *Mbnl3* | Alternative Splicing Factors | 1.1390 | 0.0116 |
| *Ptbp2* | Alternative Splicing Factors | 0.5175 | 0.0393 |
| *Ddx50* | Detected in Bact complex | 0.5738 | 0.0035 |
| *Frg1* | Detected in Bact complex | 0.6508 | 0.0016 |
| *Trim24* | Detected in Bact complex | 0.6519 | 0.0101 |
| *Matr3* | Detected in C complex | 0.4123 | 0.0135 |
| *Rbm4b* | Detected in C complex | 0.3293 | 0.0370 |
| *Toe1* | Detected in C complex | 0.5609 | 0.0035 |
| *Alyref* | EJC/mRNP | 0.4226 | 0.0124 |
| *Eif4a3* | EJC/mRNP | 0.3632 | 0.0116 |
| *Magoh* | EJC/mRNP | 0.5856 | 0.0029 |
| *Nxt1* | EJC/mRNP | 0.8774 | 0.0005 |
| *Rbm8a* | EJC/mRNP | 0.7169 | 0.0004 |
| *Rnps1* | EJC/mRNP | 0.4489 | 0.0059 |
| *Sap18* | EJC/mRNP | 0.7400 | 0.0007 |
| *Prpf38b* | Found with spliceosomes | 0.3613 | 0.0340 |
| *Prpf39* | Found with spliceosomes | 0.5487 | 0.0070 |
| *Snrnp27* | Found with spliceosomes | 0.4359 | 0.0063 |
| *Hnrnpa1* | hnRNP | 0.3357 | 0.0378 |
| *Hnrnpa3* | hnRNP | 0.4734 | 0.0039 |
| *Hnrnpab* | hnRNP | 0.3321 | 0.0367 |
| *Hnrnpc* | hnRNP | 0.6416 | 0.0010 |
| *Hnrnph2* | hnRNP | 0.4138 | 0.0044 |
| *Hnrnph3* | hnRNP | 0.3659 | 0.0417 |
| *Hnrnpk* | hnRNP | 0.3773 | 0.0104 |
| *Hnrnpl* | hnRNP | 0.4215 | 0.0102 |
| *Hnrnpm* | hnRNP | 0.3217 | 0.0169 |
| *Raly* | hnRNP | 0.4054 | 0.0130 |
| *Syncrip* | hnRNP | 0.3161 | 0.0406 |
| *Dbr1* | linked to splicing | 0.5116 | 0.0051 |
| *Nono* | linked to splicing | 0.3700 | 0.0112 |
| *Sfpq* | linked to splicing | 0.3650 | 0.0165 |
| *Lsm1* | LSm | 0.5716 | 0.0072 |
| *Lsm2* | LSm | 0.5770 | 0.0104 |
| *Lsm3* | LSm | 0.6154 | 0.0027 |
| *Lsm4* | LSm | 0.6655 | 0.0023 |
| *Lsm5* | LSm | 0.8138 | 0.0021 |
| *Lsm6* | LSm | 0.4858 | 0.0121 |
| *Lsm7* | LSm | 0.6404 | 0.0037 |
| *Naa38* | LSm | 0.6837 | 0.0016 |
| *Bcas2* | PRP19 complex | 0.6289 | 0.0023 |
| *Cdc5l* | PRP19 complex | 0.2641 | 0.0407 |
| *Ctnnbl1* | PRP19 complex | 0.3637 | 0.0136 |
| *Cwc15* | PRP19 complex | 0.5091 | 0.0074 |
| *Plrg1* | PRP19 complex | 0.4025 | 0.0037 |
| *Pqbp1* | PRP19 complex | 0.6174 | 0.0031 |
| *Prpf19* | PRP19 complex | 0.5438 | 0.0116 |
| *Aqr* | PRP19-related | 0.4110 | 0.0100 |
| *Bud31* | PRP19-related | 0.8411 | 0.0035 |
| *Crnkl1* | PRP19-related | 0.3737 | 0.0433 |
| *Isy1* | PRP19-related | 0.5165 | 0.0078 |
| *Ppie* | PRP19-related | 0.7742 | 0.0026 |
| *Ppil1* | PRP19-related | 0.5652 | 0.0022 |
| *Snw1* | PRP19-related | 0.4320 | 0.0140 |
| *Xab2* | PRP19-related | 0.4070 | 0.0118 |
| *Champ1* | RBD-containing | 0.3556 | 0.0075 |
| *Ddx1* | RBD-containing | 0.4751 | 0.0060 |
| *Ddx18* | RBD-containing | 0.4177 | 0.0078 |
| *Ddx27* | RBD-containing | 0.3697 | 0.0138 |
| *Rbm3* | RBD-containing | 0.6840 | 0.0109 |
| *Rbm42* | RBD-containing | 0.3943 | 0.0108 |
| *Rbmxl1* | RBD-containing | 0.6350 | 0.0012 |
| *Rnf20* | RBD-containing | 0.4453 | 0.0097 |
| *Rnf34* | RBD-containing | 0.2965 | 0.0418 |
| *Zc3h18* | RBD-containing | 0.3179 | 0.0274 |
| *Zc3hav1* | RBD-containing | 0.4241 | 0.0320 |
| *Zcrb1* | RBD-containing | 0.7332 | 0.0024 |
| *Zfr* | RBD-containing | 0.5644 | 0.0057 |
| *Ddx3x* | RBP | 0.2997 | 0.0233 |
| *Ewsr1* | RBP | 0.3892 | 0.0392 |
| *Ncbp2* | RBP | 0.5525 | 0.0055 |
| *Pabpc1* | RBP | 0.6314 | 0.0082 |
| *Rbm39* | RBP | 0.5664 | 0.0057 |
| *Rbm7* | RBP | 0.3734 | 0.0043 |
| *Ybx1* | RBP | 0.3054 | 0.0464 |
| *Ccar1* | Recruited at A complex | 0.3757 | 0.0491 |
| *Prpf40a* | Recruited at A complex | 0.3128 | 0.0393 |
| *Rbm5* | Recruited at A complex | 0.4306 | 0.0311 |
| *Prpf38a* | Recruited at B complex | 0.5069 | 0.0019 |
| *Smu1* | Recruited at B complex | 0.2344 | 0.0479 |
| *Wbp4* | Recruited at B complex | 0.3627 | 0.0302 |
| *Zmat2* | Recruited at B complex | 0.5729 | 0.0014 |
| *Cwc22* | Recruited at Bact complex | 0.5710 | 0.0066 |
| *Gpatch1* | Recruited at Bact complex | 0.4999 | 0.0070 |
| *Ppil2* | Recruited at Bact complex | 0.4916 | 0.0093 |
| *Yju2* | Recruited at Bact complex | 0.4577 | 0.0108 |
| *Ddx41* | Recruited at C complex | 0.4226 | 0.0145 |
| *Dhx35* | Recruited at C complex | 0.4078 | 0.0213 |

| | | | |
|---|---|---|---|
| *Sart3* | U4/U6 recycling | 0.35 | 0.03 |
| *Prpf31* | U4/U6 snRNP | 0.56 | 0.00 |
| *Cd2bp2* | U5 snRNP | 0.61 | 0.01 |
| *Ddx23* | U5 snRNP | 0.37 | 0.03 |
| *Eftud2* | U5 snRNP | 0.28 | 0.03 |
| *Snrnp40* | U5 snRNP | 0.41 | 0.02 |

| | | | |
|---|---|---|---|
| *Fam32a* | Recruited at C complex | 0.5474 | 0.0021 |
| *Fam50a* | Recruited at C complex | 0.3229 | 0.0414 |
| *Fra10ac1* | Recruited at C complex | 0.4818 | 0.0477 |
| *Nosip* | Recruited at C complex | 0.5316 | 0.0078 |
| *Ppig* | Recruited at C complex | 0.4753 | 0.0270 |
| *Ppil3* | Recruited at C complex | 0.6899 | 0.0011 |
| *Ppwd1* | Recruited at C complex | 0.5157 | 0.0083 |
| *Syf2* | Recruited at C complex | 0.5084 | 0.0030 |
| *Wdr83* | Recruited at C complex | 0.5627 | 0.0182 |
| *Bud13* | RES complex | 0.4054 | 0.0252 |
| *Rbmx2* | RES complex | 0.7100 | 0.0095 |
| *Cdc40* | Second step factors | 0.2898 | 0.0147 |
| *Dhx38* | Second step factors | 0.3915 | 0.0103 |
| *Dhx8* | Second step factors | 0.3119 | 0.0268 |
| *Slu7* | Second step factors | 0.4952 | 0.0056 |
| *Snrpd1* | Sm | 0.6074 | 0.0020 |
| *Snrpd2* | Sm | 0.6297 | 0.0011 |
| *Snrpd3* | Sm | 0.4383 | 0.0045 |
| *Snrpe* | Sm | 0.6350 | 0.0326 |
| *Snrpf* | Sm | 0.5736 | 0.0035 |
| *Srsf1* | SR protein | 0.3335 | 0.0304 |
| *Srsf10* | SR protein | 0.4026 | 0.0338 |
| *Srsf2* | SR protein | 0.4020 | 0.0417 |
| *Srsf3* | SR protein | 0.6431 | 0.0007 |
| *Srsf4* | SR protein | 0.3837 | 0.0235 |
| *Srsf6* | SR protein | 0.3836 | 0.0420 |
| *Srsf7* | SR protein | 0.6338 | 0.0015 |
| *Srsf9* | SR protein | 0.6294 | 0.0014 |
| *Tra2a* | SR protein | 0.5940 | 0.0151 |
| *Tra2b* | SR protein | 0.6088 | 0.0006 |
| *Thoc1* | TREX | 0.6602 | 0.0024 |
| *Thoc3* | TREX | 0.4102 | 0.0142 |
| *Thoc5* | TREX | 0.3704 | 0.0309 |
| *Thoc6* | TREX | 0.6065 | 0.0218 |
| *Thoc7* | TREX | 1.0824 | 0.0007 |
| *Usp39* | tri-snRNP | 0.4715 | 0.0026 |
| *Luc7l* | U1 snRNP | 0.5149 | 0.0112 |
| *Snrpa* | U1 snRNP | 0.3866 | 0.0480 |
| *Snrpc* | U1 snRNP | 0.3820 | 0.0134 |
| *Snrnp25* | U11/U12 snRNP | 0.6147 | 0.0043 |
| *Snrnp35* | U11/U12 snRNP | 0.4065 | 0.0467 |
| *Prpf3* | U4/U6 snRNP | 0.4743 | 0.0144 |
| *Prpf31* | U4/U6 snRNP | 0.6366 | 0.0007 |
| *Cd2bp2* | U5 snRNP | 0.6016 | 0.0073 |
| *Ddx23* | U5 snRNP | 0.4901 | 0.0063 |
| *Prpf6* | U5 snRNP | 0.4073 | 0.0096 |
| *Snrnp200* | U5 snRNP | 0.3593 | 0.0241 |
| *Snrnp40* | U5 snRNP | 0.4276 | 0.0082 |
| *Txnl4a* | U5 snRNP | 0.4653 | 0.0202 |

Supplementary Table 13: Dysregulated polyadenylation factors in PAN and ADR models

**PAN**

| Gene | APA Factors | logFC PANvsCtl | adj.P.Val PANvsCtl |
|---|---|---|---|
| *Cpsf1* | Core Sequence Elements and Factors | 0.367 | 0.03 |
| *Cstf1* | Core Sequence Elements and Factors | 0.307 | 0.03 |
| *Cstf2* | Core Sequence Elements and Factors | 0.471 | 0.01 |
| *Cstf3* | Core Sequence Elements and Factors | 0.38 | 0.04 |
| *Cpsf1* | Proteins of Human Pre-mRNA Cleavage Complex | 0.37 | 0.03 |
| *Cstf2* | Proteins of Human Pre-mRNA Cleavage Complex | 0.47 | 0.01 |
| *Cstf3* | Proteins of Human Pre-mRNA Cleavage Complex | 0.38 | 0.04 |
| *Ddx49* | Proteins of Human Pre-mRNA Cleavage Complex | 0.54 | 0.00 |
| *Dhx8* | Proteins of Human Pre-mRNA Cleavage Complex | 0.30 | 0.04 |
| *Farsb* | Proteins of Human Pre-mRNA Cleavage Complex | 0.40 | 0.04 |
| *Hnrnpa3* | Proteins of Human Pre-mRNA Cleavage Complex | 0.31 | 0.04 |
| *Lmna* | Proteins of Human Pre-mRNA Cleavage Complex | 0.66 | 0.01 |
| *Mdc1* | Proteins of Human Pre-mRNA Cleavage Complex | 0.67 | 0.01 |
| *Nelfb* | Proteins of Human Pre-mRNA Cleavage Complex | 0.41 | 0.02 |
| *Nelfe* | Proteins of Human Pre-mRNA Cleavage Complex | 0.57 | 0.01 |
| *Nono* | Proteins of Human Pre-mRNA Cleavage Complex | 0.35 | 0.02 |
| *Nudt21* | Proteins of Human Pre-mRNA Cleavage Complex | 0.40 | 0.01 |
| *Pabpc1* | Proteins of Human Pre-mRNA Cleavage Complex | 0.70 | 0.01 |
| *Parp1* | Proteins of Human Pre-mRNA Cleavage Complex | 0.41 | 0.02 |
| *Rpl24* | Proteins of Human Pre-mRNA Cleavage Complex | 0.48 | 0.01 |
| *Rpl27* | Proteins of Human Pre-mRNA Cleavage Complex | 0.31 | 0.05 |
| *Rpl8* | Proteins of Human Pre-mRNA Cleavage Complex | 0.42 | 0.02 |
| *Rps24* | Proteins of Human Pre-mRNA Cleavage Complex | 0.44 | 0.03 |
| *Rps5* | Proteins of Human Pre-mRNA Cleavage Complex | 0.34 | 0.04 |
| *Scaf1* | Proteins of Human Pre-mRNA Cleavage Complex | 0.35 | 0.05 |
| *Smarca4* | Proteins of Human Pre-mRNA Cleavage Complex | 0.47 | 0.03 |
| *Smarcd2* | Proteins of Human Pre-mRNA Cleavage Complex | 0.32 | 0.03 |
| *Thoc6* | Proteins of Human Pre-mRNA Cleavage Complex | 0.72 | 0.01 |
| *Top2a* | Proteins of Human Pre-mRNA Cleavage Complex | 2.63 | 0.00 |
| *Usp39* | Proteins of Human Pre-mRNA Cleavage Complex | 0.36 | 0.02 |
| *Xrcc6* | Proteins of Human Pre-mRNA Cleavage Complex | 0.74 | 0.00 |
| *Cpsf1* | Known PolyA Factors of CPSF Complex | 0.37 | 0.03 |
| *Cstf1* | Known PolyA Factors of CPSF Complex | 0.31 | 0.03 |
| *Cstf2* | Known PolyA Factors of CPSF Complex | 0.47 | 0.01 |
| *Cstf3* | Known PolyA Factors of CPSF Complex | 0.38 | 0.04 |
| *Pabpc1* | Other Known PolyA Factors | 0.70 | 0.01 |
| *Pabpc4* | Other Known PolyA Factors | 0.49 | 0.04 |
| *Ddx23* | Factors with Known Motifs | 0.37 | 0.03 |
| *Snd1* | Factors with Known Motifs | 0.68 | 0.01 |

**ADR**

| Gene | APA Factors | logFC ADRvsCtl | adj.P.Val ADRvsCtl |
|---|---|---|---|
| *Cstf2* | Core Sequence Elements and Factors | 0.3667 | 0.0333 |
| *Cstf3* | Core Sequence Elements and Factors | 0.4457 | 0.0164 |
| *Fip1l1* | Core Sequence Elements and Factors | 0.5514 | 0.0076 |
| *Cstf2* | Proteins of Human Pre-mRNA Cleavage Complex | 0.3667 | 0.0333 |
| *Cstf3* | Proteins of Human Pre-mRNA Cleavage Complex | 0.4457 | 0.0164 |
| *Nudt21* | Proteins of Human Pre-mRNA Cleavage Complex | 0.4128 | 0.0027 |
| *Fip1l1* | Proteins of Human Pre-mRNA Cleavage Complex | 0.5514 | 0.0076 |
| *Pabpc1* | Proteins of Human Pre-mRNA Cleavage Complex | 0.6314 | 0.0082 |
| *Ppp1cb* | Proteins of Human Pre-mRNA Cleavage Complex | 0.2828 | 0.0220 |
| *Top2a* | Proteins of Human Pre-mRNA Cleavage Complex | 1.9162 | 0.0056 |
| *Parp1* | Proteins of Human Pre-mRNA Cleavage Complex | 0.3186 | 0.0433 |
| *Prkdc* | Proteins of Human Pre-mRNA Cleavage Complex | 0.3892 | 0.0457 |
| *Xrcc6* | Proteins of Human Pre-mRNA Cleavage Complex | 0.6845 | 0.0021 |
| *Thoc6* | Proteins of Human Pre-mRNA Cleavage Complex | 0.6065 | 0.0218 |
| *Nelfb* | Proteins of Human Pre-mRNA Cleavage Complex | 0.4534 | 0.0054 |
| *Nelfe* | Proteins of Human Pre-mRNA Cleavage Complex | 0.6760 | 0.0030 |
| *Smarca4* | Proteins of Human Pre-mRNA Cleavage Complex | 0.4665 | 0.0212 |
| *Smarcd2* | Proteins of Human Pre-mRNA Cleavage Complex | 0.3123 | 0.0292 |
| *Erh* | Proteins of Human Pre-mRNA Cleavage Complex | 0.3311 | 0.0393 |
| *Rnf20* | Proteins of Human Pre-mRNA Cleavage Complex | 0.4453 | 0.0097 |
| *Usp39* | Proteins of Human Pre-mRNA Cleavage Complex | 0.4715 | 0.0026 |
| *Sf3b1* | Proteins of Human Pre-mRNA Cleavage Complex | 0.5080 | 0.0077 |
| *Nono* | Proteins of Human Pre-mRNA Cleavage Complex | 0.3700 | 0.0112 |
| *Srsf1* | Proteins of Human Pre-mRNA Cleavage Complex | 0.3335 | 0.0304 |
| *Srsf10* | Proteins of Human Pre-mRNA Cleavage Complex | 0.4026 | 0.0338 |
| *Hnrnpa3* | Proteins of Human Pre-mRNA Cleavage Complex | 0.4734 | 0.0039 |
| *Ddx49* | Proteins of Human Pre-mRNA Cleavage Complex | 0.6733 | 0.0008 |
| *Dhx8* | Proteins of Human Pre-mRNA Cleavage Complex | 0.3119 | 0.0268 |
| *Dhx15* | Proteins of Human Pre-mRNA Cleavage Complex | 0.3310 | 0.0133 |
| *Exosc4* | Proteins of Human Pre-mRNA Cleavage Complex | 0.3637 | 0.0229 |
| *Exosc7* | Proteins of Human Pre-mRNA Cleavage Complex | 0.3230 | 0.0375 |
| *Exosc8* | Proteins of Human Pre-mRNA Cleavage Complex | 0.5351 | 0.0201 |
| *Rps5* | Proteins of Human Pre-mRNA Cleavage Complex | 0.4909 | 0.0042 |
| *Rps24* | Proteins of Human Pre-mRNA Cleavage Complex | 0.5112 | 0.0078 |
| *Rpl8* | Proteins of Human Pre-mRNA Cleavage Complex | 0.5429 | 0.0030 |
| *Rpl9* | Proteins of Human Pre-mRNA Cleavage Complex | 0.7977 | 0.0028 |
| *Rpl24* | Proteins of Human Pre-mRNA Cleavage Complex | 0.6713 | 0.0011 |
| *Rpl27* | Proteins of Human Pre-mRNA Cleavage Complex | 0.4135 | 0.0085 |
| *Lmna* | Proteins of Human Pre-mRNA Cleavage Complex | 0.9294 | 0.0004 |
| *Farsb* | Proteins of Human Pre-mRNA Cleavage Complex | 0.4109 | 0.0268 |
| *Cstf2* | Known PolyA Factors of CPSF Complex | 0.3667 | 0.0333 |
| *Cstf3* | Known PolyA Factors of CPSF Complex | 0.4457 | 0.0164 |
| *Fip1l1* | Known PolyA Factors of CPSF Complex | 0.5514 | 0.0076 |
| *Papolg* | Other Known PolyA Factors | 0.2874 | 0.0478 |
| *Pabpc1* | Other Known PolyA Factors | 0.6314 | 0.0082 |
| *Rbm7* | Factors with Known Motifs | 0.3734 | 0.0043 |
| *Ddx3x* | Factors with Known Motifs | 0.2997 | 0.0233 |
| *Dhx15* | Factors with Known Motifs | 0.3310 | 0.0133 |
| *Ddx20* | Factors with Known Motifs | 0.3270 | 0.0494 |
| *Ddx23* | Factors with Known Motifs | 0.4901 | 0.0063 |
| *Snd1* | Factors with Known Motifs | 0.7009 | 0.0075 |
| | Factors with Known Motifs | | |

Supplementary Table 14: Dysregulated transcription and translation machinery factors in PAN and ADR models

**PAN**

Core Sequence Elements and Factors

| Gene | logFC PANvsCtl | adj.P.Val PANvsCtl |
|---|---|---|
| *Cpsf1* | 0.3674 | 0.03 |
| *Cstf1* | 0.307 | 0.03 |
| *Cstf2* | 0.471 | 0.01 |
| *Cstf3* | 0.38 | 0.04 |

Proteins of Human Pre-mRNA Cleavage Complex

| | | |
|---|---|---|
| *Cpsf1* | 0.37 | 0.03 |
| *Cstf2* | 0.47 | 0.01 |
| *Cstf3* | 0.38 | 0.04 |
| *Ddx49* | 0.54 | 0.00 |
| *Dhx8* | 0.30 | 0.04 |
| *Farsb* | 0.40 | 0.04 |
| *Hnrnpa3* | 0.31 | 0.04 |
| *Lmna* | 0.66 | 0.01 |
| *Mdc1* | 0.67 | 0.01 |
| *Nelfb* | 0.41 | 0.02 |
| *Nelfe* | 0.57 | 0.01 |
| *Nono* | 0.35 | 0.02 |
| *Nudt21* | 0.40 | 0.01 |
| *Pabpc1* | 0.70 | 0.01 |
| *Parp1* | 0.41 | 0.02 |
| *Rpl24* | 0.48 | 0.01 |
| *Rpl27* | 0.31 | 0.05 |
| *Rpl8* | 0.42 | 0.02 |
| *Rps24* | 0.44 | 0.03 |
| *Rps5* | 0.34 | 0.04 |
| *Scaf1* | 0.35 | 0.05 |
| *Smarca4* | 0.47 | 0.03 |
| *Smarcd2* | 0.32 | 0.03 |
| *Thoc6* | 0.72 | 0.01 |
| *Top2a* | 2.63 | 0.00 |
| *Usp39* | 0.36 | 0.02 |
| *Xrcc6* | 0.74 | 0.00 |

Known PolyA Factors of CPSF Complex

| | | |
|---|---|---|
| *Cpsf1* | 0.37 | 0.03 |
| *Cstf1* | 0.31 | 0.03 |
| *Cstf2* | 0.47 | 0.01 |
| *Cstf3* | 0.38 | 0.04 |

Other Known PolyA Factors

| | | |
|---|---|---|
| *Pabpc1* | 0.70 | 0.01 |
| *Pabpc4* | 0.49 | 0.04 |

Factors with Known Motifs

| | | |
|---|---|---|
| *Ddx23* | 0.37 | 0.03 |
| *Snd1* | 0.68 | 0.01 |

DNA Damage Response Factors

| | | |
|---|---|---|
| *Xrcc6* | 0.74 | 0.00 |
| *Parp1* | 0.41 | 0.02 |

RNAP II and associated factors

| | | |
|---|---|---|
| *Polr2b* | 0.31 | 0.05 |
| *Polr2j* | 0.44 | 0.01 |

Integrator Complex

**ADR**

Core Sequence Elements and Factors

| Gene | logFC ADRvsCtl | adj.P.Val ADRvsCtl |
|---|---|---|
| *Cstf2* | 0.3667 | 0.0333 |
| *Cstf3* | 0.4457 | 0.0164 |
| *Fip1l1* | 0.5514 | 0.0076 |

Proteins of Human Pre-mRNA Cleavage Complex

| | | |
|---|---|---|
| *Cstf2* | 0.3667 | 0.0333 |
| *Cstf3* | 0.4457 | 0.0164 |
| *Nudt21* | 0.4128 | 0.0027 |
| *Fip1l1* | 0.5514 | 0.0076 |
| *Pabpc1* | 0.6314 | 0.0082 |
| *Ppp1cb* | 0.2828 | 0.0220 |
| *Top2a* | 1.9162 | 0.0056 |
| *Parp1* | 0.3186 | 0.0433 |
| *Prkdc* | 0.3892 | 0.0457 |
| *Xrcc6* | 0.6845 | 0.0021 |
| *Thoc6* | 0.6065 | 0.0218 |
| *Nelfb* | 0.4534 | 0.0054 |
| *Nelfe* | 0.6760 | 0.0030 |
| *Smarca4* | 0.4665 | 0.0212 |
| *Smarcd2* | 0.3123 | 0.0292 |
| *Erh* | 0.3311 | 0.0393 |
| *Rnf20* | 0.4453 | 0.0097 |
| *Usp39* | 0.4715 | 0.0026 |
| *Sf3b1* | 0.5080 | 0.0077 |
| *Nono* | 0.3700 | 0.0112 |
| *Srsf1* | 0.3335 | 0.0304 |
| *Srsf10* | 0.4026 | 0.0338 |
| *Hnrnpa3* | 0.4734 | 0.0039 |
| *Ddx49* | 0.6733 | 0.0008 |
| *Dhx8* | 0.3119 | 0.0268 |
| *Dhx15* | 0.3310 | 0.0133 |
| *Exosc4* | 0.3637 | 0.0229 |
| *Exosc7* | 0.3230 | 0.0375 |
| *Exosc8* | 0.5351 | 0.0201 |
| *Rps5* | 0.4909 | 0.0042 |
| *Rps24* | 0.5112 | 0.0078 |
| *Rpl8* | 0.5429 | 0.0030 |
| *Rpl9* | 0.7977 | 0.0028 |
| *Rpl24* | 0.6713 | 0.0011 |
| *Rpl27* | 0.4135 | 0.0085 |
| *Lmna* | 0.9294 | 0.0004 |
| *Farsb* | 0.4109 | 0.0268 |

Known PolyA Factors of CPSF Complex

| | | |
|---|---|---|
| *Cstf2* | 0.3667 | 0.0333 |
| *Cstf3* | 0.4457 | 0.0164 |
| *Fip1l1* | 0.5514 | 0.0076 |

Other Known PolyA Factors

| | | |
|---|---|---|
| *Papolg* | 0.2874 | 0.0478 |
| *Pabpc1* | 0.6314 | 0.0082 |

Factors with Known Motifs

| | | |
|---|---|---|
| *Rbm7* | 0.3734 | 0.0043 |
| *Ddx3x* | 0.2997 | 0.0233 |
| *Dhx15* | 0.3310 | 0.0133 |
| *Ddx20* | 0.3270 | 0.0494 |
| *Ddx23* | 0.4901 | 0.0063 |
| *Snd1* | 0.7009 | 0.0075 |

DNA Damage Response Factors

| | | |
|---|---|---|
| *Prkdc* | 0.3892 | 0.0457 |
| *Xrcc6* | 0.6845 | 0.0021 |
| *Parp1* | 0.3186 | 0.0433 |

RNAP II and associated factors

| | | |
|---|---|---|
| *Polr2b* | 0.3471 | 0.0209 |
| *Polr2e* | 0.3193 | 0.0416 |
| *Polr2j* | 0.3519 | 0.0213 |

Integrator Complex

| | | |
|---|---|---|
| *Ints2* | 0.50 | 0.01 |
| *Ints3* | 0.39 | 0.03 |
| *Ints10* | 0.38 | 0.02 |

| | | |
|---|---|---|
| *Ints2* | 0.3699 | 0.0312 |
| *Ints3* | 0.4074 | 0.0156 |
| *Ints4* | 0.3873 | 0.0176 |
| *Ints6* | 0.3595 | 0.0118 |
| *Ints10* | 0.3987 | 0.0123 |

FACT Complex

| | | |
|---|---|---|
| *SUPT16H* | 0.29 | 0.04 |

Exosome

| | | |
|---|---|---|
| *MTREX* | 0.3793 | 0.0499 |

Splicing Factors

| | | |
|---|---|---|
| *Sf3a1* | 0.268 | 0.05 |
| *Sf3a2* | 0.390 | 0.02 |
| *Sf3a3* | 0.389 | 0.03 |
| *Sf3b2* | 0.297 | 0.05 |
| *Sf3b4* | 0.411 | 0.02 |
| *U2af1* | 0.391 | 0.04 |
| *Prpf19* | 0.682 | 0.01 |
| *Nono* | 0.352 | 0.02 |
| *Sfpq* | 0.398 | 0.02 |
| *Ddx39b* | 0.484 | 0.04 |

Splicing Factors

| | | |
|---|---|---|
| *Sf3a3* | 0.4134 | 0.0147 |
| *Sf3b1* | 0.5080 | 0.0077 |
| *Sf3b2* | 0.4549 | 0.0043 |
| *U2af1* | 0.5091 | 0.0070 |
| *Prpf19* | 0.5438 | 0.0116 |
| *Prpf38b* | 0.3613 | 0.0340 |
| *Nono* | 0.3700 | 0.0112 |
| *Sfpq* | 0.3650 | 0.0165 |
| *Puf60* | 0.3118 | 0.0331 |

Translation Factors

| | | |
|---|---|---|
| *Eef1g* | 0.45 | 0.01 |
| *Eif2a* | 0.30 | 0.04 |
| *Eif4a1* | 0.39 | 0.03 |
| *Eif3f* | 0.39 | 0.03 |
| *Eif3i* | 0.42 | 0.01 |
| *Rack1* | 0.38 | 0.03 |

Translation Factors

| | | |
|---|---|---|
| *Eef1g* | 0.5736 | 0.0017 |
| *Eif2a* | 0.3767 | 0.0090 |
| *Eif4a1* | 0.4423 | 0.0115 |
| *Eif3a* | 0.4041 | 0.0314 |
| *Eif3f* | 0.4617 | 0.0092 |
| *Eif3i* | 0.4300 | 0.0077 |
| *Rack1* | 0.4605 | 0.0071 |
